# Capturing human expertise for AI-assisted molecular structure annotation

**DOI:** 10.64898/2026.11.25.769360

**Authors:** Wilhan D. Gonçalves Nunes, Janele Ting, Abubaker Patan, Julius Agongo, Crystal X. Wang, Harsha Gouda, Victoria Deleray, Md Sakil Arman, Sooraj Baijnath, Ipsita Mohanty, Stephen Barnes, Lee R. Hagey, Mingxun Wang, Helena Mannochio-Russo, Pieter C. Dorrestein

**Author notes:** **Correspondence:** Helena Mannochio-Russo, Pieter C. Dorrestein. These authors contributed equally to this work (Wilhan D. Gonçalves Nunes, Janele Ting).

## Abstract

We developed a large language model-based tool that captures expert reasoning in bile acid structure elucidation, drawing on handwritten notebooks, scientific literature, and recorded voice-to-text analytical sessions. It generates human-readable reports detailing evidence for and against candidate structures, tracing each inference to its source and identifying ambiguities. When applied to data, the tool proposes bile acid conjugates lacking reference spectra.

## Main

Untargeted metabolomics generates far more MS/MS spectra than can be assigned to structures. Most spectra have no match to reference libraries, particularly for molecules arising from microbial metabolism, drug transformation, or diet. *In silico* fragmentation and machine learning can narrow potential candidate structures^1–4^, while molecular networking can propagate annotations across related spectra^5^. Both approaches scale annotation, but neither performs the final step for a genuinely novel spectrum – integrating precursor mass, co-eluting ions, molecular network neighbors, fragments, neutral losses, chemical and biochemical knowledge, structural plausibility, and the behavior of molecules in the mass spectrometer – into a specific molecular proposal, while stating what the data do and do not establish. This important step remains largely manual, and such capability is concentrated in a very small number of experienced researchers. Even a correct structural proposal is of limited value to other researchers if they cannot understand how it was reached, evaluate the supporting evidence, or trace the underlying knowledge to its original sources.

Researchers specialized in structure elucidation based on LC-MS/MS do not interpret fragment ions in isolation. Much of this pattern recognition is done in their heads, based on decades of experience reviewing spectral data. They recognize similarities to related structures while considering mass accuracy, diagnostic ions, ion ratios, neutral-loss patterns, and retention time behavior. They also account for alternative ion forms, adducts, in-source fragmentation, and chimeric spectra arising from precursor co-isolation, which can introduce misleading structural evidence. They also recognize when a seemingly useful rule does or does not apply, when an expected ion is present or absent, and when the available evidence is insufficient to make a structural proposal. Much of this knowledge comes from years to decades of interpreting spectra, cross-referencing biochemical and chemical knowledge and is rarely captured explicitly in a form that can be readily reused computationally.

Bile acids provide a useful test case because their chemistry is well characterized but combinatorial. Their chemical diversity arises from two sources: changes to the steroid core itself, such as hydroxylation, oxidation and epimerization, and changes to groups attached to the core, either at the side-chain carboxyl or on the ring hydroxyls^6^. Here we refer to any such attached group as a “cargo”, whatever its attachment site, bond type, or biological origin. This is not standard nomenclature, and is the word used by the bile acid experts whose knowledge is captured in this work, who view bile acids as carriers of other molecules. Variations in hydroxylation, oxidation, and conjugation to amino acids, amines, and other cargos produces many closely related structures^6^, further expanded by the discovery that the gut microbiome conjugates bile acids to amines and amino acids beyond taurine and glycine^7,8^, alkamine formation^9^, and reactions that form carbon-carbon bonds^10^. Recent large-scale efforts have detected thousands of candidate bile acid modifications in public metabolomics data and validated a subset with synthesized standards^7,8,11^. Because many of these lack reference spectra, many structurally undescribed bile acids likely remain to be characterized. Distinguishing these molecules by MS/MS requires recognizing diagnostic ions, ion ratios, dehydration, neutral-loss, and other fragmentation patterns that can differ subtly between related structures. Although individual rules have been at least partially described^11–13^, deciding which rule applies to a spectrum, whether the required evidence is present, and when a rule should not be generalized remains an expert task.

We captured expertise from sources that had never previously been combined: (1) a career reference of 1,683 pages documenting bile acid diversity and analysis, biological context, and cross-species analysis and interpretation; query-language-based MS/MS interpretation; (2) primary literature and spectral libraries; (3) scanned 4,473 pages of personal handwritten notebooks assembled over decades and containing diagnostic fragment ions and structural observations; and (4) voice transcripts of recorded structure-elucidation sessions in which experts, sometimes as many as six at a time, worked through individual spectra, while annotating printed spectra and MS traces while explaining their reasoning out loud or in group discussion (**Figure 1a,b**). Together, these sources represented many decades of combined experience interpreting bile acid mass spectra. The recorded sessions were particularly valuable for capturing not only their conclusions, but also what they examined, the order in which they examined it, and why particular observations supported or excluded specific structural hypotheses. The recordings that we discuss below also carried the experts’ own vocabulary into the large language model (LLM) skill, which is why its reports call any group attached to a bile acid the cargo. These reasoning sessions were transcribed and converted into source-linked records connecting each inference to its underlying handwritten notebook entry, published rule, spectrum, or structural precedent.

**Figure 1.**
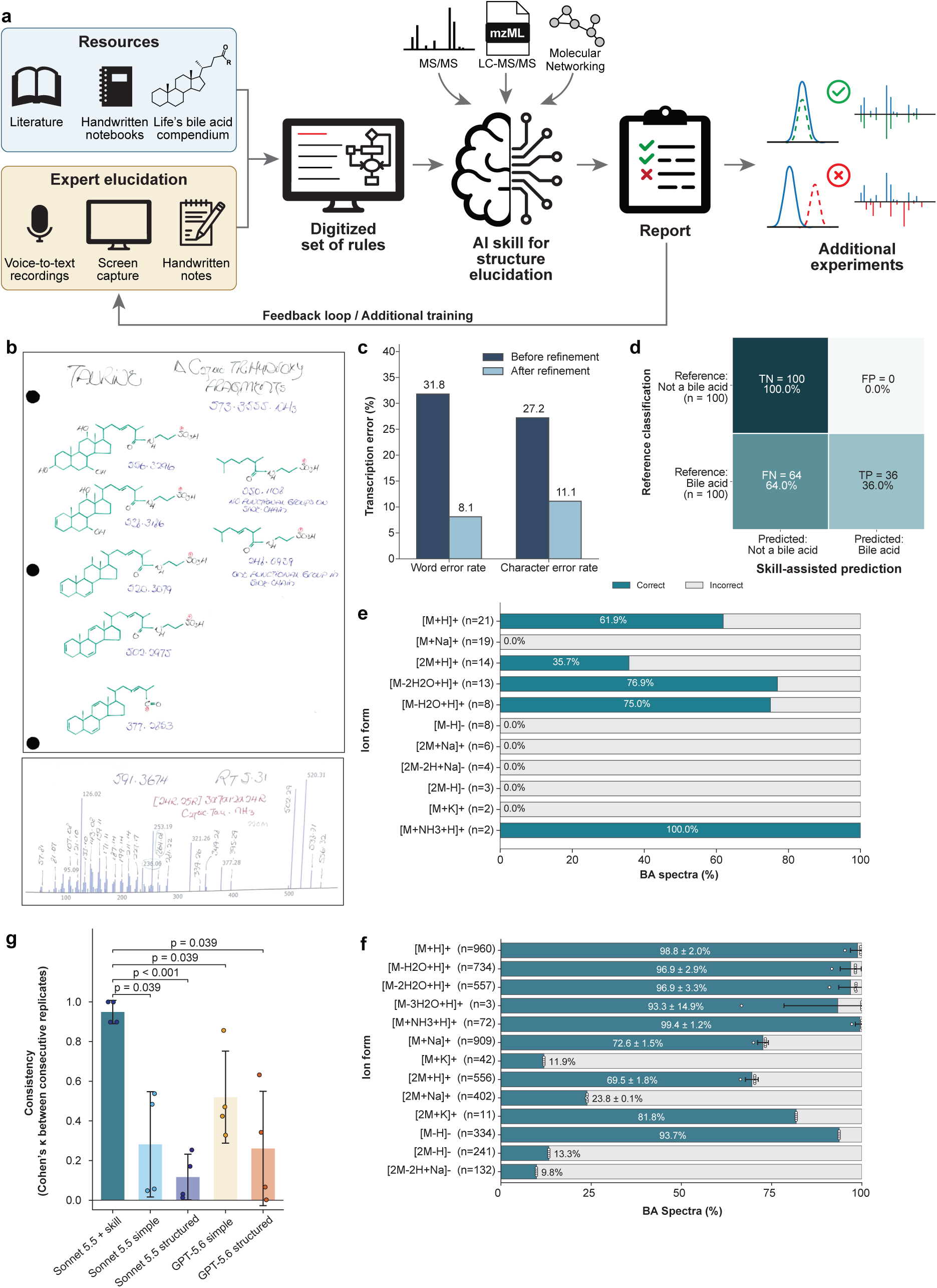
Digitization of expert knowledge for AI-assisted bile acid MS/MS annotation and performance evaluation. **(a)** Overview of the workflow for capturing and digitizing expert knowledge from scientific literature, handwritten notebooks, a comprehensive bile acid reference compendium, and recorded expert structure-elucidation sessions. The resulting knowledge is organized into a reusable AI skill that integrates MS/MS spectra, LC-MS/MS data, and molecular networking information to generate evidence-based structural annotation reports. Reports document supporting and conflicting evidence and guide additional experiments. **(b)** Representative pages from expert handwritten notebooks containing bile acid structures, diagnostic fragment ions, and annotated MS/MS spectra. **(c)** Word error rate (WER) and character error rate (CER) before and after refinement of handwriting transcription using examples of the expert’s handwriting and common transcription errors. **(d)** Confusion matrix showing the initial evaluation of skill-assisted bile acid recognition using 200 reference MS/MS spectra, comprising 100 bile acid and 100 non-bile-acid spectra. Cells indicate the number and percentage of true negatives (TN), false positives (FP), false negatives (FN), and true positives (TP). **(e)** Bile acid recognition among the 100 reference bile acid spectra from the initial evaluation, stratified by precursor ion form. **(f)** Bile acid recognition across 4,953 reference MS/MS spectra from the GNPS BILELIB19 library^7^, stratified by 13 precursor ion forms, over 5 replicates. Dots represent each replicate, error bars the standard deviation. **(g)** Comparison of bile acid recognition using Claude Sonnet 5.5 with the expert-derived skill and general-purpose language models without the skill (Claude Sonnet 5.5 and GPT-5.6), evaluated using simplified and detailed prompts. Cohen’s kappa between the same consecutive replicate pairs (higher = more consistent). Bars: mean; error bars: SD; dots: the four consecutive replicates (1-2, 2-3, 3-4, 4-5). Brackets: Welch’s t-test against Sonnet 5.5 + skill, Holm-adjusted over the 4 comparisons.

The handwritten notebooks required a dedicated approach because the model frequently misinterpreted individual characters, such as confusing the digits 8 and 2 or the letters D and O. Initial transcription resulted in a word error rate (WER) of 31.8% and a character error rate (CER) of 27.2%. We therefore provided the model with examples of the expert’s handwritten alphabet and common transcription errors to help it interpret the handwriting more reliably. Following this refinement, WER decreased to 8.1% and CER to 11.1% (**Figure 1c**). We then integrated the transcribed notebooks, recorded reasoning sessions, literature, and reference spectra into a modular knowledge base, linking individual inferences to their underlying sources. The resulting rule system encompasses bile acid core recognition, carbon number, hydroxylation, conjugation, cargo annotation, and physicochemical parameter estimation (LogP and CCS). Importantly, it also captures exceptions and conditions under which structural assignments should not be made. This knowledge was organized into a reusable LLM skill that can be applied during MS/MS spectral interpretation, enabling expert reasoning previously scattered across decades of personal notes, scientific literature, reference spectra, and tacit knowledge to be systematically applied to new measurements (**Figure 1a**).

The skill accepts different levels of experimental input, ranging from individual MS/MS spectra to complete LC-MS/MS files (mzML) accompanied by the scan number of interest. One of the biggest challenges when providing only the MS/MS and no other context is the identification of the ion form. The more data and input information provided, the more effective the skill. Although it can handle only MS/MS input, providing at least the complete LC-MS/MS run is preferable because it allows the skill’s tools to examine the chromatographic and MS1 context surrounding the selected MS/MS spectrum, improving the assessment of precursor ion forms, including adducts, multimers, and in-source fragments. This distinction is particularly important for ammonium adducts, which can otherwise lead to the incorrect inference of nitrogen-containing molecular structures when interpreted from MS/MS alone. The skill can also integrate data acquired in positive and negative ionization modes under matched chromatographic conditions, using retention time and precursor relationships to identify corresponding features across runs. These complementary measurements provide additional structural evidence, as positive-ion fragmentation is often particularly informative for the bile acid core characterization, whereas negative-ion spectra can help annotate conjugated cargos. Molecular networking information can also be incorporated, either through uploaded networking results or by retrieving outputs from GNPS2 Everything Bagel workflow^14^ through the GNPS Model Context Protocol (MCP). Network connectivity, related spectral annotations, and mass differences between connected nodes are then evaluated by the skill to provide additional context for structural hypotheses.

For each analyzed feature, it generates a structured annotation report documenting the proposed molecular classification, the evidence supporting or contradicting each hypothesis, and the limitations of the available data. Importantly, this reporting framework makes the reasoning process auditable and allows researchers to identify missing fragmentation rules, inconsistencies, and unresolved structural questions. These observations can guide further expert knowledge capture and refinement of the skill, establishing an iterative framework in which new evidence and accumulated expertise can progressively expand the knowledge base. The report is designed to be read and checked by researchers who are not structure elucidation specialists: each call is graded from confirmed to unresolved, each inference carries a source tag that leads back to the notebook entry, published rule, spectrum, or transcript behind it, and what the data cannot resolve is listed explicitly (Section 2.5). The skill can also be used interactively through the LLM’s chat interface, allowing users, whether experienced or new to bile acid analysis, to ask follow-up questions about any part of a report or hypothesis. As in the report, responses draw on the underlying corpus of information and cite the specific sources for each claim, so that users can check the evidence themselves. The report is therefore intended as a starting point for further investigation rather than a final answer.

To evaluate whether the expert-derived knowledge could support bile acid recognition, we first tested 200 randomly selected reference MS/MS spectra, including 100 bile acids represented by different ion forms and 100 non-bile acid compounds. Of the 100 reference bile acid spectra, 36 were correctly recognized as bile acids, while none of the 100 non-bile acid spectra were incorrectly classified as such (**Figure 1d**). This corresponds to a precision of 1.00 (95% CI 0.90– 1.00), a specificity of 1.00 (95% CI 0.96–1.00), a recall of 0.36 (95% CI 0.27–0.46) and an overall accuracy of 0.68. The F1 score was 0.53. F1, however, weights false positives and false negatives equally, an assumption that does not fit the intended use of the skill, which was designed as a conservative annotator, for which an incorrect bile acid assignment costs more than a missed one. When diagnostic evidence is insufficient, it withholds a classification rather than guessing. The precision-weighted F0.5, which better reflects this design, was 0.74; the recall-weighted F2 was 0.41. However, recognition among bile acid spectra varied substantially depending on the precursor ion form (**Figure 1e**), which led us to perform a systematic evaluation using a larger reference bile acid spectral library. Therefore, we expanded the analysis to 4,953 bile acid MS/MS spectra that are part of the BILELIB19 library in GNPS^7^, which contains MS/MS spectra encompassing 13 different ion forms in positive and negative ionization modes. Recall was highest for protonated and ammoniated monomers in positive ionization mode, including [M+H]+, [M−nH_2_O+H]^+^ (n = 1–3) and [M+NH_4_]^+^, ranging from 93.3 ± 14.9% to 99.4 ± 1.2% (**Figure 1f**). In contrast, performance decreased for certain adducts and multimers, including [M+K]⁺ (11.9%), [2M+Na]⁺ (23.8 ± 0.1%), and [2M−H]^−^ (13.3%). These differences reflect the greater representation of fragmentation rules for protonated ions, their dehydrated forms, and ammonium adducts in the captured expert knowledge, whereas other ion forms may exhibit distinct fragmentation behavior not yet fully represented in the knowledge base. As these are among the most commonly detected ion forms of bile acids in positive ionization mode, many bile acids in typical LC-MS/MS datasets are expected to be represented by at least one ion form that the skill recognizes more reliably.

Beyond recognition, the skill also enabled the reassessment of spectral library-based annotations through independent interpretation of experimental LC-MS/MS data. For example, when analyzing a feature whose MS/MS spectrum matched a valine-conjugated bile acid in the GNPS spectral library, the skill instead favored a 5-aminovaleric acid conjugate. Valine and 5-aminovaleric acid are isomers (C_5_H_11_NO_2_), with a branched and a linear structure, respectively. The presence of a dominant fragment ion at *m/z* 100.0758, consistent with lactam formation, together with the absence of the characteristic valine immonium ion at *m/z* 72.0808, supported the alternative annotation (**Supplementary Figure 1, Supplementary File 1**). This example illustrates how expert-derived rules can complement spectral library matching by identifying potentially incorrect annotations and proposing alternative structures based on fragmentation evidence.

We further compared the consistency of bile acid recognition using the skill-assisted approach with interpretations generated by general-purpose language models without access to the skill. We evaluated Sonnet 5.5 (Anthropic) and GPT-5.6 (OpenAI) using simplified and detailed prompts to determine whether general prompting could achieve comparable reproducibility. The skill-assisted approach was the most reproducible, with a mean Cohen’s κ between consecutive replicates of 0.949 ± 0.059, compared with 0.117 ± 0.115 (structured) and 0.282 ± 0.266 (simple) for Sonnet 5.5 without the skill, and 0.261 ± 0.288 (structured) and 0.520 ± 0.232 (simple) for GPT-5.6 across the prompts tested (**Figure 1g**). Consistent with these results, per-spectrum analysis of the 4,953 test spectra showed that the skill-assisted approach had fewer spectra with changing calls across the five replicates than any condition without the skill (**Supplementary Figure 2**). Because Claude was used throughout skill development, model-specific optimization may have influenced the relative performance of the language models. We also observed improved recognition by Claude without skill access in later evaluations, highlighting potential temporal variability in baseline performance especially as the skill is being more widely used. Despite these considerations, the improved recognition shows that expert knowledge accumulated over decades of spectral interpretation can be translated into reusable computational rules that go beyond general-purpose prompting. Recognition is not the key main point, however. The skill also generates a human-readable report for every spectrum that lays out the evidence for and against each candidate structure, traces each inference to its source, and flags what remains ambiguous, so that a reader without expert training can inspect how a call was reached and judge how far to trust it.

To evaluate whether the skill could facilitate the structural annotation of previously unannotated bile acids in biological samples, we investigated fecal metabolomics data from a cohort of individuals with and without HIV recruited through the HIV Neurobehavioral Research Center (HNRC)^15^. Experimental MS/MS spectra were searched against recently developed MassQL-based bile acid spectral libraries^11,13^, which capture characteristic fragmentation patterns of different bile acid classes but contain predominantly structurally uncharacterized spectra. From the resulting 1,366 matches, we selected two features for further investigation using the bile acid annotation skill developed in this study as they appeared to be nonstandard bile acid modifications. The first was proposed to be a C24 bile acid conjugated to a C18:1 fatty acid, with a putative oxo group at C-12 (**Supplementary File 2**). The second was assigned to a dihydroxylated C24 bile acid containing an additional moiety with the molecular formula C_9_H_17_NO_2_ (**Supplementary File 3**). When prompted to explore potential identities for this modification, the skill suggested gabapentin, a medication used to treat neuropathic pain and epilepsy, as a plausible candidate. These structural hypotheses guided the synthesis of 12-ketolithocholic acid-oleic acid and deoxycholic acid-gabapentin, respectively. Subsequent LC-MS/MS analysis demonstrated matching retention times and fragmentation spectra between the synthesized standards and the corresponding features detected in the fecal samples, supporting their structural identities (**Figure 2**).

**Figure 2.**
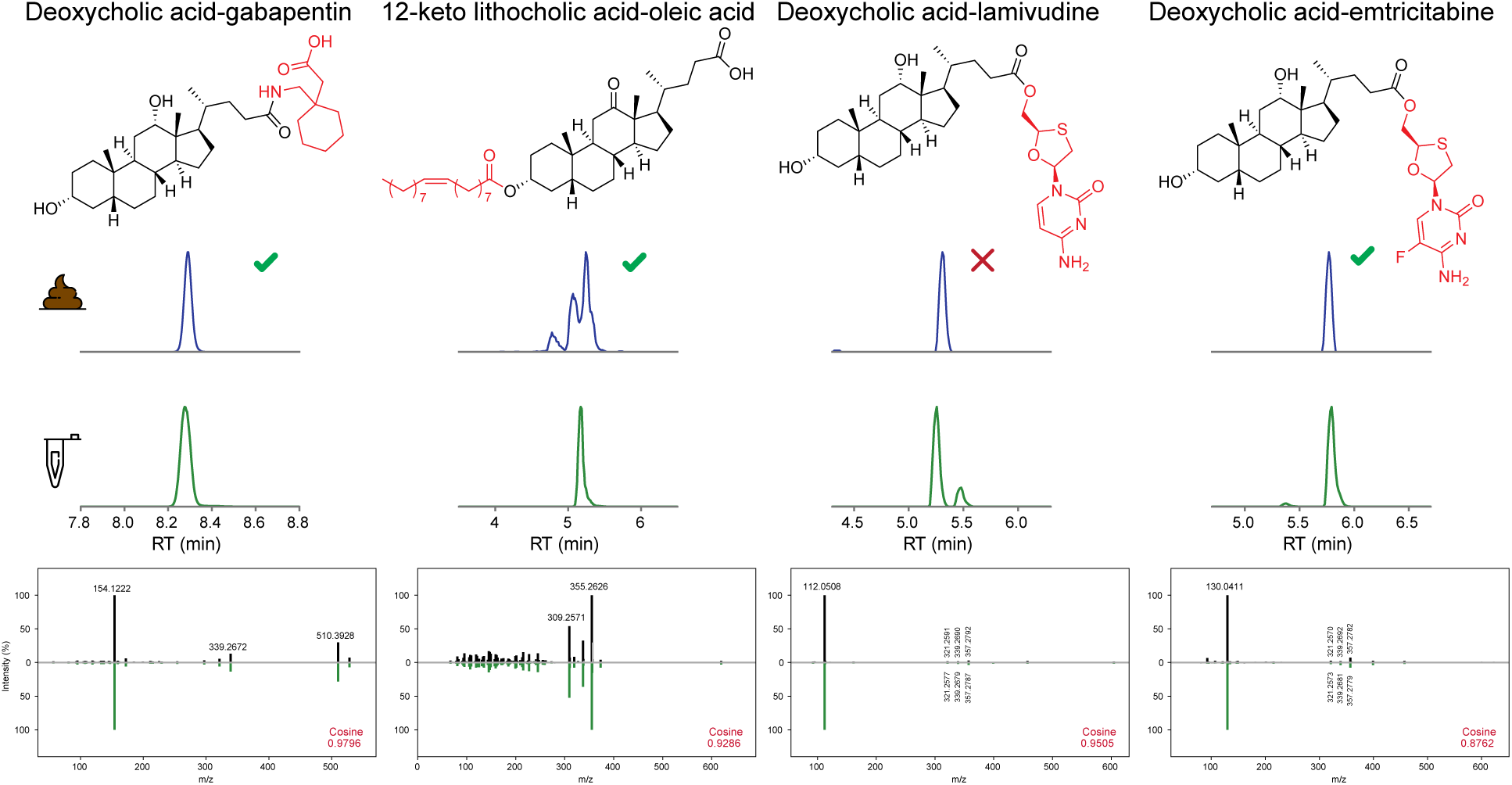
Skill-guided structural annotation and experimental validation of previously uncharacterized bile acid conjugates in human fecal samples. Four bile acid conjugates identified in the HNRC and AWI-Gen 2 cohorts were investigated using the bile acid annotation skill and subsequently evaluated against synthesized reference standards. Top: Chemical structures of the synthesized reference compounds, with the conjugating cargos highlighted in red. Middle: Extracted ion chromatograms of the corresponding features detected in fecal samples (blue) and synthesized reference standards (green). Bottom: Mirror plots comparing experimental MS/MS spectra from fecal samples and synthesized standards. The raw data to this analysis is available in MassIVE (MSV000103990). Green checkmark, at least one of the peaks retention times matches the synthetic standard. Red X none of the retention times match.

We next investigated whether the skill could identify additional drug-conjugated bile acids in a separate metabolomics dataset of samples from the AWI-Gen 2 cohort, which consists of fecal samples collected from women across four African countries^16^. Because we already had evidence for three drug-conjugated bile acids, including gabapentin in the HNRC cohort, 5-aminosalicylic acid (5-ASA) in earlier inflammatory bowel disease and rheumatoid arthritis datasets^17^, and the fluorouracil metabolite α-fluoro-β-alanine in cancer patients^18^, we asked whether other drugs also form bile acid conjugates. Rather than evaluating individual candidate spectra, we prompted the skill to find features that may correspond to bile acids modified by pharmaceutical compounds. Two of the highlighted candidates were selected for detailed interpretation. The first was proposed to be a dihydroxylated C24 bile acid esterified at the C-24 carboxyl group to a nucleoside analogue with the molecular formula C_8_H_11_N_3_O_3_S, consistent with lamivudine (**Supplementary File 4**). The second was proposed to be a dihydroxylated C24 bile acid esterified to the hydroxymethyl group of emtricitabine (**Supplementary File 5**). Notably, lamivudine and emtricitabine are nucleoside reverse transcriptase inhibitors widely used in antiretroviral therapy for HIV. To experimentally evaluate these predictions, we synthesized deoxycholic acid conjugates of both drugs and compared their retention times and MS/MS fragmentation patterns with the corresponding features detected in the biological samples (**Figure 2**). The deoxycholic acid-emtricitabine standard showed matching retention time and MS/MS fragmentation, supporting the proposed structure. In contrast, although the deoxycholic acid-lamivudine standard exhibited similar MS/MS fragmentation with high cosine similarity, a retention time difference was observed relative to the biological feature, suggesting that the detected compound may contain a different bile acid core isomer. Together, these applications demonstrate how the skill can extend beyond recognizing established bile acid fragmentation patterns to proposing specific structural modifications in previously uncharacterized biological molecules. None of the four conjugates had a matching reference spectrum in the available spectral libraries at the time of analysis.

Overall, these results show that structure-elucidation expertise accumulated over decades, much of it never written in a digitized format, can be captured from notebooks, literature and recorded reasoning sessions and deployed as a transparent, auditable component of AI-assisted metabolomics. This approach makes expert reasoning reusable, because each annotation is traceable to the spectral evidence and to the source that justifies it, and the system states what the data cannot resolve. Applied to biological data, it proposed previously undescribed bile acid conjugates, including drug-derived conjugates, three of which matched synthesized standards by retention time and MS/MS, while the fourth matched by MS/MS alone. However, several limitations remain. Our benchmark analyses evaluated bile acid recognition rather than the accuracy of finer structural assignments such as hydroxylation position, stereochemistry or cargo identity, and specificity was assessed on a limited set of non-bile acid spectra. Performance depended strongly on ion form, reflecting the emphasis of the captured knowledge on protonated and dehydrated ions. Finally, outputs depend on the underlying language model, which changes over time, and the approach was developed for a single compound class. However, the framework itself is not specific to bile acids. Source-linked rules, deterministic handling of arithmetic, and reports that make uncertainty explicit can be transferred to other compound classes, although each will require its own effort to capture the relevant expertise.

**Online methods:**

## 2. Methods

### 2.1. Overview and design principles

The skill was developed following the Agent Skill Specification, introduced by Anthropic^19,20^, which contains a SKILL.md file, and the scripts, assets and references folders. The instructions present in the skill file were designed with a three-stage principle in mind, in which the LLM should: (1) extract the evidence from the provided data; (2) compare the findings with the available, curated literature data; and (3) reason over the results to propose an annotation. An explicit rule is that the data should never be fabricated. All proposals and reasoning must first be grounded in the evidence, and only when the evidence is not enough to provide a clear explanation, the LLM should propose, and state this clearly. The source code is available on GitHub (skill implementation: https://github.com/wilhan-nunes/bile-acid-msms-annotation; data extraction: https://github.com/janting21/bile-acid-notebook-decoder). All markdown files referenced in the next sections can be accessed in its respective repository.

### 2.2. Knowledge base construction

The knowledge base combines published fragmentation rules with specialist records and biological context. Published sources include the Mass Spectrometry Query Language (MassQL^21^) bile-acid isomer filtering tree and supplementary query table,^12^ bile-acid modification work,^11^ an electrospray ionization mass-difference table^22^ and suspect-library delta tables^23^. A review of 30 papers was distilled into literature_msms_rules.md. Reference files organize diagnostic core ions, conjugate fragments, carbon count, hydroxylation position, stereochemistry, nomenclature and caveats. A reference book on bile acid evolution and nomenclature^24^ supplies nomenclature and host or diet priors, which do not count as spectral evidence. MinerU^25,26^ version 4.0.6, an open-source tool designed for precise PDF text and content extraction, was used to extract the data from the original PDFs.

For the expert reasoning, we captured the knowledge by scanning binders containing hand-written notes (laboratory notebook) collected over years, which include fragmentation patterns, neutral losses, diagnostic regions, fragment structures, stereoisomer assignment rules, etc. Also, since the rationale the expert uses from MS2 spectra to annotation is not captured in written, we recorded multiple interviews and discussions over the course of 2 months. Audio recordings were captured using a Zoom H4n Pro Handy Recorder (Zoom Corporation, Japan), while screen recordings were obtained using the recording functionality of Zoom Meetings (Zoom Communications, Inc., USA). Audio recordings were subsequently transcribed using OpenAI’s Whisper large-v3-turbo automatic speech recognition model^27^. Since the skill was improved over several iterations, the specialist’s review and feedback on annotation reports was also used. All this material is referenced in the reports, to ensure provenance of the information, through source tags, confidence tiers for the proposal, and skill’s internal files references.

#### LLM handwriting extraction

To extract information from the scanned handwritten notebooks, we developed a specialized handwriting skill using a two-stage approach. First, we evaluated one-shot prompting using Claude on 2 pages sampled from a master index, a handwritten list of IDs and taxonomies for all the samples, and a representative combination of numerical and categorical descriptors used throughout the notebooks. Transcription accuracy was assessed using word error rate (WER) and character error rate (CER), calculated by comparing the generated transcriptions against manually transcribed and human-verified reference text. In the second stage, the skill was refined by providing a reference sample of the expert’s handwritten alphabet together with a curated list of recurrent character-recognition errors, including confusion between the digits 8 and 2, the letters D and O, and the digits 2 and 0. These rules and common errors were incorporated into the skill as a document of specialized rules. The refined approach was subsequently evaluated on 10 additional pages of the master index; transcription performance was compared with the initial approach.

The transcripts, notes, and papers text, together with a core-skeleton structure sheet and a MassQL query table, were consolidated into an agent skill using Claude Code (v2.1.224, Claude Sonnet 5), prompted to summarize them into reusable rules and scripts as structured markdown files with key evidence for MS2 interpretation. Three exploration subagents ran in parallel to survey the transcripts, the papers, and the book with the ancillary materials. Their findings were organized into an entry-point workflow file (SKILL.md) and eight reference files:

1. A citation file that assigns a source tag and a confidence tier to every rule. The tiers are paper-validated, specialist heuristic, and book-derived context.
2. A diagnostic core ion file. It covers confirmation of the bile acid core, hydroxylation count from dehydrated-core ion pairs (e.g., m/z 317/335, 319/337, 321/339, and 323/341 for tetra-through mono-hydroxylated C24 cores), ketone mimicry, and the bile alcohol confound.
3. A conjugate fragment file. It covers taurine, glycine, other amino acids, polyamine, sulfate, and glucuronide cargo ions, plus a mass-parity screen for nitrogen.
4. A hydroxylation position file. It describes the low-intensity, low-m/z region markers and the published MassQL isomer filtering tree ^12^, with its branch-level false discovery rates.
5. A carbon-count table that transcribes the MassQL queries for C23–C29 cores across hydroxylation classes, including intensity-ratio queries for non-C24 homologs.
6. A core structure reference that maps substitutable skeleton positions to fragment-based conclusions.
7. A stereochemistry and caveats file. It describes the limits of α/β assignment by MS/MS, mass-defect validation of fragment ions, and known pitfalls.
8. A nomenclature glossary that includes the book’s shorthand notation, the notebook’s confidence conventions, and taxonomic priors.

Each source is available as a markdown source file for the skill, which ensures provenance of the sources for all information.

### 2.3. System architecture

The skill architecture comprises two complementary layers: a deterministic layer and a reasoning layer. The deterministic layer uses Python scripts to implement rules derived from the source materials, including the evaluation of diagnostic fragment ions, fragment presence and relative intensities, mass differences (delta masses), and other quantitative calculations. This approach minimizes the risk of LLM hallucinations in rule-based and mathematical operations. The reasoning layer leverages the LLM to interpret fragmentation patterns that cannot be fully resolved by predefined rules, including the attribution of fragment ions and the generation of hypotheses to explain MS/MS features not captured by the existing rules. Although conceptually distinct, the two layers are not entirely independent, as certain analytical steps integrate both deterministic calculations and LLM-based reasoning. The final annotation report documents the supporting evidence and reasoning, enabling the individual steps of the annotation process to be traced. Each step is further classified according to its implementation (script-based, LLM-based, or hybrid) in **Supplementary Table 1** (**Supplementary Figure 3**).

The deterministic pre-pass extracts and caches the input, screens ion forms and core neutral losses, applies MassQL queries, counts core and low-mass ions, assigns candidate elemental formulas, and compiles chromatographic and optional GNPS2 context. It prepares the annotated spectrum and fills mechanical report fields. The language model assesses conflicting assignments, interprets literature and specialist rules, compares open-ended cargo hypotheses, and records unresolved features. Structure drawing, mass checks, name cleanup and HTML rendering return to scripts. Measured values and arithmetic remain inspectable while structural choices are explicitly reasoned.

The documented workflow runs in Claude Code or claude.ai with the skill installed. Language-model use is integral to interpretation and is therefore disclosed here; the inference model and platform used for each evaluation run must also be reported with the results. The repository documents installation, dependencies and command-line execution. Runtime is measured from input extraction to report rendering with a recorded start timestamp.

### 2.4. Annotation workflow

The full annotation workflow consists of 9 steps, depicted in **Supplementary Figure 3c**, with a final, standardized interactive html report as the output. All steps are grounded in the spectral evidence compiled from the sources described earlier.

#### Input and ion form

The same path accepts mzML, an MGF entry, or a file or pasted list of m/z and intensities. Input is read once into a context cache. Available precursor, polarity, collision energy, MS1 and chromatographic facts are recorded with provenance. When a peak list lacks a precursor label, the highest-*m/z* peak is used provisionally and that assumption is reported. MS1 and chromatographic checks unavailable from an MGF or peak list are marked unavailable. Optional GNPS2 network, feature and library files provide context.

The pre-pass first tests plausible protonated, dehydrated, ammoniated, sodiated, potassiated, dimeric and negative-mode precursor forms. It retains competing forms when data do not discriminate among them. An adduct-agnostic neutral-loss ladder tests for loss of a dehydrated bile-acid core. The C24 ladder contains 358.2872, 374.2821, 390.2770 and 406.2719 Da losses for one through four hydroxyls; a rung requires a fragment at least 2% of the base peak within 0.01 Da. Positive-mode pair and ring-ion rules are not applied to negative-mode spectra; separate fragmentation studies inform the negative-mode checks.^28,29^

#### Bile acid evidence

For positive-mode spectra, the pre-pass checks dehydrated-core pairs such as m/z 319/337, 321/339, 317/335 and 323/341. It accepts each C24 pair ion at 1% of base-peak intensity, an adaptation of the published 5% floor. A pair detected only below 5% is a relaxed-gate pass: the core and overall verdict are capped at tentative, while later attributes are graded on their own evidence. The tree’s published error rates were measured at the original floor.^12^. They are not assigned to the adapted gate. A separate “dust” (core ring, low intensity fragments) test counts at least seven odd-mass, ring-like ions from m/z 100–250 at or above 1% of base peak, after isotope peaks and cargo-like ions are set aside. The count and floor are working defaults, not independently validated thresholds. Pair plus dust supports a bile-acid call; a pair without dust still supports one but leaves dust-ratio position calls unavailable. Dust without a pair is only bile-acid-like and prompts bile-alcohol and non-bile-acid steroid checks. A core neutral-loss hit can provide a tentative route when neither pair nor dust is present. If all three fail, no bile-acid structure is proposed.

#### Formula and cargo

Formula enumeration covers C, H, N, O and S for relevant fragments and neutral losses, with Na or K added for metal adducts. A plausible CHNO formula with a systematic 0.016–0.021 Da error triggers a sulfur near-miss check; precursor and cargo-retaining fragments, and the ³⁴S isotope when MS1 is available, help locate sulfur. Mass-defect outliers are classified as possible co-eluting signals, cargo ions or unresolved signals according to precursor class. Cargo and mass-difference tables name candidates but do not restrict the search. The workflow tests open-ended residual formulas, including unlisted amino-acid, amine, peptide, sugar, acyl, multi-unit and possible xenobiotic cargos. A specific cargo identity requires at least two independent supporting ions; a formula alone is tentative at most and leaves identity unresolved. Predicted but absent diagnostic ions count against a candidate.

When candidates share a formula, the ammonium-adduct result helps rank whole-molecule structures by their free acidic groups. Absence is informative only if the data could have shown the adduct and the mobile phase contains ammonium. If mobile-phase composition is unstated, ammonium is assumed and the inference remains tentative. An adduct observation does not identify a blocking group by itself. Free-acid and C/H/O-cargo readings that are isobaric remain alternatives until fragment evidence separates them.

#### Core and isomer features

Core carbon count is evaluated from side-chain-intact ions and carbon-count MassQL queries, with homolog-shift checks when an exact query is absent. Hydroxyl and ketone positions are assigned from the reached branch of the published filtering tree, with branch-specific error rates where applicable.^3^ If the tree has no valid path, relative-intensity fragment fingerprints provide a lower-confidence fallback and disagreements are recorded. Stereochemistry is assigned only where an applicable ion-ratio or comparable within-run retention-order observation supports it; retention order corroborates a spectral call and never identifies a structure alone.

#### Context, validation, and structure

With GNPS2 files, a feature join adds quantitative and network context. Analog masses and deltas come from the parsed molecular network graph, while library matches are reconciled only after the independent spectral interpretation. A library name or cosine score does not raise confidence. Before reporting, the workflow rechecks mass errors, proton-count and isotope consistency, source scope and conflicts. The final proposal may be a partial structure with unplaced groups or unresolved stereocentres; RDKit renders it, checks mass and standardizes the displayed name. An additional step normalizes the bile acid name using a PubChem lookup. No structure is drawn when the evidence does not support a bile-acid class.

### 2.5. Report generation and guardrails

Each run produces a standardized HTML report with ten sections, or eleven when a molecular-network graph is supplied. Section 1 records input provenance: precursor m/z, ion form, retention time, polarity and collision energy, together with any available instrument and isolation-window metadata. Measured values are distinguished from assumptions. Section 2 shows the annotated MS/MS spectrum in a full-range view and an expanded low-mass view, with diagnostic ions labeled and colored by interpretive role. Section 3 gives the proposed annotation and separate calls for core identity, hydroxylation, conjugation and stereochemistry. Each call is graded as confirmed, probable, tentative or unresolved.

Section 4 compiles confirmatory and exclusionary evidence: observed and calculated masses, elemental-formula assignments, mass errors, relative intensities, diagnostic ratios and neutral losses, with source citations and relevant specialist quotations. It also groups cargo alternatives by composition, directly attached residue, linkage and unresolved unit identity. For each candidate, it records the evidence for and against, its status and its confidence tier. Validation and parity checks close the section. Section 5 records the outcome of the automated classification tree. When no terminal assignment is reached, it explains why classification stalled. Section 6 lists unresolved features, the reason each remains unresolved and the observations that could distinguish the remaining alternatives.

Section 7 gives species and taxonomic context through the host prior, expected producer and plausible dietary route, and suggests follow-up checks. Species and diet are labeled as contextual priors, never as spectral evidence. Section 8 summarizes available chromatographic context, including extracted-ion chromatograms and associated peak information, or states explicitly that these data are unavailable. Section 9 presents the diagnostic decision tree, which links core and conjugate assignments, hydroxylation-position calls and excluded alternatives to the underlying evidence. Every inferential row carries a source tag and a stable evidence identifier, and the tree refers back to those identifiers. A shared color scheme links the spectrum, the evidence tables and the tree.

Section 10 presents proposed structures with standardized names, molecular formulas, calculated masses, confidence tiers and, for complete structures, InChIKeys. The drawings explicitly distinguish unresolved stereochemistry, unplaced substituents and tentative structural units. Accompanying notes state the unresolved features and assumptions. When a molecular-network graph is supplied, Section 11 reports network analogs, mass differences, spectral similarities and available library annotations. These are framed as hypotheses about relatedness, not as structural evidence. Each report records its start and render times, which give its runtime.

### 2.6. Iterative development with expert feedback

The changelog, available in the github repository, records how reports review altered rules during the skill development, including affected files and dates. These are versioned rule changes, not independent validation.

### 2.7. Evaluation design

The skill was first tested on 200 randomly selected reference MS/MS spectra: 100 from bile acids, covering different ion forms, and 100 from non-bile acid compounds. Each spectrum received a binary call (bile acid or not), which was compared with the reference identity. A bile acid called as such was counted as a true positive (TP), and a bile acid not called as such, including cases where the skill withheld a classification because the evidence was insufficient, as a false negative (FN). A non-bile acid called a bile acid was counted as a false positive (FP), and all other non-bile acids as true negatives (TN). We calculated precision as TP/(TP + FP), recall as TP/(TP + FN), specificity as TN/(TN + FP) and accuracy as (TP + TN)/(TP + TN + FP + FN).Two-sided 95% confidence intervals (CIs) for precision, recall and specificity were calculated using the exact Clopper–Pearson method, which is based on the binomial distribution. For x successes in n trials, the lower and upper limits are the 0.025 quantile of Beta(x, n − x + 1) and the 0.975 quantile of Beta(x + 1, n − x), with the lower limit set to 0 when x = 0 and the upper limit to 1 when x = n. The denominators were the number of positive calls for precision (n = 36), the number of reference bile acids for recall (n = 100) and the number of non-bile acids for specificity (n = 100). This method was chosen because it remains valid for proportions at or near 0 or 1, where normal-approximation intervals collapse or exceed these bounds, although it tends to be conservative. The intervals treat spectra as independent observations. The F-score was calculated as Fβ = (1 + β²) × precision × recall / (β² × precision + recall). We report F1 (β = 1), F0.5 (β = 0.5), which weights precision more heavily, and F2 (β = 2), which weights recall more heavily.. To evaluate the performance of the LLM in bile acid annotation in the expanded number of spectra, with and without the use of the specialized skill, we used the BILELIB19 spectral library, comprising 4,953 bile acid entries acquired in positive and negative ionization modes. The spectral library was anonymized by removing all metadata that could reveal compound identities. The spectra were then randomly reordered and assigned sequential identifiers, with a separate mapping retained to link each anonymized identifier to its original spectrum The anonymized spectral library was provided to Claude Code (Sonnet 5.5) for performance evaluation using the built-in skill-creator plugin. The spectra were divided into batches of 50, and multiple independent subagents were deployed to perform the annotations. In total, 200 subagents were used, including 100 with access to the specialized bile acid annotation skill and 100 without skill access. Each subagent was instructed using the following prompt: “Classify every spectrum in the chunk as bile acid or not. Output verdicts.tsv with columns scan, verdict (Bile acid | Non bile acid), note.” 95% confidence intervals for precision, recall and specificity were calculated using the exact Clopper–Pearson method.

### 2.8. Comparison with general-purpose language models

To evaluate whether general-purpose language models could recognize bile acid MS/MS spectra without access to the specialized skill, the anonymized BILELIB19 spectral library was analyzed using Claude Sonnet 5.5 (Anthropic) and GPT-5.6 (OpenAI). Two prompting strategies were evaluated: a simplified prompt requesting direct classification of each spectrum as a bile acid or non-bile acid, and a detailed prompt providing information about bile acid chemistry, fragmentation behavior, conjugation, regioisomerism, and stereochemistry, together with instructions for systematic spectral interpretation. Both approaches were repeated 5 times each and generated CSV files containing binary (yes/no) classifications associated with individual scan numbers. These predictions were matched to the reference spectra, and recognition performance was calculated as the percentage of bile acid spectra correctly classified. Performance was further stratified by positive and negative ionization modes. All results can be found in the **Supplementary Table 2.** The complete prompts are provided in the **Supplementary Methods**.

### 2.9. Skill-guided annotation of bile acid conjugates in biological samples

To evaluate the application of the bile acid annotation skill to biological datasets, untargeted LC-MS/MS data from two independent human cohorts were investigated. The first dataset comprised fecal samples from individuals with and without HIV enrolled through the HIV Neurobehavioral Research Center (HNRC). LC-MS/MS files were subjected to the Everything Bagel workflow^14^ in GNPS2^30^, and searched against MassQL-based bile acid spectral libraries^11,13^ to identify features containing fragmentation patterns characteristic of bile acid classes. Selected features were subsequently investigated using the bile acid annotation skill. For each feature, the corresponding mzML file and molecular networking information in GraphML format were provided, together with the GNPS2 job identifier and feature or scan number. The skill was prompted to annotate the selected MS/MS spectrum and incorporate molecular networking connectivity information, without specifying candidate molecular structures or conjugating groups. The skill was used to interpret fragmentation evidence and propose candidate structural assignments. Additional follow-up prompts were used when necessary to explore possible identities of unresolved molecular modifications. The approach was further applied to a separate metabolomics dataset of fecal samples from the AWI-Gen 2 cohort, focusing on potential drug-conjugated bile acids. Candidate spectra were selected for detailed interpretation using the skill, which was prompted to evaluate their fragmentation patterns and propose possible bile acid structures and conjugating moieties. The resulting structural hypotheses were used to guide the synthesis of reference compounds.

### 2.10. Synthesis

#### 12-keto lithocholic acid-oleic acid

In a 20 mL scintillation vial, oleic acid (30 mg, 0.11 mmol, 1.2 equiv) was added, followed by 4-dimethylaminopyridine (DMAP, 13 mg, 0.11 mmol, 1.2 equiv). Subsequently, triethylamine (Et₃N, 17 µL, 0.12 mmol, 1.4 equiv) and 2,4,6-trichlorobenzoyl chloride (TCBC, 26 mg, 0.11 mmol, 1.2 equiv) were added sequentially in anhydrous tetrahydrofuran (THF, 2 mL). The reaction mixture was stirred at room temperature for 30 min to facilitate the formation of the corresponding mixed anhydride intermediate. Following activation, 12-keto bile acid (69 mg, 0.18 mmol, 2.0 equiv), previously dissolved in THF, was added to the reaction mixture. The reaction was then stirred at 40 °C overnight. Upon completion, THF was removed by evaporation, and the resulting crude residue was reconstituted in HPLC-grade solvents for direct LC–MS/MS analysis without further purification.

#### Deoxycholic acid-gabapentin

In a 20 mL scintillation vial, deoxycholic acid (10 mg, 0.025 mmol, 1.0 equiv) was dissolved in N,N-dimethylformamide (DMF, 2 mL), followed by the sequential addition of HATU (12 mg, 0.031 mmol, 1.2 equiv) and N,N-diisopropylethylamine (DIPEA, 18 µL, 0.10 mmol, 4.0 equiv). The reaction mixture was stirred at room temperature for 20 min to facilitate activation of the carboxylic acid. Subsequently, gabapentin (4.4 mg, 0.025 mmol, 1.0 equiv) was added, and the reaction mixture was stirred at room temperature overnight to promote amide bond formation. Upon completion, DMF was removed by evaporation, and the resulting crude residue was used for LC– MS/MS analysis without further purification.

#### Deoxycholic acid-emtricitabine

In a 20 mL scintillation vial, deoxycholic acid (30 mg, 0.076 mmol, 1.0 equiv) was dissolved in N,N-dimethylformamide (DMF, 2 mL), followed by the addition of 1-ethyl-3-(3-dimethylaminopropyl)carbodiimide (EDC, 18 mg, 0.092 mmol, 1.2 equiv). The reaction mixture was stirred at room temperature for 20 min to facilitate activation of the carboxylic acid. Subsequently, emtricitabine (19 mg, 0.076 mmol, 1.0 equiv) and 4-dimethylaminopyridine (DMAP, 9.3 mg, 0.076 mmol, 1.0 equiv) were added, and the reaction mixture was stirred at room temperature overnight to promote ester bond formation. Upon completion, DMF was removed by evaporation, and the resulting crude residue was reconstituted in HPLC-grade solvents and directly subjected to LC–MS/MS analysis without further purification.

#### Deoxycholic acid-lamivudine

In a 20 mL scintillation vial, deoxycholic acid (30 mg, 0.076 mmol, 1.0 equiv) was dissolved in N,N-dimethylformamide (DMF, 2 mL), followed by the addition of 1-ethyl-3-(3-dimethylaminopropyl)carbodiimide (EDC, 18 mg, 0.092 mmol, 1.2 equiv). The reaction mixture was stirred at room temperature for 20 min to facilitate activation of the carboxylic acid. Subsequently, lamivudine (18 mg, 0.076 mmol, 1.0 equiv) and 4-dimethylaminopyridine (DMAP, 9.3 mg, 0.076 mmol, 1.0 equiv) were added, and the reaction mixture was stirred at room temperature overnight to promote ester bond formation. Upon completion, DMF was removed by evaporation, and the resulting crude residue was reconstituted in HPLC-grade solvents and directly subjected to LC–MS/MS analysis without further purification.

### 2.11. Retention time and MS/MS matching for level 1 annotation

#### HNRC dataset

To achieve level 1 annotation of bile acid conjugates detected in the HNRC dataset (sample P4_D6_Sample_X3162200, dataset MSV000092833), synthesized reference standards were analyzed by LC-MS/MS, and their retention times and fragmentation spectra were compared with those of the corresponding features detected in biological samples. For the confirmation of 12-ketolithocholic acid-oleic acid, samples were analyzed using a Vanquish UHPLC system coupled to a Q Exactive Orbitrap mass spectrometer (Thermo Fisher Scientific). Chromatographic separation was performed using a Kinetex Polar C18 column (100 × 2.1 mm, 2.6 μm particle size, 100 Å pore size; Phenomenex, Torrance, CA, USA) maintained at 55 °C. The mobile phases consisted of (A) acetonitrile/water (60:40, v/v) and (B) isopropanol/acetonitrile (90:10, v/v), both containing 10 mM ammonium formate and 0.1% formic acid, at a flow rate of 0.4 mL/min, with injection volumes set at 2 µL. The gradient was programmed as follows: 0–1 min, 30% B; 1–7.5 min, 30–99% B; 7.5–9.3 min, 99% B; 9.3–9.5 min, 99–30% B; and 9.5–11 min, 30% B. Mass spectrometric analysis was performed using electrospray ionization (ESI) in positive ionization mode. Full-scan MS1 spectra were acquired over an m/z range of 100–1,500 at a resolution of 70,000 (at m/z 200), with an automatic gain control (AGC) target of 5 × 10⁴, and a maximum injection time of 100 ms. Data-dependent MS/MS spectra were acquired for up to five precursor ions per MS1 scan using higher-energy collisional dissociation (HCD), at a resolution of 17,500 (at m/z 200), an AGC target of 1 × 10⁵, and a maximum injection time of 50 ms. The precursor isolation window was set to 2 m/z, with an isolation offset of 0 m/z, and stepped normalized collision energies of 25, 40, and 60 were employed.

For the confirmation of deoxycholic acid-gabapentin (sample P3_D9_Sample_X3157299, dataset MSV000092833), chromatographic separation was performed using the same Kinetex Polar C18 column maintained at 40 °C. The mobile phases consisted of (A) water and (B) acetonitrile, both containing 0.1% formic acid, at a flow rate of 0.5 mL/min, and the injection volumes were set at 10 µL. The gradient was programmed as follows: 0–0.5 min, 5% B; 0.5–1.1 min, 5–20% B; 1.1–7.5 min, 20–60% B; 7.5–8.5 min, 60–100% B; 8.5–10 min, 100% B; 10–10.1 min, 100–5% B; and 10.1–12 min, 5% B. Mass spectrometric analysis was performed using positive ESI. Full-scan MS1 spectra were acquired over an m/z range of 100–900 at a resolution of 35,000 (at m/z 200), an AGC target of 1 × 10⁶, and a maximum injection time of 100 ms. Targeted MS/MS spectra were acquired by parallel reaction monitoring (PRM) for precursor ions at m/z 530.4204, 546.4153, and 562.4102, at a resolution of 17,500 (at m/z 200), an AGC target of 2 × 10⁵, and a maximum injection time of 100 ms. The precursor isolation window was set to 1 m/z, with an isolation offset of 0 m/z, and stepped normalized collision energies of 25, 40, and 60 were employed.

#### AWI-Gen 2 cohort

For the confirmation of deoxycholic acid–lamivudine and deoxycholic acid–emtricitabine, LC-MS/MS analyses were performed using the AG078_AGINCOURT_6_B6 and SW034_SOWETO_3_B10 samples, respectively. Samples (4 μL) were injected into a Vanquish UHPLC system coupled to a Q Exactive Orbitrap mass spectrometer (Thermo Fisher Scientific). Chromatographic separation was performed at a column temperature of 40 °C, with a flow rate of 0.5 mL/min. The gradient consisted of 0–1 min, 5% B; 1–7.5 min, 5–99% B; 7.5–9.3 min, 99% B; 9.3–9.5 min, 99–5% B; and 9.5–11 min, 5% B. Mass spectrometry analysis was performed using electrospray ionization (ESI) in positive ionization mode. Full-scan MS1 spectra were acquired over an m/z range of 100–1,500 at a resolution of 70,000 (at m/z 200), with an automatic gain control (AGC) target of 5 × 10⁴ and a maximum injection time of 100 ms. Data-dependent MS/MS spectra were acquired for up to five precursor ions per MS1 scan, using a precursor inclusion list, at a resolution of 17,500 (at m/z 200), an AGC target of 1 × 10⁵, and a maximum injection time of 50 ms. The precursor isolation window was set to 2 m/z, with an isolation offset of 0 m/z. Higher-energy collisional dissociation (HCD) was performed using stepped normalized collision energies of 25, 40, and 60.

## Data availability

The BILELIB19 reference library used to benchmark the skill is publicly available through GNPS⁷. All annotation reports generated by the skill, both in the BILELIB19 benchmark and for the four use cases, are available on Zenodo (https://doi.org/10.5281/zenodo.23934605).

The HNRC fecal metabolomics data are available in MassIVE under accession MSV000092833. Feature finding and molecular networking were performed with the GNPS2 Everything Bagel workflow (https://gnps2.org/status?task=78d08e8bc3fe463a950c3976402acb4b). The subsequent search against the MassQL-based bile acid libraries is available at https://gnps2.org/status?task=23f842011b3d4a62b9b6251a1b7c98bb.

For the AWI-Gen 2 cohort, the Everything Bagel analysis is available at https://gnps2.org/status?task=9b80f45bebfc466a8ff172404300b51d.

The LC-MS/MS data used to match retention times and MS/MS spectra between the synthesized standards and the biological samples are available in MassIVE under accession MSV000103990.

MS/MS spectra of the synthesized standards were deposited in the GNPS2 library and are accessible through the following Universal Spectrum Identifiers (USIs):

Deoxycholic acid-gabapentin: mzspec:GNPS:USER-CONTRIBUTED-GNPS2:accession:GNPS2LIB0000000EXFW

12-keto lithocholic acid-oleic acid: mzspec:GNPS:USER-CONTRIBUTED-GNPS2:accession:GNPS2LIB0000000EXFX

Deoxycholic acid-lamivudine: mzspec:GNPS:USER-CONTRIBUTED-GNPS2:accession:GNPS2LIB0000000EXFV

Deoxycholic acid-emtricitabine: mzspec:GNPS:USER-CONTRIBUTED-GNPS2:accession:GNPS2LIB0000000EXFY

## Code availability

The source code of the bile acid MS/MS annotation skill is available at https://github.com/wilhan-nunes/bile-acid-msms-annotation. The skill used to transcribe the handwritten notebooks is available at https://github.com/janting21/bile-acid-notebook-decoder. The scripts used to create the figures in the manuscript are available at https://github.com/helenamrusso/Bile_acid_annotation_skill_manuscript.

## Acknowledgements

P.C.D. acknowledges support from EnvedaGives. H.M-R. acknowledges support from the National Institute of Neurological Disorders and Stroke (NINDS; K99NS147878). C.X.W. acknowledges support from the National Institute on Aging (NIA; K23AG102190). H.G. acknowledges support from the American Heart Association (26POST1559436). We thank the H3Africa Wits-INDEPTH Partnership for Genomics Studies (AWI-Gen) and the Sydney Brenner Institute for Molecular Bioscience at the University of the Witwatersrand, Johannesburg, South Africa, for providing samples from the AWI-Gen 2 Microbiome Study. AWI-Gen is funded by the National Human Genome Research Institute (NHGRI) and the National Institute of Environmental Health Sciences (NIEHS) of the National Institutes of Health (NIH) under award U54HG006938 and by the Department of Science and Innovation, South Africa, under award DST/CON 0056/2014. We also thank Scott Hazelhurst, Ami S. Bhatt, Jane A. Cook, Angela S. Hickey, Ariel Gulasch and Helen Choi.

## Author contributions

W.D.G.N., J.T., H.M-R. and P.C.D. designed, developed and tested the bile acid annotation skill. L.R.H., V.D., S.Barnes, P.C.D., W.D.G.N., H.M-R. and J.T. participated in expert discussions and structure elucidation sessions that informed the development of the skill. C.X.W. and H.G. contributed to conceptualization and provided guidance on data analysis and skill evaluation. A.P. and J.A. synthesized the reference compounds. H.M-R. performed the experimental validation through retention time and MS/MS spectral matching. J.T. and W.D.G.N. conducted the benchmark analyses. S.Baijnath coordinated sample handling for the AWI-Gen 2 cohort. M.S.A. and I.M. developed the nomenclature standardization module incorporated into the skill. M.W. contributed to experimental design and computational infrastructure. P.C.D., H.M-R. and W.D.G.N. wrote the manuscript. All authors reviewed and revised the manuscript.

## AI disclosure

In the Dorrestein Lab, the use of AI, generative AI, and large language models (LLMs), and software that uses these technologies, both free and commercial, is encouraged across all aspects of research, including literature review, coding, data analysis, and text editing. All figures and analyses are original. For transparency and reproducibility, all raw data, derived data tables, and final code used in this research are made accessible and linked with this manuscript.

## References

1. Yang, T. et al. AI-guided discovery and expansion of the human bile acid landscape. Cell Metab. S1550-4131(26)00437–7 (2026).

2. Dührkop, K. et al. SIRIUS 4: a rapid tool for turning tandem mass spectra into metabolite structure information. Nat. Methods 16, 299–302 (2019).

3. Xing, S., Shen, S., Xu, B., Li, X. & Huan, T. BUDDY: molecular formula discovery via bottom-up MS/MS interrogation. Nat. Methods 20, 881–890 (2023).

4. Hong, Y., Li, S., Ye, Y. & Tang, H. FIDDLE: a deep learning method for chemical formulas prediction from tandem mass spectra. Nat. Commun. 16, 11102 (2025).

5. Watrous, J. et al. Mass spectral molecular networking of living microbial colonies. Proc. Natl. Acad. Sci. U. S. A. 109, E1743–52 (2012).

6. Mohanty, I. et al. The changing metabolic landscape of bile acids - keys to metabolism and immune regulation. Nat. Rev. Gastroenterol. Hepatol. 21, 493–516 (2024).

7. Gentry, E. C. et al. Reverse metabolomics for the discovery of chemical structures from humans. Nature 626, 419–426 (2024).

8. Quinn, R. A. et al. Global chemical effects of the microbiome include new bile-acid conjugations. Nature 579, 123–129 (2020).

9. Agongo, J., et al. Alkamines reveal a hidden layer of steroid and drug metabolism. bioRxivorg (2026) doi:10.64898/2026.05.13.724743.

10. Ding, Y. et al. Identification of gut microbial bile acid metabolic enzymes via an AI-assisted pipeline. Cell 188, 6012–6027.e20 (2025).

11. Mohanty, I. et al. The underappreciated diversity of bile acid modifications. Cell 187, 1801–1818.e20 (2024).

12. Mohanty, I. et al. MS/MS mass spectrometry filtering tree for bile acid Regio- and stereoisomer annotation. Anal. Chem. 98, 4571–4584 (2026).

13. Mannochio-Russo, H. et al. The evolving bileome: A pan-repository landscape of expanded bile acid and related steroidal signatures. bioRxiv (2026) doi:10.64898/2026.10.10.758218.

14. Shin, Y., et al. The Everything Bagel Feature Finder: Ultra-fast automated feature finding for untargeted metabolomics. bioRxiv (2026) doi:10.64898/2026.08.17.744735.

15. Mannochio-Russo, H. et al. The microbiome diversifies long- to short-chain fatty acid-derived N-acyl lipids. Cell 188, 4154–4169.e19 (2025).

16. Maghini, D. G. et al. Expanding the human gut microbiome atlas of Africa. Nature 638, 718– 728 (2025).

17. Charron-Lamoureux, V. et al. Pan-repository analysis reveals a drug-activating function of microbial bile acid conjugation. bioRxiv (2026) doi:10.64898/2026.03.03.709330.

18. Sweeny, D. J., Barnes, S., Heggie, G. D. & Diasio, R. B. Metabolism of 5-fluorouracil to an N-cholyl-2-fluoro-beta-alanine conjugate: previously unrecognized role for bile acids in drug conjugation. Proc. Natl. Acad. Sci. U. S. A. 84, 5439–5443 (1987).

19. Anthropic. Agent Skills. https://platform.claude.com/docs/en/agents-and-tools/agent-skills/overview.

20. Agent Skills. Agent Skills Specification. https://agentskills.io.

21. Damiani, T. et al. A universal language for finding mass spectrometry data patterns. Nat. Methods 22, 1247–1254 (2025).

22. Nash, W. J., Ngere, J. B., Najdekr, L. & Dunn, W. B. Characterization of electrospray ionization complexity in untargeted metabolomic studies. Anal. Chem. 96, 10935–10942 (2024).

23. Bittremieux, W. et al. Open access repository-scale propagated nearest neighbor suspect spectral library for untargeted metabolomics. Nat. Commun. 14, 8488 (2023).

24. Hagey, L. R. A Study on the Evolution and History of Bile Acids. Bile Acids - Discovering the World of Bile Acids https://bile.ucsd.edu/explore-the-book/.

25. Niu, J., et al. MinerU2.5: A decoupled vision-language model for efficient high-resolution document parsing. arXiv *[cs.CV]* (2025) doi:10.48550/arXiv.2509.22186.

26. Wang, B., et al. MinerU2.5-Pro: Pushing the limits of data-centric document parsing at scale. arXiv [cs.CV] (2026) doi:10.48550/arXiv.2604.04771.

27. Radford, A., et al. Robust speech recognition via large-scale weak supervision. arXiv [eess.AS] (2022) doi:10.48550/arXiv.2212.04356.

28. Qiao, X. et al. A tandem mass spectrometric study of bile acids: interpretation of fragmentation pathways and differentiation of steroid isomers. Steroids 77, 204–211 (2012).

29. Lan, K. et al. Key role for the 12-hydroxy group in the negative ion fragmentation of unconjugated C24 bile acids. Anal. Chem. 88, 7041–7048 (2016).

30. Wang, M. et al. Sharing and community curation of mass spectrometry data with Global Natural Products Social Molecular Networking. Nat. Biotechnol. 34, 828–837 (2016).

