## Supplementary Material for "Capturing human expertise for AI-assisted molecular structure annotation"

**Correspondence:**

### Supplementary Figures

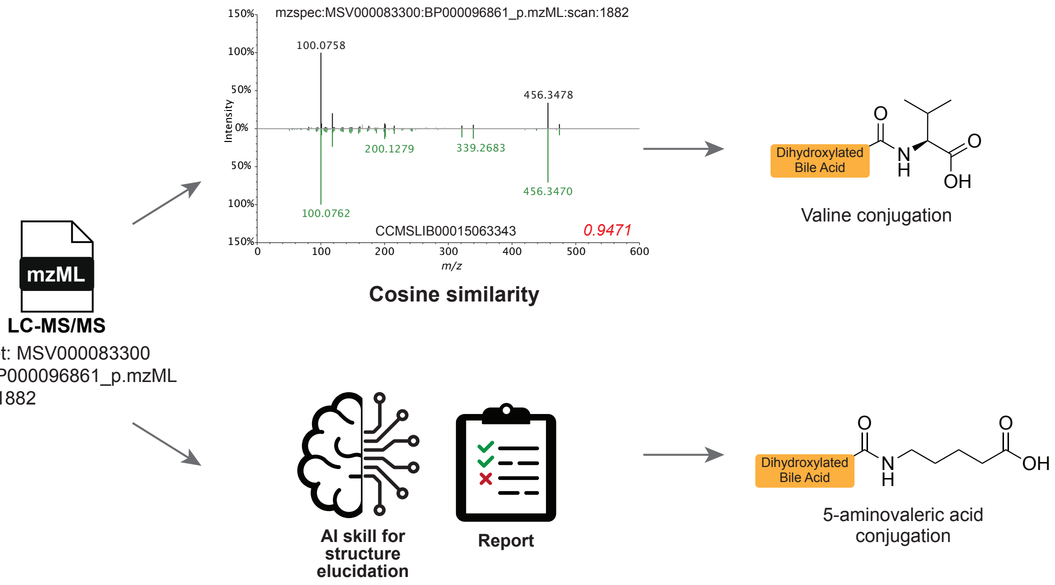


**Supplementary Figure 1. Expert-derived structural interpretation enables reassessment of spectral library-based annotations.** Comparison of two annotation strategies applied to the same experimental LC-MS/MS scan (scan 1882, MassIVE dataset MSV000083300). Spectral library matching (top) yielded a match to a valine-conjugated dihydroxylated bile acid (GNPS accession CCMSLIB00015063343). Independent interpretation of the experimental data using the bile acid annotation skill (bottom) instead favored a 5-aminovaleric acid conjugate.

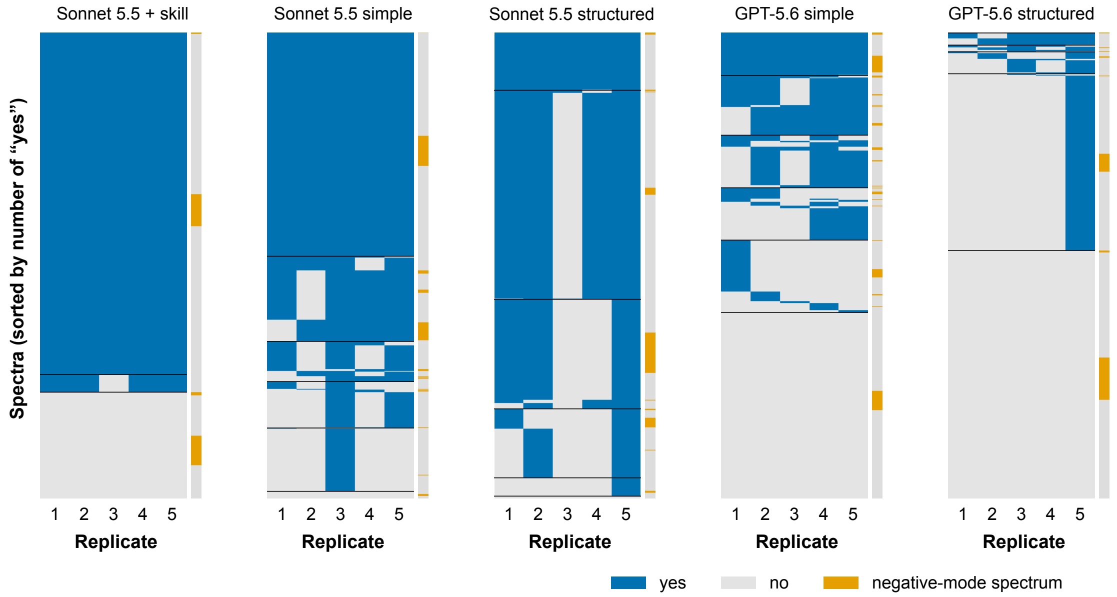


**Supplementary Figure 2.** Per-spectrum calls across the five replicates for each condition. Each panel shows the 4,953 benchmark spectra (all confirmed bile acids) for one condition, from left to right: Sonnet 5.5 + skill, Sonnet 5.5 simple prompt, Sonnet 5.5 structured prompt, GPT-5.6 simple prompt, and GPT-5.6 structured prompt. Rows represent spectrum and columns are replicates 1–5. Blue means the replicate identified the spectrum as a bile acid ("yes") and grey means it did not ("no"). Spectra are sorted from top to bottom by the number of replicates calling "yes" (5 to 0), and within each group by call pattern across replicates 1–5. Thin black lines separate the groups. The same row order is used for the strip to the right of each matrix, which marks negative-mode spectra in orange and positive-mode ones in grey. A fully reproducible condition therefore appears as solid blue and solid grey blocks, and spectra whose calls change between replicates appear as the mixed rows in between.

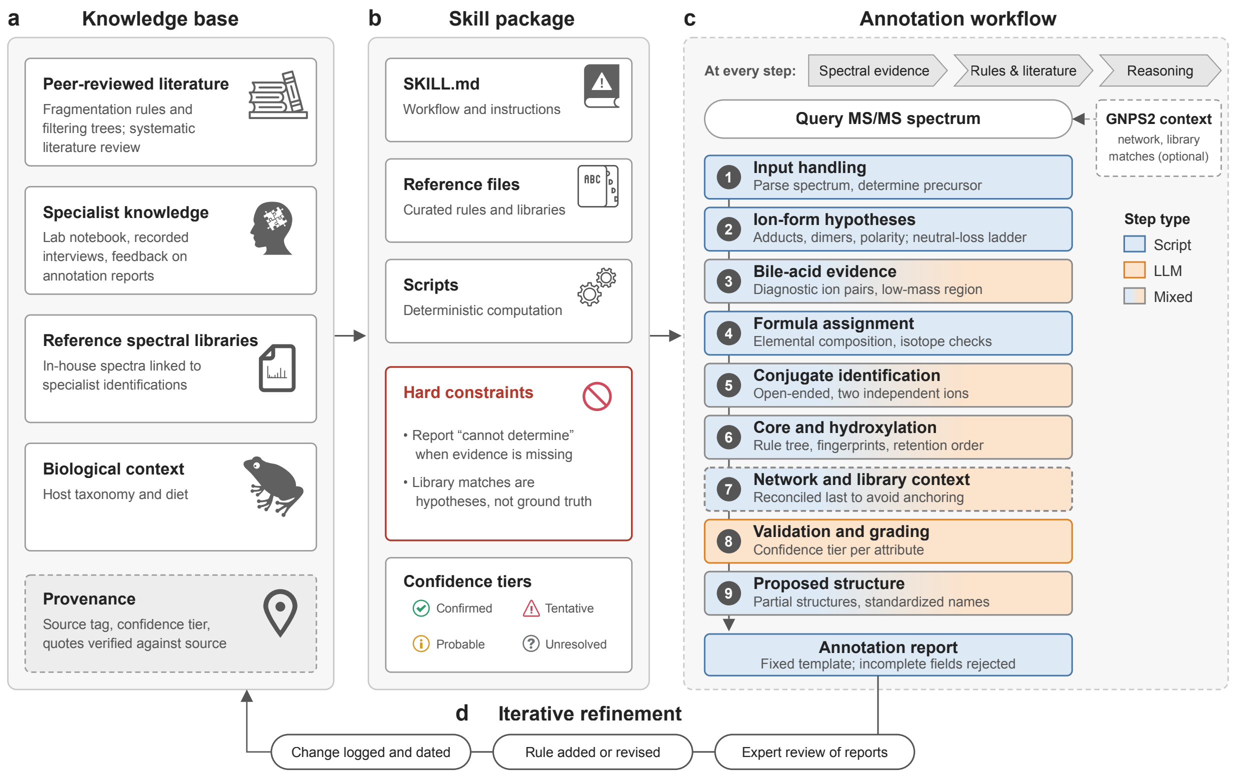


**Supplementary Figure 3.** Knowledge sources, skill package and annotation workflow. **(a)** Published, specialist, spectral-library and biological sources are recorded with provenance. **(b)** The skill combines instructions, reference files, scripts and explicit confidence rules. **(c)** Spectral evidence is linked to rules before structural reasoning; network and library context is reconciled after spectral interpretation. **(d)** Expert report review leads to dated rule revisions.

### Supplementary Methods

#### General-purpose language model benchmarking: prompting strategies

To assess bile acid recognition without access to the specialized annotation skill, Claude Sonnet 5.5 (Anthropic) and GPT-5.6 (OpenAI) were evaluated using two prompting strategies. The simplified prompt requested direct binary classification of MS/MS spectra, whereas the detailed prompt provided additional background on bile acid chemistry and instructed the models to consider precursor information, conjugation, fragmentation patterns, substituent positions, stereochemistry, and uncertainty.

Both prompting strategies were done in replicates (n = 5), ultimately generated CSV files containing a binary yes/no classification for each spectrum, associated with its scan number. The resulting classifications were matched to the corresponding entries in the anonymized BILELIB19 library. Recognition was calculated as the proportion of reference bile acid spectra classified as bile acids, overall and separately for positive and negative ionization modes.

All 4,953 benchmark spectra are confirmed bile acids, so recall was defined as the fraction of scans that a replicate classified correctly. For Sonnet 5.5 + skill, recall was calculated separately for each of the five replicates within each ion form, using the ion-form annotation from the library. The mean and standard deviation (SD) across replicates were then computed per ion form. Because the replicates are repeated runs on the same scans, the SD describes run-to-run variability, not uncertainty about the underlying recall. Some Ion forms are underrepresented in the library, with few scans (e.g., $[M-3H_2O+H]^+$, n = 3), which should be interpreted with caution.

Because the benchmark library consisted exclusively of reference bile acid spectra, this evaluation measured bile acid recognition sensitivity rather than specificity or overall classification accuracy. The detailed prompt also assumed that the input represented a bile acid and requested structural assignments, whereas the simplified prompt explicitly requested binary classification. These differences were retained as part of the evaluated prompting strategies.

**Consistency between consecutive replicates**

Consistency was measured as Cohen's κ, which corrects the observed agreement between two sets of yes/no calls for the agreement expected by chance given each replicate's yes-rate. For each condition, κ was computed over all 4,953 scans for each pair of consecutive replicates (1-2, 2-3, 3-4, 4-5), giving four values per condition. Each of the four other conditions was compared with Sonnet 5.5 + skill using a two-sided Welch's t-test, with Holm adjustment over the four comparisons. An exact Mann-Whitney test was run as a nonparametric check. Consecutive pairs share a replicate, so the four values per condition are not fully independent. They should be read as a descriptive comparison, not a formal inference.

The complete prompts used for the evaluations are reproduced below.

*Prompt 1. Simplified prompting*
Here is an mgf file with spectra that I want to annotate if they are bile acids or not. Inspect each of them and generate an output table with yes or no for each scan number.

*Prompt 2. Detailed prompting*
You are an expert in LC-MS/MS interpretation of bile acids. You will be given an MS/MS spectrum (image or peak list) of a bile acid. Your task is to assign the position and stereochemistry of each substituent on the steroid core, or state that it cannot be assigned from the evidence.

Background

- Bile acids are C24 steroids with a 5β-cholanic acid core (four fused rings, A–D) and a carboxylic acid side chain ending at C24.
- Hydroxyl or keto groups occur most often at C3, C6, C7 and C12, and less often at other positions, including the side chain (e.g. C23).
- Each hydroxyl can be α (below the ring plane) or β (above). Primary bile acids in humans are 3α-hydroxylated.
- Common examples: cholic acid (3α,7α,12α), chenodeoxycholic (3α,7α), deoxycholic (3α,12α), lithocholic (3α), ursodeoxycholic (3α,7β), hyodeoxycholic (3α,6α), muricholic acids (3α,6β,7α/β). Many other regio- and stereoisomers exist.
- The C24 acid is often conjugated with taurine or glycine, or carries other amine conjugates. Conjugation changes the precursor mass and adds conjugate-derived fragments, but does not by itself reveal core stereochemistry.
- Isomers share a formula and produce very similar fragment ions dominated by sequential water losses from the core. Differences between isomers, if present, are usually in relative intensities rather than unique peaks.
- A keto group and two hydroxyls can yield the same core fragment masses.

Reasoning steps (write these out before answering)

1. Precursor: determine m/z, adduct, polarity, and molecular formula. Count total oxygens and degrees of unsaturation.
2. Conjugate: identify the conjugate from the precursor mass and fragments.
3. Core substituents: infer the number of hydroxyl and keto groups from the precursor mass and the water-loss series.
4. Position and stereochemistry: reason from the fragment pattern and relative intensities toward the most likely positions and α/β configurations. Name the alternative isomers you considered and why you rejected them.
5. Confidence: if the evidence does not distinguish between candidates, say so rather than guessing.

Input

Instrument: {instrument}
Polarity / adduct: {adduct}
Collision energy: {collision_energy}
Precursor m/z: {precursor_mz}
Spectrum: {spectrum_image_or_peak_list}

Output

After your reasoning, return only this JSON:

{
assignment: "<e.g. 3α,7β-dihydroxy, taurine conjugate> or \"cannot assign\"",
substituents: [{"position": "C3", "group": "OH|keto", "stereo": "α|β|n/a"}],
conjugate: "<taurine|glycine|other|none>",
alternatives_considered: ["..."],

confidence: "high|medium|low"
}

Note:
for the input files, it is not guaranteed to always contain instrument and adduct information, as these are optional. Still follow the same Bile Acid predictions in prompt as best you can without this information if not provided.
