## Supplementary File 1 for "Capturing human expertise for AI-assisted molecular structure annotation"

### Bile-Acid MS/MS Annotation Report

BP000096861_p.mzML — Scan 1882 | Generated with the "bile-acid-msms-annotation-v57" skill

Run time: 6 min 52 s, start to finish (started 2026-11-26 01:10:24 UTC, report rendered 2026-11-26 01:17:16 UTC)

#### 1. Precursor / Input Summary

| Field | Value | How obtained |
|---|---|---|
| Precursor m/z | 474.35809 | mzML scan header |
| Ion form | [M+H−H2O]+, z = 1 (set after review of MS1 co-elution and MS2; the unmodified script chose [M+NH4]+) | MS1 scan 1877: [M+H]+ 492.36832 and [M+Na]+ 514.34845. The script's fixed 0.01 Da window had counted a 229.1596 peak (about 43 ppm off) in favour of [M+NH4]+. An explicit ion-form override was added in a working copy of the scripts and the pipeline was re-run (see C11, X14). |
| Neutral monoisotopic mass | 491.36138 | from the adduct assignment |
| Neutral formula candidates (±5 ppm, C/H/N≤3/O/S≤2) | C29H49NO5 (+0.62 ppm); also within 5 ppm: C30H53NS2 (-1.15 ppm) | formula_table.py (mass alone does not choose among these) |
| Retention time | 4.490 min (11.0 min total run) | mzML |
| Polarity / collision energy | positive ESI / HCD 30 | mzML; filter: FTMS + p ESI d Full ms2 474.3581@hcd30.00 [50.0000-500.0000] |
| Instrument | Q Exactive (Xcalibur 2.9-290033/2.9.0.2926; pwiz 3.0.9172) | mzML header |
| Isolation window | 474.360 –0.75/+0.75 Da; 1 MS1 ion(s) ≥1% inside it | mzML; MS1 scan 1877 |
| Peaks in MS2 | 120; base peak 100.0758 | mzML |
| Other MS2 on this precursor (whole run) | none | extract_context.py |

| Field | Value | How obtained |
|---|---|---|
| Engine cross-check | Pure evaluator and massql engine agree on every query | --engine both |

Collision energy 30 is below the NCE ~44–45 at which the dust-region rules were characterised. The dust region was still evaluated; expect fewer and weaker dust ions and some ratio shift. Lower energy alone does not lower a tier (stereochemistry_and_caveats.md §4).

#### 2. Annotated MS/MS Spectrum

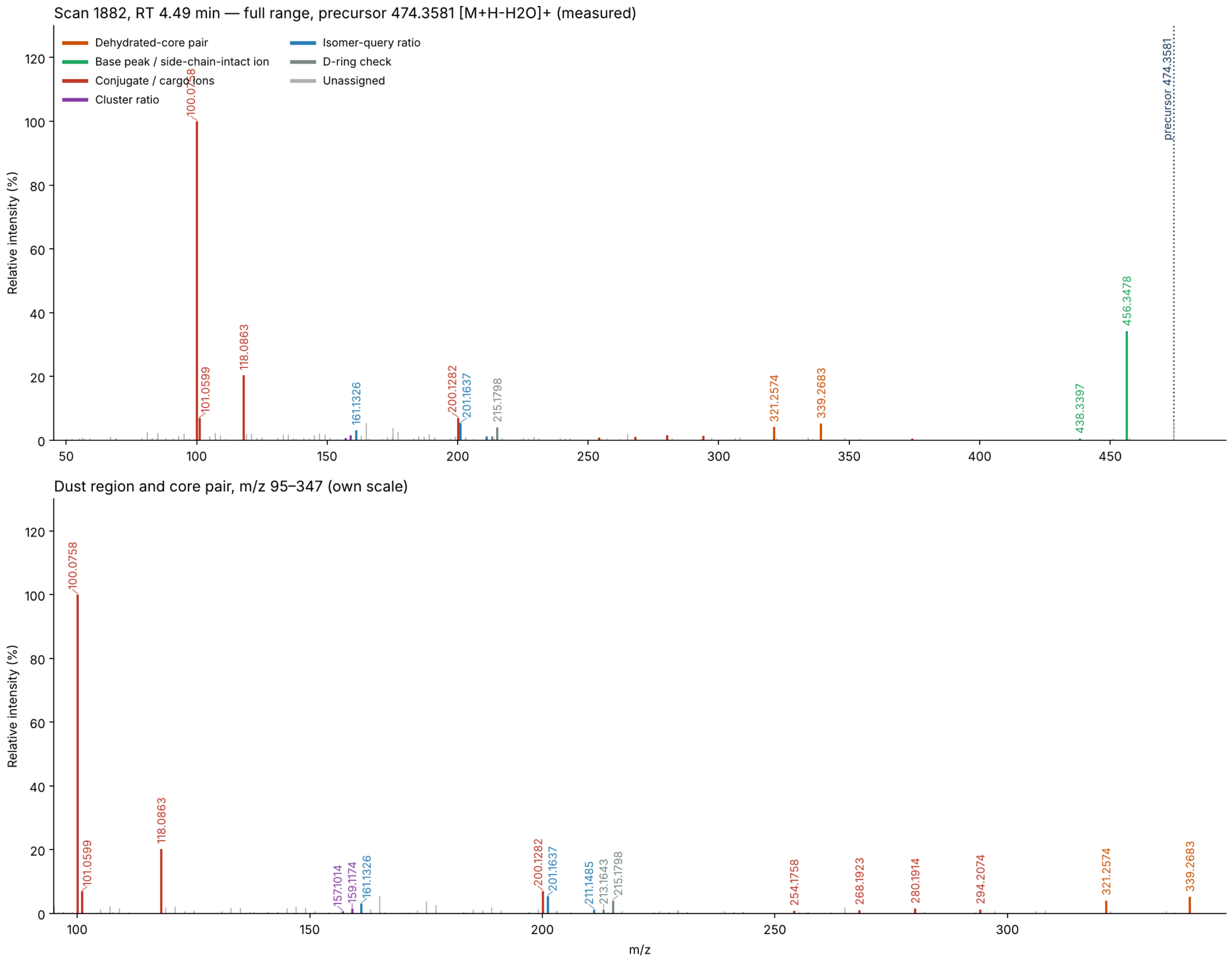


Scan 1882 (RT 4.49 min, precursor 474.3581 [M+H-H2O]+). Top: full range. Bottom: m/z 95–347 on its own scale, where the cargo ions and the ring-derived “dust” sit. Colors follow the shared palette; gray stems are unassigned peaks.

#### 3. Proposed Annotation

C24oic(OH)2–C5H11NO2 amide (positions 3,7 favoured) — Dihydroxy C24 bile acid carrying a C5H11NO2 amino acid; the 474.358 precursor is [M+H–H2O]+ of C29H49NO5 — i.e. **5-aminovaleric acid conjugate of a 3,7-dihydroxy C24 bile acid** tentative overall; other non-α C5H11NO2 isomers, other positions, and all α/β configurations are not excluded.

| Aspect | Call | Tier |
|---|---|---|
| Bile acid | Yes. Pair 339.268 (5.17%) / 321.257 (4.06%) passes only the 1% floor, not the published 5% floor; 26 dust ions | Tentative |
| Parent and ion form | $C_{29}H_{49}NO_5$; precursor is $[M+H-H_2O]^+$ | Probable |
| Core carbon count | C24 | Probable |
| Hydroxylation count | 2, saturated (no ketone) | Probable |
| Cargo composition | $C_5H_{11}NO_2$ amino acid; see A1 | Probable |
| Cargo identity | 5-aminovaleric acid favoured; see A2 – A5 | Tentative |
| Bond | C-24 amide; see A6 – A7 | Tentative |
| Hydroxyl positions | 3,7 favoured (fallback reading, no tree leaf) | Tentative |
| α/β at the hydroxyls, and C5 ring fusion | Not determined | Unresolved |

Standard C24 cholanoic core (core_structure_reference.md) with two hydroxyls and the C-24 carboxyl joined to the amino group of a C5 amino acid.

#### 4. Evidence Table

##### Confirmatory (positive) evidence

| Ref | Obs. m/z | %base | Assignment | Calc. | Δppm | Interpretation | Source |
|---|---|---|---|---|---|---|---|
| C1 | 339.2683<br>321.2574 | 5.17<br>4.06 | $C_{24}H_{35}O^+$<br>$C_{24}H_{33}^+$ | 339.2682<br>321.2577 | +0.2<br>-0.8 | Core-class query **C24 - Dihydroxy** passed: dehydrated-core pair present (query targets 339.27, 321.26, ±0.01 Da, ≥1% each — floor adapted from the published 5%; *passes only under the adapted floor (relaxed gate): core call and overall tier ≤ tentative; later steps graded on their own evidence*); the tree's **Dihydroxy_stage1** query passes on the same two ions — “the more hydroxyls, the smaller the number” | carbon_count_massql_table.md — [TrackSheet], [Mohanty2026-AC]; floor 5 % → 1 % [Adaptation-2026-10-18]; scripts/rules/massql_queries.yaml — [Mohanty2026-AC], [Transcript-UCSD9] |

| Ref | Obs. m/z | %base | Assignment | Calc. | Δppm | Interpretation | Source |
|---|---|---|---|---|---|---|---|
| C2 | 26 ring ions | ≥1 | odd-mass C/H/O | — | max \|3.8\| | Dust region m/z 100–250: 26 ring-derived ions (need 7) → passes: 101.0599 ($C_5H_9O_2^+$), 105.0700 ($C_8H_9^+$), 107.0856 ($C_8H_{11}^+$), 109.1011 ($C_8H_{13}^+$), 119.0856 ($C_9H_{11}^+$), 121.1014 ($C_9H_{13}^+$), 133.1012 ($C_{10}H_{13}^+$), 135.1164 ($C_{10}H_{15}^+$), 145.1014 ($C_{11}H_{13}^+$), 147.1168 ($C_{11}H_{15}^+$), 149.1326 ($C_{11}H_{17}^+$), 159.1174 ($C_{12}H_{15}^+$), 161.1326 ($C_{12}H_{17}^+$), 163.1121 ($C_{11}H_{15}O^+$), 173.1325 ($C_{13}H_{17}^+$), 175.1484 ($C_{13}H_{19}^+$), 177.1273 ($C_{12}H_{17}O^+$), 185.1324 ($C_{14}H_{17}^+$), 187.1485 ($C_{14}H_{19}^+$), 189.1279 ($C_{13}H_{17}O^+$), 189.1641 ($C_{14}H_{21}^+$), 199.1486 ($C_{15}H_{19}^+$), 201.1637 ($C_{15}H_{21}^+$), 211.1485 ($C_{16}H_{19}^+$), 213.1643 ($C_{16}H_{21}^+$), 215.1798 ($C_{16}H_{23}^+$). Set aside as N/S-bearing (cargo-like): 100.0758, 118.0863, 200.1282 — “the dust tells you where the things are attached. It doesn't tell you... if it was alpha or beta.” | dust_region_rules.md — [Specialist heuristic — Revision-2026-09-21], [Transcript-UCSDxx] |
| C3 | 474.3581 | MS1 | $[M+H-H_2O]^+$, z = 1 | — | — | Parent assignment $[M+H-H_2O]^+$: $[M+H]^+$ 492.3683 (calc. 492.3687) and $[M+Na]^+$ 514.3484 (calc. 514.3506) in MS1 scan 1877; isotope spacing 1.0053 Da. Tier probable; chosen on co-elution and fragment evidence (C11, X14), not on the script score. | MS1 scan 1877; reviewer-set ion form |

| Ref | Obs. m/z | %base | Assignment | Calc. | Δppm | Interpretation | Source |
|---|---|---|---|---|---|---|---|
| C4 | 211.1485<br>201.1637 | 1.18<br>5.33 | C16H19+<br>C15H21+ | 211.1481<br>201.1638 | +1.8<br>-0.6 | Di-3,7-OH\|Di-3,6-OH\|Di-3,12b-OH\|Di-7,12b-OH query PASSED (3,7-OH2; 3,6-OH2; 3,12b-OH2; 7,12b-OH2): obs 4.540× vs required 4.2× ±40% (2.52–5.88) — “a big 201 would mean it's a 3,7 [substitution]... a small 201 is dihydroxy [3,12, i.e. missing the 7-position].” Here 201/211 = 4.54, near the paper’s 7α example (about 4:1) and far from its 12α example (about 1:2). The tree did not reach this node (Section 5), so no published FDR applies. | scripts/rules/massql_queries.yaml — [Mohanty2026-AC], [Specialist heuristic] |
| C5 | 161.1326<br>201.1637 | 3.10<br>5.33 | C12H17+<br>C15H21+ | 161.1325<br>201.1638 | +0.9<br>-0.6 | Di-3,7-OH\|Di-3,12b-OH\|Di-7,12b-OH query PASSED (3,7-OH2; 3,12b-OH2; 7,12b-OH2): obs 1.719× vs required 2× ±20% (1.6–2.4) — *near a bound* — “keno always has a nice 201” | scripts/rules/massql_queries.yaml — [Mohanty2026-AC], [Notebook p.14], [Transcript-STE0xx] |
| C6 | 491.36138 (neutral) | — | C29H49NO5 | 491.36107 | +0.6 | C29H49NO5 = C24H40O4 (saturated dihydroxy C24 acid) + C5H11NO2 − H2O. It agrees with the core pair C1 and the cargo ions C7 . A keto analog would be 2 Da lighter. The C30H53NS2 alternative is set aside in X15 . — “close is not close enough... it has to be exact.” | precursor arithmetic (formula_table.py), [Specialist heuristic] |

| Ref | Obs. m/z | %base | Assignment | Calc. | Δppm | Interpretation | Source |
|---|---|---|---|---|---|---|---|
| C7 | 100.0758<br>118.0863<br>101.0599 | 100<br>20.30<br>6.87 | $C_5H_{10}NO^+$<br>$C_5H_{12}NO_2^+$<br>$C_5H_9O_2^+$ | 100.0757<br>118.0863<br>101.0597 | +1.0<br>+0.6<br>+2.0 | These are [X+H−H2O]+, [X+H]+ and [X+H−NH3]+ of the free $C_5H_{11}NO_2$ amino acid, in that order. A base peak at [AA+H−18] is the reported behaviour of terminal (ω-) amino acids such as 5-aminovaleric acid. α-Amino acids instead give [AA+H−46] ( X11 ). | literature_msms_rules.md §3.3 — [Garcia2025-JAFC] (QTOF, CE 30; LCA-core standard) |
| C8 | 456.3478<br>438.3397 | 34.14<br>0.43 | $C_{29}H_{46}NO_3^+$<br>$C_{29}H_{44}NO_2^+$ | 456.3472<br>438.3367 | +1.4<br>+6.9 | [M+H−2H2O]+ and [M+H−3H2O]+. These side-chain-intact ions keep the nitrogen, so the nitrogen unit is covalently bound. | diagnostic_core_ions.md §2; formula_table.py |
| C9 | 456.3478<br>−339.2683 | 34.14<br>5.17 | loss of<br>$C_5H_{11}NO_2$ | 117.0790<br>(obs. 117.0795) | 0.5 mDa<br>— | The step from the side-chain-intact ion to the core acylium is the whole $C_5H_{11}NO_2$ amino acid (observed 117.0795). Separately, the Step 0 ladder (C24, 2 OH) gives 474.3581 − 374.2821 = 100.0760, the base-peak residue (calc. 100.0757). The Step 0 route is capped at tentative. | conjugate_fragments.md amidate arithmetic; SKILL.md Step 0 B — [Benchmark201] |
| C10 | 200.1282<br>254.1758<br>268.1923<br>280.1914<br>294.2074<br>374.2693 | 6.98<br>0.76<br>1.04<br>1.47<br>1.27<br>0.50 | $C_{10}H_{18}NO_3^+$<br>$C_{14}H_{24}NO_3^+$<br>$C_{15}H_{26}NO_3^+$<br>$C_{16}H_{26}NO_3^+$<br>$C_{17}H_{28}NO_3^+$<br>$C_{23}H_{36}NO_3^+$ | 200.1281<br>254.1751<br>268.1907<br>280.1907<br>294.2064<br>374.2690 | +0.5<br>+3.0<br>+5.9<br>+2.5<br>+3.6<br>+0.9 | A series of cargo-retaining ions (all NO3). $C_{10}H_{18}NO_3^+$ has the size of a C20–C24 side chain still joined to the C5 amino acid. This reading is this report's interpretation, not a published rule. | formula_table.py; reasoning |

| Ref | Obs. m/z | %base | Assignment | Calc. | Δppm | Interpretation | Source |
|---|---|---|---|---|---|---|---|
| C11 | 474.358<br>492.368<br>514.348 | MS1<br>MS1<br>MS1 | precursor<br>[M+H]+<br>[M+Na]+ | —<br>—<br>— | —<br>—<br>— | In the cached MS1 scans, 492.368 is detected only at 4.46–4.50 min and 514.348 only at 4.48–4.49 min, inside the 474.358 peak. The 492/474 height ratio is not constant (about 0.08–0.2). The whole-run traces also show [M+H]+ and [M+Na]+ at a second peak (4.11 min), where 474.358 also peaks. | cached MS1 scans and XICs |
| C12 | 157.1014<br>159.1174<br>161.1326 | 0.63<br>1.53<br>3.10 | $C_{12}H_{13}^+$<br>$C_{12}H_{15}^+$<br>$C_{12}H_{17}^+$ | 157.1012<br>159.1168<br>161.1325 | +1.2<br>+3.8<br>+1.0 | 161 dominates (159/161 = 0.49). This does not look like the 3α,7α sketch (159 ≈ 161). The same shape appears in several library entries (3α,7β; 3α,6α/β; 3α,12α), so it cannot choose among them. Read at HCD 30. | hydroxylation_position_rules.md §A; fragment_pattern_fingerprints.md — [Notebook p.14], [Notebook-FragmentPatterns] |
| C13 | 215.1798<br>213.1643 | 3.98<br>1.23 | $C_{16}H_{23}^+$<br>$C_{16}H_{21}^+$ | 215.1794<br>213.1638 | +1.9<br>+2.3 | 215 is stronger than 213, so the “213 instead of 215” sign of extra D-ring unsaturation is not seen. Context only. | hydroxylation_position_rules.md §C–D — [Transcript-STE003] |

#### Exclusionary (negative) evidence — what this rules out

| Ref | Not observed / failed | Rules out | Source |
|---|---|---|---|
| X1 | Dihydroxy_stage2 failed: no peak within 0.01 Da of 321.2600 at >= 5% base peak | The Dihydroxy_stage2 bin (Amidated dihydroxy) | scripts/rules/massql_queries.yaml — [Mohanty2026-AC] |
| X2 | Di-1-ketone failed: no peak within 20 ppm of 161.1320 at >= 95% base peak | The Di-1-ketone bin (1 ketone) — below the node where the tree stopped, so this is the query's own result, not a tree step | scripts/rules/massql_queries.yaml — [Mohanty2026-AC] |

| Ref | Not observed / failed | Rules out | Source |
|---|---|---|---|
| X3 | Di-3,12a-OH\|Di-7,12a-OH failed: 201.163: obs 4.540× vs required 0.6× ±80% (0.12–1.08) | The Di-3,12a-OH\|Di-7,12a-OH bin (3,12a-OH2; 7,12a-OH2) — below the node where the tree stopped, so this is the query's own result, not a tree step | scripts/rules/massql_queries.yaml — [Mohanty2026-AC] |
| X4 | Di-3,12a-OH failed: 239.179: obs 1.803× vs required 1× ±30% (0.7–1.3) | The Di-3,12a-OH bin (3,12a-OH2) — below the node where the tree stopped, so this is the query's own result, not a tree step | scripts/rules/massql_queries.yaml — [Mohanty2026-AC] |
| X5 | Di-7,12a-OH failed: 239.179: obs 1.803× vs required 0.25× ±30% (0.175–0.325) | The Di-7,12a-OH bin (7,12a-OH2) — below the node where the tree stopped, so this is the query's own result, not a tree step | scripts/rules/massql_queries.yaml — [Mohanty2026-AC] |
| X6 | Di-3,7-OH failed: 173.132: obs 0.273× vs required 0.5× ±30% (0.35–0.65) | The Di-3,7-OH bin (3,7-OH2) — below the node where the tree stopped, so this is the query's own result, not a tree step | scripts/rules/massql_queries.yaml — [Mohanty2026-AC] |
| X7 | Di-3,12b-OH\|Di-7,12b-OH failed: 173.132: obs 0.273× vs required 1× ±20% (0.8–1.2) | The Di-3,12b-OH\|Di-7,12b-OH bin (3,12b-OH2; 7,12b-OH2) — below the node where the tree stopped, so this is the query's own result, not a tree step | scripts/rules/massql_queries.yaml — [Mohanty2026-AC] |
| X8 | Di-3,6-OH failed: 201.163: obs 1.719× vs required 0.7× ±60% (0.28–1.12) | The Di-3,6-OH bin (3,6-OH2) — below the node where the tree stopped, so this is the query's own result, not a tree step | scripts/rules/massql_queries.yaml — [Mohanty2026-AC] |

| Ref | Not observed / failed | Rules out | Source |
|---|---|---|---|
| X9 | Core-class rows absent: C24 - Nonhydroxy (343.3/325.29); C24 - Monohydroxy (341.28/323.27); C24 - Trihydroxy (337.25/319.24); C24 - Tetrahydroxy (335.24/317.23); C24 - Pentahydroxy (333.22/315.21); C23 - Monohydroxy (327.27/309.26/309.26/285.25); C23 - Dihydroxy (325.25/307.24/307.24/283.24); C23 - Trihydroxy (323.24/305.23/305.23/281.23); C23 - Tetrahydroxy (321.22/303.21/303.21/279.22); C23 - Pentahydroxy (319.21/301.19/301.19/277.19); C27 - Monohydroxy (383.33/159.12); C27 - Dihydroxy (381.32/159.12); C27 - Trihydroxy (379.29/159.12); C27 - Tetrahydroxy (377.28/159.12); C27 - Pentahydroxy (375.27/159.12); Aceto dihydroxy (347.294/329.284); C24 alkamine - Monohydroxy (359.295/341.284/341.284/359.295); C24 alkamine - Dihydroxy (357.279/339.268/339.268/357.279); C24 alkamine - Trihydroxy (355.263/337.253/337.253/355.263); C24 alkamine - Tetrahydroxy (353.248/335.237/335.237/353.248); C24 alkamine - Pentahydroxy (351.232/333.221/333.221/351.232); C28 alcohol - 1OH (125.13/143.14); C28 alcohol - 2OH (123.12/141.13); C28 alcohol - 3OH (121.1/139.11) | The hydroxylation counts and core classes those rows define | carbon_count_massql_table.md — [TrackSheet], [Mohanty2026-AC] |
| X10 | no 86.0964/104.1070/124.9998/184.0733 (PC / lysoPC head group); no 149.0233/167.0339 (phthalate); no 85.0284 (acylcarnitine); no 44.0262 Da repeat series (PEG) (±0.005 Da, ≥1% base) | Those non-bile-acid contaminant classes | nomenclature_glossary.md |
| X11 | No 72.0808 immonium ion ($C_4H_{10}N^+$), although the scan starts at m/z 50 | Disfavours valine and the other α-amino $C_5H_{11}NO_2$ isomers, which are expected to release [AA+H−46] | literature_msms_rules.md §3.3 — [Garcia2025-JAFC] |
| X12 | No 126.022 (taurine), 76.039 (glycine), 90.055 (alanine), 86.096 / 132.102 (leucine) | No support for taurine, glycine, alanine or leucine cargo, including the isobaric C23 + leucine split. The same absences give no support to ester-type $C_5H_{11}NO_2$ isomers that would release a smaller amino-acid ion | conjugate_fragments.md; reasoning |
| X13 | No prominent 209.1325 and no 309.2577 | Absent 209 rules nothing out. There is no 12-ketone marker | hydroxylation_position_rules.md §D, §D.1 |

| Ref | Not observed / failed | Rules out | Source |
|---|---|---|---|
| X14 | [M+NH4]+ reading of 474.358: the MS2 has no NH3-loss ion at 457.33, yet large fragments keep N; in MS1, 457.334 and 439.321 persist across 4.34–4.64 min while 474.358 is seen only around 4.36 (weak) and 4.46–4.52 min | Excludes the script's original [M+NH4]+ assignment of a C29H44O4 neutral | MS1/MS2 cache; reasoning |
| X15 | No $^{34}S$ A+2 peak on the precursor; core and side-chain-intact ions fit CHNO without a shared offset | Weighs against the C30H53NS2 neutral formula. The one S-fitting fragment (202.963, 0.57%) is not a bile-acid-type ion and is not used | stereochemistry_and_caveats.md §2a |
| X16 | No 357.2788 partner for the C24 dihydroxy alkamine pair | No support for a C24 alkamine reading | carbon_count_massql_table.md |
| X17 | No [M+NH4]+ trace for M = 491.361 (509.395) | Rules out nothing: the script grades ammonium as ambiguous, so it is not used to rank cargo isomers | dust_region_rules.md §5 |

#### Cargo evidence — what is attached to the bile acid

Every cargo candidate still in play, grouped by the question it answers. Each row cites the C/X rows above that support or argue against it; candidates already ruled out appear only in the exclusionary table. The Section 3 calls and Section 6 open questions are drawn from this table.

| Ref | Candidate | Evidence for | Evidence against | Status | Tier |
|---|---|---|---|---|---|
| **1. Cargo composition** → C5H11NO2 (residue C5H9NO) (probable) | | | | | |
| A1 | **C5H11NO2 amino acid**<br>1 N, 2 O | Exact parent mass; free ion and its losses; whole-unit neutral loss<br>C6 C7 C9 | Rests on the parent assignment<br>C3 | Supported | Probable |
| **2. Which C5H11NO2 isomer** → 5-aminovaleric acid favoured (tentative) | | | | | |
| A2 | **5-Aminovaleric acid**<br>ω-amino acid; free COOH | [AA+H−18] is the base peak; no α-type immonium<br>C7 X11 | No standard; further loss ion C5H7O+ (83.049) not seen; the rule comes from an LCA-core QTOF study | Favoured | Tentative |
| A3 | **3- or 4-aminopentanoic acid, or a branched non-α isomer**<br>same formula; free COOH | Same ions are not excluded<br>C7 | The literature rule does not test these isomers | Possible | Tentative |

| Ref | Candidate | Evidence for | Evidence against | Status | Tier |
|---|---|---|---|---|---|
| A4 | **Valine, norvaline or isovaline** α-amino; free COOH | Same composition; valine is the listed cargo in the delta table C6 | No 72.081 immonium; base peak is the −18 ion X11 C7 | Disfavoured | Tentative |
| A5 | **Ester-type isomer (e.g. GABA methyl ester, β-alanine ethyl ester)** no free acid | Same composition C6 | No smaller amino-acid ion seen; ammonium cannot help rank X12 X17 | Disfavoured | Tentative |
| **3. Bond to the bile acid** → C-24 amide favoured (tentative) | | | | | |
| A6 | **Amide at C-24** N-acyl amino acid | Cargo-retaining ions include a side-chain-sized C10H18NO3+ C10 C8 | Mass cannot tell an amide from a ring ester | Favoured | Tentative |
| A7 | **Ester on a ring hydroxyl** O-aminoacyl bile acid | Isobaric with the amide C6 | A side-chain-sized cargo ion is harder to explain C10 | Disfavoured | Tentative |

##### Step-7 validation pass

Every peak was re-checked against an elemental formula (C, H, N ≤3, O, S ≤2; even-electron ion, electron mass applied): 106 of 120 fit within ±10 ppm (largest |error| 9.6 ppm; ions used as evidence above: ≤1.8 ppm). 2 peak(s) are $^{13}C$ isotope peaks of a stronger ion and are not fragments. No plausible formula: 50.3208, 52.1982, 56.0501, 59.0497, 82.2944, 107.6347, 165.0892, 166.0955, 206.1273, 223.9152, 254.2027, 451.2414 — not used as evidence. Sulfur near-miss check (§2a): fragment 202.9629: CHNO C6H3O8 off -0.0193 Da (-95.2 ppm) → C6H3O6S fits. Shared CHNO offset: none. $^{34}S$ A+2 on the precursor: absent — no peak at the 34S position; weighs against S if the A+2 region was resolved.

Odd/even parity check (conjugate_fragments.md): 24 even-mass fragment(s) (88.0756, 100.0758, 118.0863, 138.0903, 154.0862, 200.1282, 214.1455, 254.1758, 262.1848, 268.1923, 280.1914, 282.2068, 294.2074, 306.2062, 308.2232, 322.2396, 334.2390, 336.2532, 348.2546, 353.9851, 374.2693, 438.3397, 456.3478, 474.3637) — each carries an odd number of N in its best formula. 0 odd-mass ion(s) carry an even number of N (e.g. N2). The nitrogen rule holds for every assigned ion.

#### 5. Why the Automated Tree Stalled at “Dihydroxy_stage1”

The pre-pass reached **Dihydroxy_stage1** (not a leaf): Dihydroxy_stage2 did not match. The unmet conditions:

- **Dihydroxy_stage2**: MS2PROD=321.26 resolves to **321.2600** — no peak within 0.01 Da of 321.2600 at >= 5% base peak. Target – H2O (303.2494) is also absent.

The stall is mechanical: 321.2574 is present within tolerance but at 4.06%, under the 5% floor of Dihydroxy_stage2. The two queries below it that passed on their own would point to the set 3,7 / 3,12β / 7,12β, but every leaf query failed, so even a completed path would not name an isomer. The position call stays *tentative* with no published FDR. MS2 at NCE 45 on this compound would be the way to test the tree.

#### 6. Unresolved Aspects

1. **Hydroxyl positions have no published tree leaf.** The tree stopped at Dihydroxy_stage1 (Section 5). The queries that passed on their own leave a set (3,7; 3,12β; 7,12β) (C4 , C5 ). All leaf queries failed ( X3 – X8 ). The 12β leaves need 173/175 ≈ 1.0 and the observed value is 0.273 ( X7 ). The 3,7 leaf also failed, but more narrowly: 0.273 against a floor of 0.35 ( X6 ). So 3,7 is a favoured reading, not a result. 3,6 failed its own query ( X8 ).
2. **Conflict between two position clues.** The notebook reads a 161-dominant cluster as 3α,12α-like ( C12 ). The 201/211 ratio of 4.54 ( C4 ) and the failed 12α queries ( X3 – X5 ) point away from 12α. The paper-derived ratio is given more weight, and the conflict is not resolved. The 3,7/3,12β query also passed only 7.4% above its lower bound (1.719 against 1.6); both engines agree.
3. **Stereochemistry is not determined.** This includes the α/β configuration at both hydroxyls and 5α versus 5β. The 161-dominant shape differs from the 3α,7α sketch, but that is too weak to call 7β (stereochemistry_and_caveats.md §1).
4. **Cargo isomer and bond.** 5-Aminovaleric acid is favoured over the other isomers ( A2 – A5 ) without a standard. A synthetic standard run by the same method (MS2 and retention time) would settle it. The same applies to amide versus ester ( A6 , A7 ).
5. **Relaxed-gate core call.** The published 5% pair floor was not met (321.257 is at 4.06%), so the published FDR does not apply and the core call and overall tier are capped at tentative.
6. **Collision energy.** HCD 30 is below the NCE of about 44–45 at which the dust and ratio rules were characterised. The rules were also built mainly on taurine and glycine amidates.
7. **Ammonium adduct is ambiguous.** The only ion 17.0265 below the precursor (457.334) does not track it ( X14 ), and no 509.395 trace was seen ( X17 ). If the parent has a free COOH, as proposed, a lack of ammonium adduct runs against the usual trend. It is a trend only and changes no tier.
8. **Unassigned ions.** 474.3637 (6.0%) sits 5.6 mDa above the header precursor and has no acceptable formula. It may be leftover precursor-region signal, but that is not shown. 165.0892 (5.4%) was flagged as a candidate cargo ion; the nearest formula, $C_{10}H_{13}O_2^+$, is 11 ppm off and it matches no known cargo ion. Neither is used. Other unassigned peaks are listed in the validation pass.
9. **Single MS2, no references.** There is no other MS2 on the 474.358 precursor in the run. There is no MS2 of 492.368 or 514.348 within ±10 s (the rest of the run was not searched for them). There is no GNPS2 context, and this run is not in the retention-time order library.

#### 7. Species / Taxonomic Context

Prior, not evidence. It changes no tier.

| Part | Finding | Source |
|---|---|---|
| **Host prior** | None available — no host or sample type recognisable in the file name, and none is recorded in the mzML. No prior was applied. | file name |
| **Expected producer** | 5-Aminovaleric acid conjugates have been reported among microbially conjugated bile acids from the human gut, and 5-aminovalerate amidates appear in public data. The host of this sample is unknown, so typicality cannot be judged. | [Garcia2025-JAFC]; [Mohanty2024-Cell] (literature_msms_rules.md) |
| **Plausible route** | Diet not provided. Microbial conjugation is the context given by the literature above; this spectrum cannot show origin. | source_and_diet_context.md §3 |

Follow-ups that would test the route:

- Record the host, sample type and diet for this file.
- Run a 5-aminovaleric acid conjugate standard (3,7-dihydroxy epimers) and a valine conjugate on the same method.
- Acquire MS2 at NCE 45 on 492.368 and 474.358.

#### 8. Chromatographic Context (Supporting, Not Evidence)

Extracted-ion chromatograms (±5 ppm) over the whole 11.0-min run, for M = 491.36138 ([M+H-H2O]+, measured (MS1 ion-form assessment)):

| Trace | Chromatographic peaks (RT min, apex intensity) |
|---|---|
| 492.3687 [M+H]+ | 4.11 (9.7e5), 4.48 (6.2e5) ← this scan, 5.42 (3.4e5), 5.89 (8.4e5), 6.00 (5.8e5), 6.14 (2.2e5), 6.50 (6.7e4) |
| 509.3952 [M+NH4]+ | no peak above 2% of the trace maximum (trace empty) |
| 514.3506 [M+Na]+ | 4.11 (1.2e5), 4.48 (5.1e5) ← this scan |
| 530.3245 [M+K]+ | 2.71 (9.6e5) |
| 474.3581 [M+H-H2O]+ (precursor) | 3.30 (3.1e5), 4.11 (6.3e5), 4.34 (1.9e5), 4.49 (6.0e6) ← this scan, 5.96 (3.6e5) |

Supporting orientation only: retention time alone identifies nothing (stereochemistry_and_caveats.md). No other MS2 was acquired on this precursor in the run.

#### 9. Diagnostic Decision Tree

Each node's accent color is the color of the corresponding ion family highlighted in the Section 2 spectrum. The numbered chips (C = confirmatory, X = exclusionary, A = cargo evidence) are the Ref column of the Section 4 evidence tables, so every node traces back to its ions, mass errors and sources.

Dehydrated-core pair · Base peak / side-chain-intact ion · Conjugate / cargo ions · Position indicator · Cluster ratio · Isomer-query ratio · D-ring check · Grouping / precursor arithmetic · Ruled out (no diagnostic ion family)

✓ supports the annotation · ⊘ alternative ruled out (dashed box) · – context only

**MS2 scan 1882**
precursor 474.3581 [M+H−H2O]+ · 120 peaks · base peak 100.0758

**Core identity & conjugate**
carbon count, OH count, cargo

- ✓ **Parent C29H49NO5** — [M+H]+ and [M+Na]+ in MS1 — C3 C6 C11
- ✓ **C24 dihydroxy core pair** — relaxed gate; 26 dust ions — C1 C2
- ✓ **C5H11NO2 cargo** — 118.086, base 100.076, 117.079 loss — C7 C8 C9 A1
- ✓ **5-Aminovaleric favoured** — −18 ion dominant, no 72.081 — C7 X11 A2
- ✓ **C-24 amide favoured** — side-chain-sized cargo ion — C10 A6

**Hydroxylation position**
tree stalled at stage 1 (Section 5)

- ✓ **201/211 = 4.54** — 7α-example-like, not 12α-like — C4
- – **3,7 / 3,12β / 7,12β query** — set only; near bound; 3,7 closest leaf — C5 X6 X7
- – **161-dominant cluster** — shape not specific; conflicts for 12α — C12

**Alternatives ruled out or unsupported**
Section 4, exclusionary table

- ⊘ **[M+NH4]+ of C29H44O4** — N retained; companions do not track — X14
- ⊘ **12α bins** — leaf ratios failed — X3 X4 X5
- ⊘ **3,6-(OH)2 query** — 201/161 outside range — X8
- – **α-Amino isomers disfavoured** — no immonium; not fully excluded — X11 A4

↓

**3,7-dihydroxy C24 bile acid – 5-aminovaleric acid amide (Tentative)**

C24, 2 OH and C5H11NO2 are probable; isomer, bond and positions tentative; α/β unresolved

#### 10. Proposed Final Structures

One row per structure the reasoning in Sections 3–9 still supports, drawn with RDKit with the ABCD rings in the standard bile-acid orientation. A partial structure draws only what the evidence claims: wavy bonds mark stereocentres that are not resolved, substituents that are counted but not placed are listed under the drawing and their possible rings shaded, and a highlighted unit is weaker than the rest (the Notes say why). The Tier column is copied from the reasoning. Names are standardized with the metabolite-annotation-cleanup rules (PubChem's title when the exact, stereo-defined structure's record lists the proposed name; otherwise the cleaned label); standardization never changes a tier.

| Structure | Name | Tier | Notes |
|---|---|---|---|
| 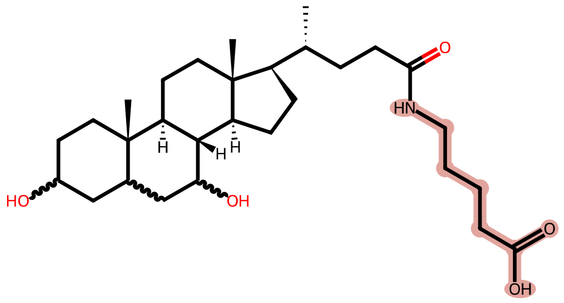 | **3,7-dihydroxycholan-24-oyl-5-aminovaleric acid** partial structure<br>$C_{29}H_{49}NO_5$ · 491.36107 Da<br>AALAHUXOORKJHH-CTAHYTHFSA-N | Tentative | The only structure drawn; C3, C5 and C7 configuration left open.<br>Wavy bonds: stereochemistry not resolved at C3, C5, C7<br>Highlighted unit: $C_5H_{11}NO_2$ composition is probable; the straight-chain 5-amino isomer and the amide bond are tentative.<br>Not resolved:<br>• α/β at C3 and C7, and 5α/5β<br>• Positions: 3,12β and 7,12β remain in a passed query set; no leaf was reached<br>• Cargo isomer: other non-α $C_5H_{11}NO_2$ amino acids; α-isomers disfavoured<br>• Amide versus ring ester<br>Assumed, not shown by this spectrum:<br>• Usual backbone configuration (8β, 9α, 10β, 13β, 14α, 17β) and 20R; this spectrum does not show it<br>Name: no PubChem record for this structure; cleaned label shown |

*Generated by the "bile-acid-msms-annotation-v57" skill (deterministic MassQL pre-pass, engine cross-checked) plus literature-grounded reasoning in the referenced skill files. Source: BP000096861_p.mzML, scan 1882. The ion form was set to $[M+H-H_2O]^+$ after review, using an override added to a working copy of ms1_context.py. Run time: 6 min 52 s, start to finish (started 2026-11-26 01:10:24 UTC, report rendered 2026-11-26 01:17:16 UTC).*
