## Supplementary File 2 for "Capturing human expertise for AI-assisted molecular structure annotation"

### Bile-Acid MS/MS Annotation Report

P3_C5_Sample_x3160658.mzML — Scan 3663 | Generated with the bile-acid-msms-annotation-v5-6 skill

Run time: 5 min 3 s, start to finish (started 2026-10-27 20:26:06 UTC, report rendered 2026-10-27 20:31:09 UTC)

#### 1. Precursor / Input Summary

| Field | Value | How obtained |
|---|---|---|
| Precursor m/z | 672.55323 | mzML scan header |
| Ion form | Unresolved; [M+H]+ used only as a computational fallback. [M+NH4]+ remains a formula-compatible alternative. | ion form unresolved from MS1: no adduct is distinguishable from the MS1 evidence: [M+H]+, [M+NH4]+, [M+Na]+, [M+K]+, [M+H-H2O]+, [2M+H]+, [2M+Na]+ are all consistent with the observed ion |
| Neutral formula candidates (±5 ppm, C/H/N≤3/O/S≤2) | C38H77N3O2S2 (+0.34 ppm); also within 5 ppm: C46H73NS (-0.63 ppm); C37H73N3O7 (+1.64 ppm); C42H73NO5 (-4.35 ppm); these are under the [M+H]+ fallback, not an established neutral formula. | formula_table.py (mass alone does not choose among these) |
| Retention time | 9.119 min (10.0 min total run) | mzML |
| Polarity / collision energy | positive ESI / HCD 30 | mzML; filter: FTMS + p ESI d Full ms2 672.5532@hcd30.00 [50.0000-705.0000] |
| Instrument | Q Exactive (Xcalibur 2.9-290033/2.9.0.2926; pwiz 3.0.23118) | mzML header |
| Isolation window | 672.553 ±1.5 Da; purity not established | mzML; nearest MS1 includes 672.55322. Prefill reported zero ions despite the cached precursor detection; that count is not used. |
| Peaks in MS2 | 98; base peak 355.2625 | mzML |
| Other MS2 on this precursor (whole run) | scan 3129 @ 7.768 min, scan 3201 @ 7.948 min, scan 3363 @ 8.363 min, scan 3433 @ 8.533 min, scan 3503 @ 8.709 min, scan 3737 @ 9.291 min | extract_context.py |
| Engine cross-check | not run (pure evaluator only; no borderline ratio) | --engine pure (default) |

Collision energy 30 is below the NCE ~44–45 at which the dust-region rules were characterised. The dust region was still evaluated; expect fewer and weaker dust ions and some ratio shift. Lower energy alone does not lower a tier (stereochemistry_and_caveats.md §4).

#### 2. Annotated MS/MS Spectrum

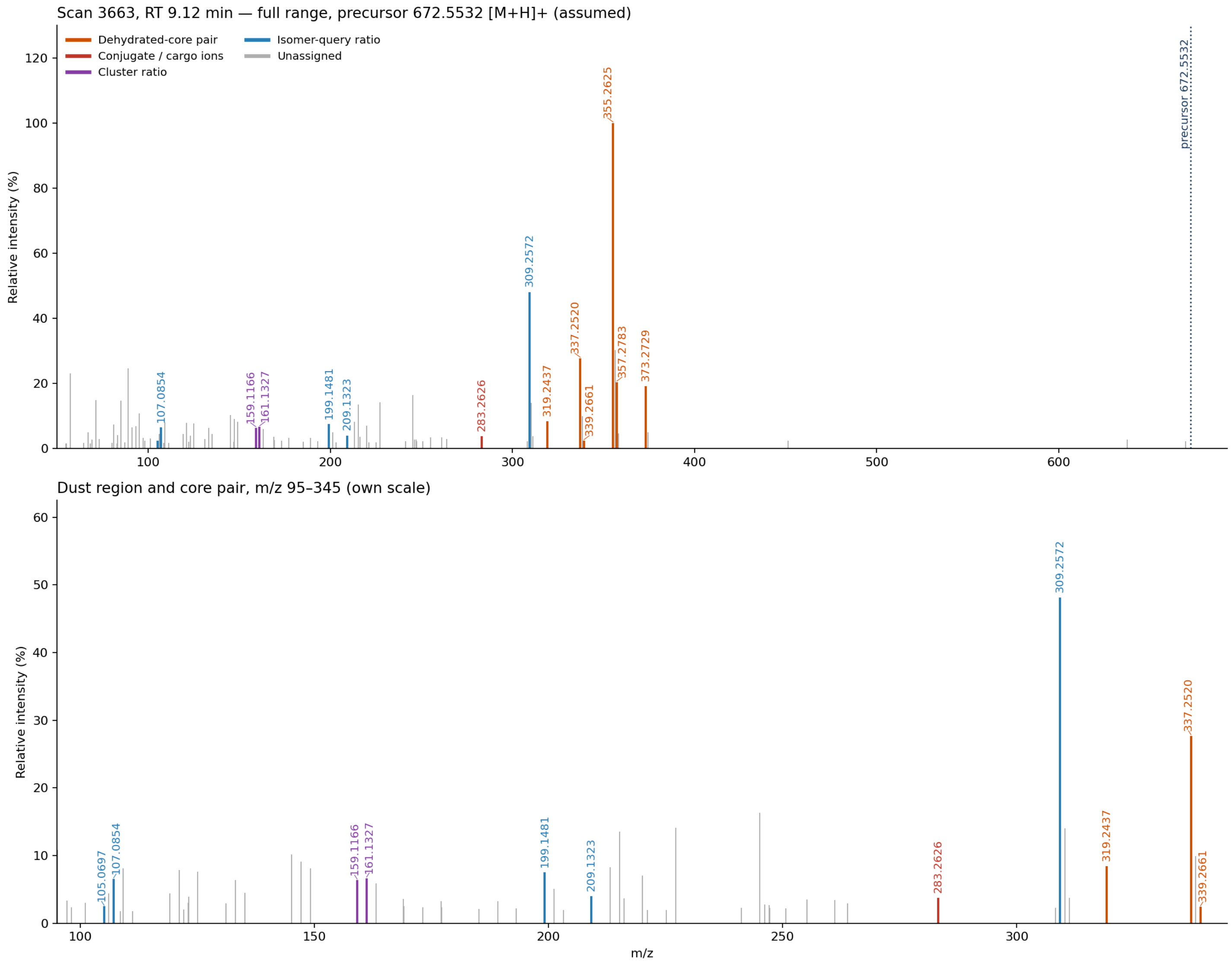


Scan 3663 (RT 9.12 min, precursor 672.5532 [M+H]+). Top: full range. Bottom: m/z 95–345 on its own scale, where the cargo ions and the ring-derived “dust” sit. Colors follow the shared palette; gray stems are unassigned peaks.

#### 3. Proposed Annotation

Modified C24 bile acid; 12-oxo-like fragmentation — Bile-acid core probable; C18-derived modification and exact ion form unresolved — i.e. **C24 bile-acid derivative, possibly a C18:1 fatty-acyl conjugate** Overall tentative. A 12-oxo-like core is favored by fragment ratios, but no terminal isomer leaf was reached. Neither a specific cargo nor an intact structure is identified.

| Aspect | Call | Tier |
|---|---|---|
| Bile-acid family / carbon count | C24 bile-acid core supported by the ladder and abundant ring fragments C1 – C4 , C8 | Probable |
| Core oxygenation | Trihydroxy-equivalent pattern; hydroxy-keto reading favored; true OH count remains open | Tentative |
| Ketone position | 12-oxo-like fragmentation: 309.2572 is strong and the 309/319 query passes C6 ; not a reached leaf | Tentative |

| Aspect | Call | Tier |
|---|---|---|
| Ion form | Unresolved: [M+H]+ fallback versus [M+NH4]+ or dehydrated ammonium hypotheses C11 | Unresolved |
| Cargo | C18-derived unit; C18:1 fatty-acyl ester candidate — see A1–A4 | Tentative |
| Exact compound / stereochemistry | Cannot determine linkage, specific bile acid, fatty-chain isomer or stereochemistry | Unresolved |

Standard C24 bile-acid connectivity supported by the oxygenated carbon-retaining fragment ladder. O-acyl hydroxy-keto and N-acyl alkamine scaffolds are hypotheses, not resolved structures.

#### 4. Evidence Table

##### Confirmatory (positive) evidence

| Ref | Obs. m/z | %base | Assignment | Calc. | Δppm | Interpretation | Source |
|---|---|---|---|---|---|---|---|
| C1 | 337.2520<br>319.2437 | 27.69<br>8.40 | $C_{24}H_{33}O^+$<br>$C_{24}H_{31}^+$ | 337.25259<br>319.24203 | -1.71<br>+5.28 | Core-class query **C24 - Trihydroxy** passed: dehydrated-core pair present (query targets 337.25, 319.24, ±0.01 Da, ≥1% each — floor adapted from the published 5%); the tree's **Trihydroxy_stage1** query passes on the same two ions — "the more hydroxyls, the smaller the number" | carbon_count_massql_table.md — [TrackSheet], [Mohanty2026-AC]; floor 5 % → 1 % [Adaptation-2026-10-18]; scripts/rules/massql_queries.yaml — [Mohanty2026-AC], [Transcript-UCSD9] |
| C2 | 357.2783<br>339.2661<br>339.2661<br>357.2783 | 20.38<br>2.39<br>2.39<br>20.38 | $C_{24}H_{37}O_2^+$<br>$C_{24}H_{35}O^+$<br>$C_{24}H_{35}O^+$<br>$C_{24}H_{37}O_2^+$ | 357.2788<br>339.26824<br>339.26824<br>357.2788 | -1.5<br>-6.37<br>-6.37<br>-1.5 | Core-class query **C24 alkylamine - Dihydroxy** passed: dehydrated-core pair present (query targets 357.279, 339.268, 339.268, 357.279, ±0.01 Da) — "the more hydroxyls, the smaller the number" Screening overlap: this query does not itself establish nitrogen on the ring or discriminate an N-linked cargo from an oxygenated-core dehydration ladder. | carbon_count_massql_table.md — [TrackSheet], [Mohanty2026-AC], [Transcript-UCSD9] |
| C3 | 355.2625<br>337.2520<br>337.2520<br>355.2625 | 100.00<br>27.69<br>27.69<br>100.00 | $C_{24}H_{35}O_2^+$<br>$C_{24}H_{33}O^+$<br>$C_{24}H_{33}O^+$<br>$C_{24}H_{35}O_2^+$ | 355.2632<br>337.25259<br>337.25259<br>355.2632 | -1.7<br>-1.71<br>-1.71<br>-1.7 | Core-class query **C24 alkylamine - Trihydroxy** passed: dehydrated-core pair present (query targets 355.263, 337.253, 337.253, 355.263, ±0.01 Da) — "the more hydroxyls, the smaller the number" Screening overlap: this query does not itself establish nitrogen on the ring or discriminate an N-linked cargo from an oxygenated-core dehydration ladder. | carbon_count_massql_table.md — [TrackSheet], [Mohanty2026-AC], [Transcript-UCSD9] |

| Ref | Obs. m/z | %base | Assignment | Calc. | Δppm | Interpretation | Source |
|---|---|---|---|---|---|---|---|
| C4 | 41 ring ions | ≥1 | odd-mass C/H/O | — | max \|10.0\| | Dust region m/z 100–250: 41 ring-derived ions (need 7) → passes: 101.0593 (C5H9O2+), 105.0697 (C8H9+), 107.0854 (C8H11+), 109.1011 (C8H13+), 111.1159 (C8H15+), 119.0853 (C9H11+), 121.1011 (C9H13+), 123.0804 (C8H11O+), 123.1165 (C9H15+), 125.0958 (C8H13O+), 131.0848 (C10H11+), 133.1007 (C10H13+), 135.0803 (C9H11O+), 135.1171 (C10H15+), 145.1011 (C11H13+), 147.0807 (C10H11O+), 147.1164 (C11H15+), 149.1323 (C11H17+), 159.1166 (C12H15+), 161.1327 (C12H17+), 163.1481 (C12H19+), 169.1012 (C13H13+), 173.1336 (C13H17+), 177.1638 (C13H21+), 185.1319 (C14H17+), 189.1632 (C14H21+), 193.1214 (C12H17O2+), 199.1481 (C15H19+), 201.1632 (C15H21+), 203.1774 (C15H23+), 209.1323 (C16H17+), 213.1640 (C16H21+), 215.1793 (C16H23+), 221.1541 (C14H21O2+), 225.1637 (C17H21+), 227.1429 (C16H19O+), 227.1780 (C17H23+), 241.1598 (C17H21O+), 245.1533 (C16H21O2+), 247.1687 (C16H23O2+), 247.2397 (C18H31+). Set aside as N/S-bearing (cargo-like): 106.0499, 122.0603, 145.0603, 169.0257, 220.0376 — "the dust tells you where the things are attached. It doesn't tell you... if it was alpha or beta." | dust_region_rules.md — [Specialist heuristic — Revision-2026-09-21], [Transcript-UCSDxx] |
| C5 | 107.0854<br>105.0697<br>199.1481<br>209.1323 | 6.52<br>2.48<br>7.51<br>4.00 | C8H11+<br>C8H9+<br>C15H19+<br>C16H17+ | 107.0855<br>105.0699<br>199.1481<br>209.1325 | -1.2<br>-1.6<br>-0.3<br>-0.7 | Tri-All_ketone_NEW query PASSED (1 hydroxyl and 1 ketone on bile acid core; 3,12,16-OH3): obs 0.380× vs required 0.55× ±40% (0.33–0.77); obs 0.533× vs required 0.6× ±45% (0.33–0.87) This is a standalone passing query below a failed upstream gate, not a reached isomer leaf. At HCD 30 it is supporting, tentative evidence only. | scripts/rules/massql_queries.yaml — [Mohanty2026-AC] |
| C6 | 319.2437<br>309.2572 | 8.40<br>48.10 | C24H31+<br>C23H33+ | 319.24203<br>309.2577 | +5.28<br>-1.4 | Tri-3a_12k query PASSED (3a,12-oxo): obs 5.726× vs required 5.5× ±20% (4.4–6.6) — "big 309, it's a 12-ketone due to C-ring stabilization" This is a standalone passing query below a failed upstream gate, not a reached isomer leaf. At HCD 30 it is supporting, tentative evidence only. | scripts/rules/massql_queries.yaml — [Mohanty2026-AC], [Notebook p.28/31] |

| Ref | Obs. m/z | %base | Assignment | Calc. | Δppm | Interpretation | Source |
|---|---|---|---|---|---|---|---|
| C7 | 671.54595 (neutral) | — | Formula/adduct alternatives, not a unique neutral formula | 672.55615 (candidate ion) | −4.34 | C42H73NO5 as [M+H]+ and C42H70O5 as [M+NH4]+ have exactly the same ion composition C42H74NO5+. Both calculate to 672.55615: measured error −0.00292 Da (−4.34 ppm). C42H72O6 as [M+NH4−H2O]+ also has that composition. The [M+H]+ neutral C42H73NO5 is −0.00292 Da (−4.35 ppm) from the fallback neutral mass. Mass-only formulas with RDBE 2 or 3 cannot contain a four-ring steroid; C46H73NS has insufficient oxygen for the oxygenated-core products. These alternatives assume the core and precursor are from the same species. — “close is not close enough... it has to be exact.” | precursor arithmetic (formula_table.py), [Specialist heuristic] |
| C8 | 373.27289<br>355.26254<br>309.25723 | 19.058<br>100.0<br>48.104 | C24H37O3+<br>C24H35O2+<br>C23H33+ | 373.27372<br>355.26316<br>309.25768 | -2.23<br>-1.73<br>-1.44 | C24H37O3+ → C24H35O2+ is a water-loss pair; C23H33+ at 309.2572 is 48.10% of base. Together these support an oxygenated C24 core and the tentative 12-oxo-like reading. The same products may be reached through other precursor oxygenation or conjugation pathways. | diagnostic_core_ions.md; hydroxylation_position_rules.md §B/§D; [Notebook p.28/31]; [Mohanty2026-AC] |
| C9 | 283.26257 | 3.767 | C18H35O2+ | 283.26316 | -2.06 | C18H35O2+ fits protonated C18:1 fatty acid (−0.00058 Da, −2.06 ppm). This is only one cargo-like product: no second independent C18:1 cargo marker or intact-cargo loss is established. It therefore supports a fatty-acyl hypothesis, not a named fatty acid. | formula_table.py; SKILL.md Step 3 two-independent-ion requirement |
| C10 | 159.11661<br>161.13274 | 6.357<br>6.651 | C12H15+<br>C12H17+ | 159.11683<br>161.13248 | -1.38<br>1.61 | 159 and 161 are nearly equal (161/159 = 1.046), compatible with a 3,7-like heuristic cluster. The 3a7a sketch has low-medium digitization confidence and ambiguous rank labels; it does not distinguish this hydroxy-keto/conjugated spectrum. No position or stereo assignment follows. | hydroxylation_position_rules.md §A; fragment_pattern_fingerprints.md; [Notebook-FragmentPatterns] |
| C11 | 672.55322 (MS1) | MS1 | z=1 in scan header; adduct unresolved | — | — | Nearest MS1 scan 3659 contains the target, but no discriminating protonated/ammoniated/sodiated companion was found. The observed nearby spacing 0.99987 Da is not a clean 1.00335-Da carbon-isotope spacing, so isotope evidence is not used to settle ion form. [M+H]+ remains an explicit fallback, not a measurement. | context_3663.json ion_form; SKILL.md Step 0 |

#### Exclusionary (negative) evidence — what this rules out

| Ref | Not observed / failed | Rules out | Source |
|---|---|---|---|
| X1 | Trihydroxy_stage2 failed: no peak within 0.01 Da of 282.2762 at >= 2% base peak | This specific query/bin was not satisfied; not a universal exclusion of the molecule under a different ion form or cargo. The Trihydroxy_stage2 bin (Amidated trihydroxy) | scripts/rules/massql_queries.yaml — [Mohanty2026-AC] |
| X2 | Tri-3k_7or12_OH failed: 213.163: obs 1.102× vs required 2× ±20% (1.6–2.4) | This specific query/bin was not satisfied; not a universal exclusion of the molecule under a different ion form or cargo. The Tri-3k_7or12_OH bin (3oxo,7-OH; 3-oxo,12-OH) — below the node where the tree stopped, so this is the query's own result, not a tree step | scripts/rules/massql_queries.yaml — [Mohanty2026-AC] |
| X3 | Tri-3,6,7_NEW failed: no peak within 20 ppm of 191.1430; reference variable 'Y' was never set -- its reference ion was not found; 213.163: obs 1.102× vs required 0.75× ±20% (0.6–0.9) | This specific query/bin was not satisfied; not a universal exclusion of the molecule under a different ion form or cargo. The Tri-3,6,7_NEW bin (3,6,7-OH3) — below the node where the tree stopped, so this is the query's own result, not a tree step | scripts/rules/massql_queries.yaml — [Mohanty2026-AC] |
| X4 | Tri-3,6b,7a-OH\|Tri-3,6a,7b-OH failed: 161.132: obs 1.046× vs required 0.9× ±10% (0.81–0.99) | This specific query/bin was not satisfied; not a universal exclusion of the molecule under a different ion form or cargo. The Tri-3,6b,7a-OH\|Tri-3,6a,7b-OH bin (3,6b,7a-OH3; 3,6a,7b-OH3) — below the node where the tree stopped, so this is the query's own result, not a tree step | scripts/rules/massql_queries.yaml — [Mohanty2026-AC] |
| X5 | Tri-3,6b,7a-OH failed: 309.257: obs 6.344× vs required 1× ±20% (0.8–1.2) | This specific query/bin was not satisfied; not a universal exclusion of the molecule under a different ion form or cargo. The Tri-3,6b,7a-OH bin (3,6b,7a-OH3) — below the node where the tree stopped, so this is the query's own result, not a tree step — “big 309, it's a 12-ketone due to C-ring stabilization” | scripts/rules/massql_queries.yaml — [Mohanty2026-AC], [Notebook p.28/31] |

| Ref | Not observed / failed | Rules out | Source |
|---|---|---|---|
| X6 | Tri-3,6a,7a-OH failed: 185.132: obs 0.912× vs required 1× ±5% (0.95–1.05) | This specific query/bin was not satisfied; not a universal exclusion of the molecule under a different ion form or cargo. The Tri-3,6a,7a-OH bin (3,6a,7a-OH3) — below the node where the tree stopped, so this is the query's own result, not a tree step | scripts/rules/massql_queries.yaml — [Mohanty2026-AC] |
| X7 | Tri-3,7a,12a-OH_NEW failed: 199.148: obs 1.877× vs required 0.6× ±45% (0.33–0.87) | This specific query/bin was not satisfied; not a universal exclusion of the molecule under a different ion form or cargo. The Tri-3,7a,12a-OH_NEW bin (3,7a,12a-OH3) — below the node where the tree stopped, so this is the query's own result, not a tree step | scripts/rules/massql_queries.yaml — [Mohanty2026-AC] |
| X8 | Core-class rows absent: C24 - Nonhydroxy (343.3/325.29); C24 - Monohydroxy (341.28/323.27); C24 - Dihydroxy (339.27/321.26); C24 - Tetrahydroxy (335.24/317.23); C24 - Pentahydroxy (333.22/315.21); C23 - Monohydroxy (327.27/309.26/309.26/285.25); C23 - Dihydroxy (325.25/307.24/307.24/283.24); C23 - Trihydroxy (323.24/305.23/305.23/281.23); C23 - Tetrahydroxy (321.22/303.21/303.21/279.22); C23 - Pentahydroxy (319.21/301.19/301.19/277.19); C27 - Monohydroxy (383.33/159.12); C27 - Dihydroxy (381.32/159.12); C27 - Trihydroxy (379.29/159.12); C27 - Tetrahydroxy (377.28/159.12); C27 - Pentahydroxy (375.27/159.12); Aceto dihydroxy (347.294/329.284); C24 alkylamine - Monohydroxy (359.295/341.284/341.284/359.295); C24 alkylamine - Tetrahydroxy (353.248/335.237/335.237/353.248); C24 alkylamine - Pentahydroxy (351.232/333.221/333.221/351.232); C28 alcohol - 1OH (125.13/143.14); C28 alcohol - 2OH (123.12/141.13); C28 alcohol - 3OH (121.1/139.11) | This specific query/bin was not satisfied; not a universal exclusion of the molecule under a different ion form or cargo. The hydroxylation counts and core classes those rows define | carbon_count_massql_table.md — [TrackSheet], [Mohanty2026-AC] |
| X9 | no 149.0233/167.0339 (phthalate); no 85.0284 (acylcarnitine); no 44.0262 Da repeat series (PEG) (±0.005 Da, ≥1% base) | Those non-bile-acid contaminant classes | nomenclature_glossary.md |

| Ref | Not observed / failed | Rules out | Source |
|---|---|---|---|
| X10 | [M+NH4]+ 689.5798 absent from MS1 scan 3659 | Conditional ranking only: if the target is [M+H]+ and ammonium was present in the mobile phase, blocked-acid candidates are favored and free-acid candidates disfavored. This search does NOT exclude the target itself being [M+NH4]+ or [M+NH4−H2O]+. Mobile-phase ammonium is assumed, not given; no categorical carboxyl assignment follows. — “What we don't see is informative as well.” | dust_region_rules.md §5 / §5a — [Specialist heuristic — Revision-2026-10-12] |
| X11 | No independent corroborating C18:1 cargo product or diagnostic covalent cleavage was established; protonated-fatty-acid ion alone is insufficient. | A specific oleoyl/oleic acid identification and a definite fatty-acyl linkage. | SKILL.md Step 3 ≥2-ion requirement |
| X12 | 108.55597 remains unexplained. 124.99972 matches a phosphorus-containing PC/lysoPC marker outside the CHNOS formula search; other N/S-like minor ions have no coherent cargo assignment. | A clean single-parent spectrum is not established. The isolated phosphorus marker does not identify a phosphatidylcholine or make it the cargo. | nomenclature_glossary.md PC marker screen; formula_table.py element limits |

#### Cargo evidence — what is attached to the bile acid

Every cargo candidate still in play, grouped by the question it answers. Each row cites the C/X rows above that support or argue against it; candidates already ruled out appear only in the exclusionary table. The Section 3 calls and Section 6 open questions are drawn from this table.

| Ref | Candidate | Evidence for | Evidence against | Status | Tier |
|---|---|---|---|---|---|
| **1. Cargo composition and bond** → Specific identity unresolved; C18:1 fatty-acyl unit is a candidate | | | | | |
| A1 | **O-acyl C18:1 ester of a hydroxy-keto C24 bile acid, ammoniated** Neutral C42H70O5; [M+NH4]+ | Formula-compatible ion, C18:1 product and 12-oxo-like core fragments. C7 C8 C9 C6 | One cargo marker only; NH4 adduct and bond are not verified. X11 | Possible | Tentative |
| A2 | **N-acyl C18:1 alkamine bile acid** Neutral C42H73NO5; [M+H]+ | Same ion formula; alkamine screens and C18:1 product are compatible. C2 C3 C7 C9 | No nitrogen-retaining steroid product or diagnostic N-acyl cleavage places the nitrogen on the ring. X11 | Possible | Tentative |
| A3 | **Amide-linked nitrogen-containing C18 unit on a trihydroxy acid** Net C18H33N; free reactant C18H35NO | Alternative formula partition under protonation; blocked C24 acid is favored by conditional ammonium absence. C1 C7 X10 | No two independent ions identify a C18 aminoalcohol or its bond. Cargo formula depends on the assumed core. X11 | Possible | Tentative |

| Ref | Candidate | Evidence for | Evidence against | Status | Tier |
|---|---|---|---|---|---|
| A4 | **Dehydrated ammonium ion of a trihydroxy C24 C18:1 ester**<br>Neutral C42H72O6; [M+NH4−H2O]+ | Same ion formula and oxygenated core products are possible.<br>C1 C7 C8 C9 | No higher ammonium parent or same-run coelution establishes this in-source origin.<br>X11 | Possible | Tentative |
| **2. Ranking by ammonium** → Conditional on the unresolved ion form; does not settle cargo identity | | | | | |
| A5 | **Blocked C24 carboxyl, under [M+H]+ interpretation**<br>No free acidic group assumed | Expected ammonium partner was absent; if protonation is correct this favors a blocked acid over an otherwise comparable free acid.<br>X10 | Target may itself be ammoniated. Mobile-phase ammonium not supplied; no unconditional ranking between A1–A4 is defensible.<br>C11 | Possible | Tentative |

##### Step-7 validation pass

Every peak was re-checked against an elemental formula (C, H, N ≤3, O, S ≤2; even-electron ion, electron mass applied): 78 of 98 fit within ±10 ppm (largest |error| 9.8 ppm; ions used as evidence above: ≤5.5 ppm). 7 peak(s) are $^{13}$C isotope peaks of a stronger ion and are not fragments. No plausible formula: 54.8663, 55.0548, 64.3938, 68.1934, 80.2118, 82.6963, 90.9766, 108.5560, 124.9997, 250.7240, 263.9344, 669.6685, 701.6418 — not used as evidence. 4 ion(s) have a second formula within 5 ppm (see formula_table.tsv) and are named by both where cited. Sulfur near-miss check (§2a): not triggered on the precursor or the main S-bearing ions. Shared CHNO offset: none. $^{34}$S A+2 on the precursor: absent — no peak at the 34S position; weighs against S if the A+2 region was resolved. Core-ion formulas were selected from chemically coherent alternatives: sulfur-rich smallest-error formulas are not automatically accepted. The 337 product fits C24H33O+ at −0.00058 Da (−1.71 ppm), 319 fits C24H31+ at +0.00169 Da (+5.28 ppm), and 339 fits C24H35O+ at −0.00216 Da (−6.37 ppm). The 124.99972 phosphate-like ion is outside the CHNOS search, not proof of no possible formula.

Odd/even parity check (conjugate_fragments.md): 6 even-mass fragment(s) (98.0961, 106.0499, 122.0603, 220.0376, 308.2593, 637.5170) — each carries an odd number of N in its best formula. 8 odd-mass ion(s) carry an even number of N (two nitrogen atoms). Nitrogen-rule violations: 637.5170. Nitrogen in the ion formula does not settle covalent nitrogen versus an ammonium adduct.

#### 5. Why the Automated Tree Stalled at “Trihydroxy_stage1”

The pre-pass reached **Trihydroxy_stage1** (not a leaf): Trihydroxy_stage2 did not match. The unmet conditions:

- **Trihydroxy_stage2**: MS2PROD=X-390.277 resolves to **282.2762** — no peak within 0.01 Da of 282.2762 at >= 2% base peak. (X = precursor 672.5532 as acquired, adduct assumed.) Target − H2O (264.2657) is also absent.

The gate for protonated amidates failed, while the ion form and bond type remain unresolved. Downstream 12-oxo queries are positive standalone tests, not a reached leaf and not an associated published FDR. Reacquire a verified protonated species or compare standards before calling the isomer.

#### 6. Unresolved Aspects

1. **<built-in method title of str object at 0x7f6e66155480>**
2. **<built-in method title of str object at 0x7f6e6623baa0>**
3. **<built-in method title of str object at 0x7f6e662caa60>**
4. **<built-in method title of str object at 0x7f6e662ccbb0>**
5. **<built-in method title of str object at 0x7f6e7f251a70>**
6. **<built-in method title of str object at 0x7f6e66299110>**
7. **<built-in method title of str object at 0x7f6e66293220>**
8. **<built-in method title of str object at 0x8e62cd0>**

9. **<built-in method title of str object at 0x7f6e6629ee90>**
10. **<built-in method title of str object at 0x7f6e66155550>**
11. **<built-in method title of str object at 0x7f6e66148fd0>**
12. **<built-in method title of str object at 0x7f6e66155620>**
13. **<built-in method title of str object at 0x7f6e662ad590>**
14. **<built-in method title of str object at 0x7f6e660fc3d0>**
15. **<built-in method title of str object at 0x7f6e662ad7d0>**
16. **<built-in method title of str object at 0x7f6e6629f210>**
17. **<built-in method title of str object at 0x7f6e66149070>**
18. **<built-in method title of str object at 0x7f6e661556f0>**

#### 7. Species / Taxonomic Context

No biological-source prior was used.

| Part | Finding | Source |
|---|---|---|
| **Host prior** | None available — no host or sample type recognisable in the file name, and none is recorded in the mzML. No prior was applied. | file name |
| **Expected producer** | No producer assigned; host versus microbial origin cannot be inferred from the spectral pattern. | |
| **Plausible route** | Diet not supplied; no source route assigned. | source_and_diet_context.md §3 |

Follow-ups that would test the route:

- Obtain sample/tissue and extraction metadata if biological attribution is needed.

#### 8. Chromatographic Context (Supporting, Not Evidence)

Extracted-ion chromatograms (±5 ppm) over the whole 10.0-min run, for M = 671.54595 ([M+H]+, assumed (ion form not resolved from MS1; Step 0: no form's neutral mass fits a free bile acid or known conjugate within 8 ppm)): All adduct labels in these traces are conditional on the [M+H]+ fallback.

| Trace | Chromatographic peaks (RT min, apex intensity) |
|---|---|
| 672.5532 [M+H]+ (precursor) | 7.45 (1.9e5), 7.76 (2.3e6), 7.91 (4.5e5), 8.16 (9.1e4), 8.42 (2.2e6), 8.59 (5.1e5), 9.07 (1.2e6) ← this scan, 9.17 (3.3e5) ← this scan |
| 689.5798 [M+NH4]+ | 5.41 (4.5e4) |
| 694.5352 [M+Na]+ | 6.77 (1.9e5), 6.87 (1.8e5), 6.98 (1.9e5), 7.19 (1.3e5), 7.36 (1.5e6), 7.49 (7.1e5), 7.71 (2.5e5), 7.91 (5.1e4) |
| 710.5091 [M+K]+ | 7.46 (2.7e5), 7.56 (1.2e5), 7.68 (2.6e5) |
| 654.5427 [M+H-H2O]+ | no peak above 2% of the trace maximum (trace empty) |

Supporting orientation only: retention time alone identifies nothing (stereochemistry_and_caveats.md). 6 other MS2 scan(s) were acquired on this precursor in the run. No isotope/adduct assignment or structure is inferred from these peaks alone.

#### 9. Diagnostic Decision Tree

Each node's accent color is the color of the corresponding ion family highlighted in the Section 2 spectrum. The numbered chips (C = confirmatory, X = exclusionary, A = cargo evidence) are the Ref column of the Section 4 evidence tables, so every node traces back to its ions, mass errors and sources; N chips are Section 11 network analogs, cited only by context (–) nodes.

Dehydrated-core pair · Conjugate / cargo ions · Position indicator · Cluster ratio · Isomer-query ratio · Grouping / precursor arithmetic · Ruled out (no diagnostic ion family)

✓ supports the annotation · ⊘ alternative ruled out (dashed box) · – context only

**Feature 21149 / scan 3663**
Evidence first; unresolved adduct limits interpretation

**Core identity & conjugate**

- **✓ C24 bile-acid core** — Core ladder and abundant ring fragments support class. C1 C4 C8
- **– C18 cargo candidate** — One fatty-acid-like ion; specific cargo unresolved. C9 A1 A2
- **– Adduct unresolved** — Several structures share the same ion formula. C7 C11

**Hydroxylation position**

- **✓ 12-oxo-like pattern** — Strong 309 and passing standalone ratio; tentative. C5 C6
- **– Tree gate failed** — No terminal isomer bin reached. X1
- **– Fingerprint inconclusive** — Nearly equal 159/161; no stereo claim. C10

**Alternatives and context**

- **– Cargo naming not justified** — Missing second independent cargo marker. X11
- **– Conditional ammonium ranking** — Requires correct [M+H]+ interpretation. X10 A5
- **– Distant keto analog** — Network hypothesis only; not an identification. N4

↓

**Modified C24 bile acid with 12-oxo-like fragmentation (Tentative)**

Cargo and adduct unresolved. C18:1-acyl conjugation is a testable hypothesis.

#### 10. Proposed Final Structures

One row per structure the reasoning in Sections 3–9 still supports, drawn with RDKit with the ABCD rings in the standard bile-acid orientation. A partial structure draws only what the evidence claims: wavy bonds mark stereocentres that are not resolved, substituents that are counted but not placed are listed under the drawing and their possible rings shaded, and a highlighted unit is weaker than the rest (the Notes say why). The Tier column is copied from the reasoning. Names are standardized with the metabolite-annotation-cleanup rules (PubChem's title when the exact, stereo-defined structure's record lists the proposed name; otherwise the cleaned label); standardization never changes a tier.

| Structure | Name | Tier | Notes |
|---|---|---|---|
| 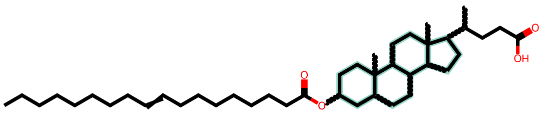<br>+ 1 =O (positions not resolved) | **Candidate O-acyl C18:1 hydroxy-keto C24 acid (illustrative)**<br>partial structure<br>$C_{42}H_{70}O_5$ · 654.52233 Da<br>drawn skeleton: $C_{42}H_{72}O_4$ | Tentative | NH4 adduct hypothesis; example linkage shown at C3, actual site unresolved.<br>Wavy bonds: stereochemistry not resolved at C3, C5, C8, C9, C10, C13, C14, C17, C20<br>1 × =O on rings A–D (shaded); positions not resolved<br>Not resolved:<br>• Ester site, ketone position, chain double-bond position/geometry, all stereochemistry<br>• Whether the precursor is ammoniated and this bond exists<br>Assumed, not shown by this spectrum:<br>• C3 ester placement and unbranched chain with one arbitrarily placed C=C are illustrative, not assignments<br>Name: partial structure (substituent positions not resolved): no PubChem lookup; cleaned label shown |
| 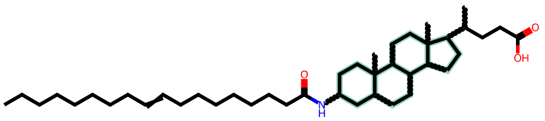<br>+ 2 OH (positions not resolved) | **Candidate N-acyl C18:1 alkamine C24 acid (illustrative)**<br>partial structure<br>$C_{42}H_{73}NO_5$ · 671.54887 Da<br>drawn skeleton: $C_{42}H_{73}NO_3$ | Tentative | Protonated, nitrogen-bearing neutral hypothesis; ring amide placement is illustrative only.<br>Wavy bonds: stereochemistry not resolved at C3, C5, C8, C9, C10, C13, C14, C17, C20<br>2 × OH on rings A–D (shaded); positions not resolved<br>Not resolved:<br>• Nitrogen location and N-acyl bond; OH sites; C=C position/geometry; stereochemistry<br>Assumed, not shown by this spectrum:<br>• C3 amide and straight C18:1 chain are illustrative, not identified<br>Name: partial structure (substituent positions not resolved): no PubChem lookup; cleaned label shown |

#### 11. Network Analogs (graphml; hypotheses about relatedness, not evidence)

Network analogs from HNRC_batch6_EB_101626-78d08e8bc3fe463a950c3976402acb4b-network_singletons.graphml: query node 21149 (m/z 672.5555), component of 80 node(s), 2 direct neighbour(s), 10 library-annotated. Shown: every library-annotated node and every direct neighbour.

| Ref | Node | m/z | Hops | Cosine | Δ query–node (Da) | Candidates (delta tables) | Implication |
|---|---|---|---|---|---|---|---|
| N1 | 1290 (R)-4-((3R,5R,8R,9S,10S,12S,13R,14S,17R)-3,12-dihydroxy-10,… | 373.2734 | 5 (via 15013 → 11041 → 10185 → 2053) | 0.836 (path min) | +299.2821 first candidate +3.1 ppm | (no rationale in the paper's table) (count 16) [Bittremieux2023] ↳ 21149->15013: -0.9845 → O-NH (reverse) [NashDunn2024]; +HN -O: Amidation (count 954) [Bittremieux2023] ↳ 15013->11041: +0.0005 → 63Cu37Cl - 65Cu35Cl difference [NashDunn2024] ↳ 11041->10185: +0.9835 → O-NH [NashDunn2024]; +O -HN: Amide hydroxylation; Deamidation (count 572) [Bittremieux2023] ↳ 10185->2053: +282.2560 → (no rationale in the paper's table) (count 42) [Bittremieux2023]; +C18H34O2 9-octadecenoic acid [Bittremieux2023] ↳ 2053->1290: +17.0266 → +NH3 [M+NH4]+ vs [M+H]+ [skill]; [M+H]+ - [M+NH4]+ difference (or NH3 loss) [NashDunn2024] | query vs node 1290 (5 hops): Δ +299.2821 is a frequent suspect delta (count 16) with no known rationale [Bittremieux2023] — hypothesis |

| Ref | Node | m/z | Hops | Cosine | Δ query–node (Da) | Candidates (delta tables) | Implication |
|---|---|---|---|---|---|---|---|
| N2 | 1329 (R)-4-((3R,5R,8R,9S,10S,12S,13R,14S,17R)-3,12-dihydroxy-10,… | 373.2733 | 5 (via 15013 → 11041 → 10185 → 2053) | 0.836 (path min) | +299.2822 first candidate +3.3 ppm | (no rationale in the paper's table) (count 16) [Bittremieux2023] ↳ 21149->15013: -0.9845 → O-NH (reverse) [NashDunn2024]; +HN -O: Amidation (count 954) [Bittremieux2023] ↳ 15013->11041: +0.0005 → 63Cu37Cl - 65Cu35Cl difference [NashDunn2024] ↳ 11041->10185: +0.9835 → O-NH [NashDunn2024]; +O -HN: Amide hydroxylation; Deamidation (count 572) [Bittremieux2023] ↳ 10185->2053: +282.2560 → (no rationale in the paper's table) (count 42) [Bittremieux2023]; +C18H34O2 9-octadecenoic acid [Bittremieux2023] ↳ 2053->1329: +17.0267 → +NH3 [M+NH4]+ vs [M+H]+ [skill]; [M+H]+ - [M+NH4]+ difference (or NH3 loss) [NashDunn2024] | query vs node 1329 (5 hops): Δ +299.2822 is a frequent suspect delta (count 16) with no known rationale [Bittremieux2023] — hypothesis |

| Ref | Node | m/z | Hops | Cosine | Δ query–node (Da) | Candidates (delta tables) | Implication |
|---|---|---|---|---|---|---|---|
| N3 | 147<br>Spectral Match to 12-Ketodeoxycholic acid from NIST14 [M+H]+ | 391.2838 | 5 (via 15013 → 11041 → 10185 → 2053) | 0.836 (path min) | +281.2717 | no table entry<br>↳ 21149->15013: -0.9845 → O-NH (reverse) [NashDunn2024]; +HN -O: Amidation (count 954) [Bittremieux2023]<br>↳ 15013->11041: +0.0005 → 63Cu37Cl - 65Cu35Cl difference [NashDunn2024]<br>↳ 11041->10185: +0.9835 → O-NH [NashDunn2024]; +O -HN: Amide hydroxylation; Deamidation (count 572) [Bittremieux2023]<br>↳ 10185->2053: +282.2560 → (no rationale in the paper's table) (count 42) [Bittremieux2023]; +C18H34O2 9-octadecenoic acid [Bittremieux2023]<br>↳ 2053->147: -0.9838 → O-NH (reverse) [NashDunn2024]; +HN -O: Amidation (count 954) [Bittremieux2023] | query vs node 147 (5 hops): Δ +281.2717 matches no table entry; any formula arithmetic is a labelled hypothesis |

| Ref | Node | m/z | Hops | Cosine | Δ query–node (Da) | Candidates (delta tables) | Implication |
|---|---|---|---|---|---|---|---|
| N4 | 353<br>12-Ketolithocholic acid [M+NH3+H]+ | 408.3104 | 5 (via 15013 → 11041 → 10185 → 1170) | 0.836 (path min) | +264.2451<br>first candidate +0.1 ppm | (no rationale in the paper's table) (count 180) [Bittremieux2023]<br>↳ 21149->15013: -0.9845 → O-NH (reverse) [NashDunn2024]; +HN -O: Amidation (count 954) [Bittremieux2023]<br>↳ 15013->11041: +0.0005 → 63Cu37Cl - 65Cu35Cl difference [NashDunn2024]<br>↳ 11041->10185: +0.9835 → O-NH [NashDunn2024]; +O -HN: Amide hydroxylation; Deamidation (count 572) [Bittremieux2023]<br>↳ 10185->1170: +184.2886 → no table entry<br>↳ 1170->353: +79.9570 → +SO3 sulfation [skill]; +O3S: O-Sulfonation (count 625) [Bittremieux2023] | query vs node 353 (5 hops): Δ +264.2451 is a frequent suspect delta (count 180) with no known rationale [Bittremieux2023] — hypothesis<br>ion forms differ (query [M+H]+, node [M+NH4]+): the Δ may be an ion-form difference rather than a structural one |

| Ref | Node | m/z | Hops | Cosine | Δ query–node (Da) | Candidates (delta tables) | Implication |
|---|---|---|---|---|---|---|---|
| N5 | 1052 (4R)-4-((3S,5R,9S,10S,13R,14S,17R)-3-hydroxy-10,13-dimethyl… | 373.2735 | 6 (via 15013 → 11041 → 10185 → 1170 → 1099) | 0.836 (path min) | +299.2820 first candidate +3.0 ppm | (no rationale in the paper's table) (count 16) [Bittremieux2023] ↳ 21149->15013: -0.9845 → O-NH (reverse) [NashDunn2024]; +HN -O: Amidation (count 954) [Bittremieux2023] ↳ 15013->11041: +0.0005 → 63Cu37Cl - 65Cu35Cl difference [NashDunn2024] ↳ 11041->10185: +0.9835 → O-NH [NashDunn2024]; +O -HN: Amide hydroxylation; Deamidation (count 572) [Bittremieux2023] ↳ 10185->1170: +184.2886 → no table entry ↳ 1170->1099: +78.9537 → no table entry ↳ 1099->1052: +36.0402 → no table entry | query vs node 1052 (6 hops): Δ +299.2820 is a frequent suspect delta (count 16) with no known rationale [Bittremieux2023] — hypothesis ion forms differ (query [M+H]+, node [M-H2O+H]+): the Δ may be an ion-form difference rather than a structural one |

| Ref | Node | m/z | Hops | Cosine | Δ query–node (Da) | Candidates (delta tables) | Implication |
|---|---|---|---|---|---|---|---|
| N6 | 1275 (4R)-4-((3S,5R,9S,10S,13R,14S,17R)-3-hydroxy-10,13-dimethyl… | 391.2840 | 6 (via 15013 → 11041 → 10185 → 2053 → 1290) | 0.836 (path min) | +281.2715 | no table entry ↳ 21149->15013: -0.9845 → O-NH (reverse) [NashDunn2024]; +HN -O: Amidation (count 954) [Bittremieux2023] ↳ 15013->11041: +0.0005 → 63Cu37Cl - 65Cu35Cl difference [NashDunn2024] ↳ 11041->10185: +0.9835 → O-NH [NashDunn2024]; +O -HN: Amide hydroxylation; Deamidation (count 572) [Bittremieux2023] ↳ 10185->2053: +282.2560 → (no rationale in the paper's table) (count 42) [Bittremieux2023]; +C18H34O2 9-octadecenoic acid [Bittremieux2023] ↳ 2053->1290: +17.0266 → +NH3 [M+NH4]+ vs [M+H]+ [skill]; [M+H]+ - [M+NH4]+ difference (or NH3 loss) [NashDunn2024] ↳ 1290->1275: -18.0106 → -H2O hydration (+H2O) (reverse) [skill]; -H2O H2O (water) (reverse) [NashDunn2024] | query vs node 1275 (6 hops): Δ +281.2715 matches no table entry; any formula arithmetic is a labelled hypothesis |

| Ref | Node | m/z | Hops | Cosine | Δ query–node (Da) | Candidates (delta tables) | Implication |
|---|---|---|---|---|---|---|---|
| N7 | 1624 (4R)-4-((3S,5R,9S,10S,13R,14S,17R)-3-hydroxy-10,13-dimethyl… | 355.2628 | 6 (via 15013 → 11041 → 10185 → 2053 → 147) | 0.836 (path min) | +317.2927 | no table entry ↳ 21149->15013: -0.9845 → O-NH (reverse) [NashDunn2024]; +HN -O: Amidation (count 954) [Bittremieux2023] ↳ 15013->11041: +0.0005 → 63Cu37Cl - 65Cu35Cl difference [NashDunn2024] ↳ 11041->10185: +0.9835 → O-NH [NashDunn2024]; +O -HN: Amide hydroxylation; Deamidation (count 572) [Bittremieux2023] ↳ 10185->2053: +282.2560 → (no rationale in the paper's table) (count 42) [Bittremieux2023]; +C18H34O2 9-octadecenoic acid [Bittremieux2023] ↳ 2053->147: -0.9838 → O-NH (reverse) [NashDunn2024]; +HN -O: Amidation (count 954) [Bittremieux2023] ↳ 147->1624: +36.0210 → H2O+H2O (water) [NashDunn2024]; +H4O2: 2x water (count 1078) [Bittremieux2023] | query vs node 1624 (6 hops): Δ +317.2927 matches no table entry; any formula arithmetic is a labelled hypothesis ion forms differ (query [M+H]+, node [M-2H2O+H]+): the Δ may be an ion-form difference rather than a structural one |

| Ref | Node | m/z | Hops | Cosine | Δ query–node (Da) | Candidates (delta tables) | Implication |
|---|---|---|---|---|---|---|---|
| N8 | 16964<br>3a, 12k [M+H-2H2O]+ | 354.2498 | 6 (via 15013 → 11041 → 10185 → 2053 → 1329) | 0.836 (path min) | +318.3057 | no table entry<br>↳ 21149->15013: -0.9845 → O-NH (reverse) [NashDunn2024]; +HN -O: Amidation (count 954) [Bittremieux2023]<br>↳ 15013->11041: +0.0005 → 63Cu37Cl - 65Cu35Cl difference [NashDunn2024]<br>↳ 11041->10185: +0.9835 → O-NH [NashDunn2024]; +O -HN: Amide hydroxylation; Deamidation (count 572) [Bittremieux2023]<br>↳ 10185->2053: +282.2560 → (no rationale in the paper's table) (count 42) [Bittremieux2023]; +C18H34O2 9-octadecenoic acid [Bittremieux2023]<br>↳ 2053->1329: +17.0267 → +NH3 [M+NH4]+ vs [M+H]+ [skill]; [M+H]+ - [M+NH4]+ difference (or NH3 loss) [NashDunn2024]<br>↳ 1329->16964: +19.0235 → no table entry | query vs node 16964 (6 hops): Δ +318.3057 matches no table entry; any formula arithmetic is a labelled hypothesis<br>ion forms differ (query [M+H]+, node [M-3H2O+H]+): the Δ may be an ion-form difference rather than a structural one |

| Ref | Node | m/z | Hops | Cosine | Δ query–node (Da) | Candidates (delta tables) | Implication |
|---|---|---|---|---|---|---|---|
| N9 | 4176 (4R)-4-((3S,5R,9S,10S,13R,14S,17R)-3-hydroxy-10,13-dimethyl… | 355.2628 | 6 (via 15013 → 11041 → 10185 → 2053 → 1290) | 0.836 (path min) | +317.2927 | no table entry ↳ 21149->15013: -0.9845 → O-NH (reverse) [NashDunn2024]; +HN -O: Amidation (count 954) [Bittremieux2023] ↳ 15013->11041: +0.0005 → 63Cu37Cl - 65Cu35Cl difference [NashDunn2024] ↳ 11041->10185: +0.9835 → O-NH [NashDunn2024]; +O -HN: Amide hydroxylation; Deamidation (count 572) [Bittremieux2023] ↳ 10185->2053: +282.2560 → (no rationale in the paper's table) (count 42) [Bittremieux2023]; +C18H34O2 9-octadecenoic acid [Bittremieux2023] ↳ 2053->1290: +17.0266 → +NH3 [M+NH4]+ vs [M+H]+ [skill]; [M+H]+ - [M+NH4]+ difference (or NH3 loss) [NashDunn2024] ↳ 1290->4176: +18.0106 → hydration (+H2O) [skill]; +H2O H2O (water) [NashDunn2024] | query vs node 4176 (6 hops): Δ +317.2927 matches no table entry; any formula arithmetic is a labelled hypothesis ion forms differ (query [M+H]+, node [M-H2O+H]+): the Δ may be an ion-form difference rather than a structural one |

| Ref | Node | m/z | Hops | Cosine | Δ query–node (Da) | Candidates (delta tables) | Implication |
|---|---|---|---|---|---|---|---|
| N10 | 31090<br>12-Ketolithocholic acid [M+NH3+H]+ | 407.2423 | 8 (via 15013 → 11041 → 10185 → 1170 → 1099 → 22720 → 10391) | 0.836 (path min) | +265.3132 | no table entry<br>↳ 21149->15013: -0.9845 → O-NH (reverse) [NashDunn2024]; +HN -O: Amidation (count 954) [Bittremieux2023]<br>↳ 15013->11041: +0.0005 → 63Cu37Cl - 65Cu35Cl difference [NashDunn2024]<br>↳ 11041->10185: +0.9835 → O-NH [NashDunn2024]; +O -HN: Amide hydroxylation; Deamidation (count 572) [Bittremieux2023]<br>↳ 10185->1170: +184.2886 → no table entry<br>↳ 1170->1099: +78.9537 → no table entry<br>↳ 1099->22720: +1.1931 → no table entry<br>↳ 22720->10391: +0.8401 → no table entry<br>↳ 10391->31090: +0.0382 → -O +CH4 oxidation (reverse) [Bittremieux2023] | query vs node 31090 (8 hops): Δ +265.3132 matches no table entry; any formula arithmetic is a labelled hypothesis |
| N11 | 15013<br>unannotated | 673.5400 | 1 | 0.836 | -0.9845<br>first candidate -0.7 ppm | O-NH (reverse) [NashDunn2024]<br>+HN -O: Amidation (count 954) [Bittremieux2023]<br>-O +HN amide hydroxylation (reverse) [Bittremieux2023] | query = unannotated node 15013 -0.9845 (O-NH) — hypothesis |

| Ref | Node | m/z | Hops | Cosine | Δ query–node (Da) | Candidates (delta tables) | Implication |
|---|---|---|---|---|---|---|---|
| N12 | 17305 unannotated | 672.4889 | 1 | 0.833 | +0.0666 | no table entry | query vs unannotated node 17305: Δ +0.0666 matches no table entry; any formula arithmetic is a labelled hypothesis |

Direct neighbors N11 and N12 have cosine scores 0.836 and 0.833 and query-minus-node differences −0.9845 and +0.0666 Da, respectively. Neither has a library annotation; the former supports only an O/NH or ion-form hypothesis, while the latter has no named modification. The distant 12-ketolithocholic-acid-labeled analog N4 is five hops away and differs by +264.2451 Da, matching $C_{18}H_{32}O$; this is compatible with an acylation hypothesis, not proof of identity. Other distant oxo-bile-acid annotations N3 , N5 –N9 are consistent with testing a keto-core reading. Hypothesis searches matched $C_{18}H_{32}O$ at N4 , $C_{18}H_{34}O_2$ at N26 / N32 / N66 , and $C_{18}H_{35}NO$ at N3 / N6 ; they do not locate nitrogen or establish a covalent cargo. No $C_{18}H_{33}N$ difference matched. Zero-mass hop labels involving copper/chloride are generic table coincidences and are not evidence for metals or halogens.

Δ = query m/z − node m/z, both read from the graphml nodes (positive: the query is heavier). Edge-stored deltas are a check only; their sign is not trusted. Hypotheses about relatedness, not evidence: a neighbour's library name is not an annotation of the query, and no row raises a tier (cap tentative). Graphml fields read: cosine ← cosine_score, edge_type ← EdgeType, mass_difference ← mass_difference, scan1 ← scan1, scan2 ← scan2, best_ion ← Best Ion, component ← component, ion_mode ← IonMode, library_mass_diff ← MassDiff, library_name ← Compound_Name, library_score ← MQScore, library_shared_peaks ← SharedPeaks, library_smiles ← Smiles, library_spectrum_id ← SpectrumID, mz ← mz, neutral_mass ← neutral M mass, rt_min ← rt_min, rt_seconds ← RTMean. Edge-stored deltas (240 component edges): 10 disagree in magnitude with the node m/z and 4 carry the opposite sign to m/z(source) − m/z(target); node m/z was used throughout. 67 further node(s) of the component are listed in prepass.md / prepass.json.

---

*Generated by the bile-acid-msms-annotation-v5-6 skill (deterministic MassQL pre-pass, pure evaluator) plus literature-grounded reasoning in the referenced skill files. Source: P3_C5_Sample_x3160658.mzML, scan 3663. Positions in candidate drawings are illustrative; exact identity is unresolved. Run time: 5 min 3 s, start to finish (started 2026-10-27 20:26:06 UTC, report rendered 2026-10-27 20:31:09 UTC).*
