## Supplementary File 3 for "Capturing human expertise for AI-assisted molecular structure annotation"

### Bile-Acid MS/MS Annotation Report

P1_G7_Sample_X3148665.mzML — Scan 2595 | Generated with the bile-acid-msms-annotation-v5-6 skill

Run time: 27 min 8 s, start to finish (started 2026-10-27 17:04:00 UTC, report rendered 2026-10-27 17:31:08 UTC)

#### 1. Precursor / Input Summary

| Field | Value | How obtained |
|---|---|---|
| Precursor m/z | 546.41502 | mzML scan header |
| Ion form | [M+H]+, z = 1 (measured, not assumed) | MS1 scan 2592; isotope spacing 1.0033 Da; corroborated by [M+Na]+ 568.39679, [M+K]+ 584.37146, [M+H-H2O]+ 528.40002 in the same MS1 |
| Neutral monoisotopic mass | 545.40774 | from the adduct assignment |
| Neutral formula candidates (±5 ppm, C/H/N≤3/O/S≤2) | C33H55NO5 (-0.52 ppm); also within 5 ppm: C34H59NS2 (-2.11 ppm); C37H55NS (+4.07 ppm) | formula_table.py (mass alone does not choose among these) |
| Retention time | 6.439 min (10.0 min total run) | mzML |
| Polarity / collision energy | positive ESI / HCD 30 | mzML; filter: FTMS + p ESI d Full ms2 546.4150@hcd30.00 [50.0000-575.0000] |
| Instrument | Q Exactive (Xcalibur 2.9-290033/2.9.0.2926; pwiz 3.0.23118) | mzML header |
| Isolation window | 546.415 –1.5/+1.5 Da; 6 MS1 ion(s) ≥1% inside it | mzML; MS1 scan 2592 |
| Peaks in MS2 | 199; base peak 154.1225 | mzML |
| Other MS2 on this precursor (whole run) | scan 2327 @ 5.768 min, scan 2469 @ 6.121 min | extract_context.py |
| Engine cross-check | not run (pure evaluator only; no borderline ratio) | --engine pure (default) |

Collision energy 30 is below the NCE ~44–45 at which the dust-region rules were characterised. The dust region was still evaluated; expect fewer and weaker dust ions and some ratio shift. Lower energy alone does not lower a tier (stereochemistry_and_caveats.md §4).

#### 2. Annotated MS/MS Spectrum

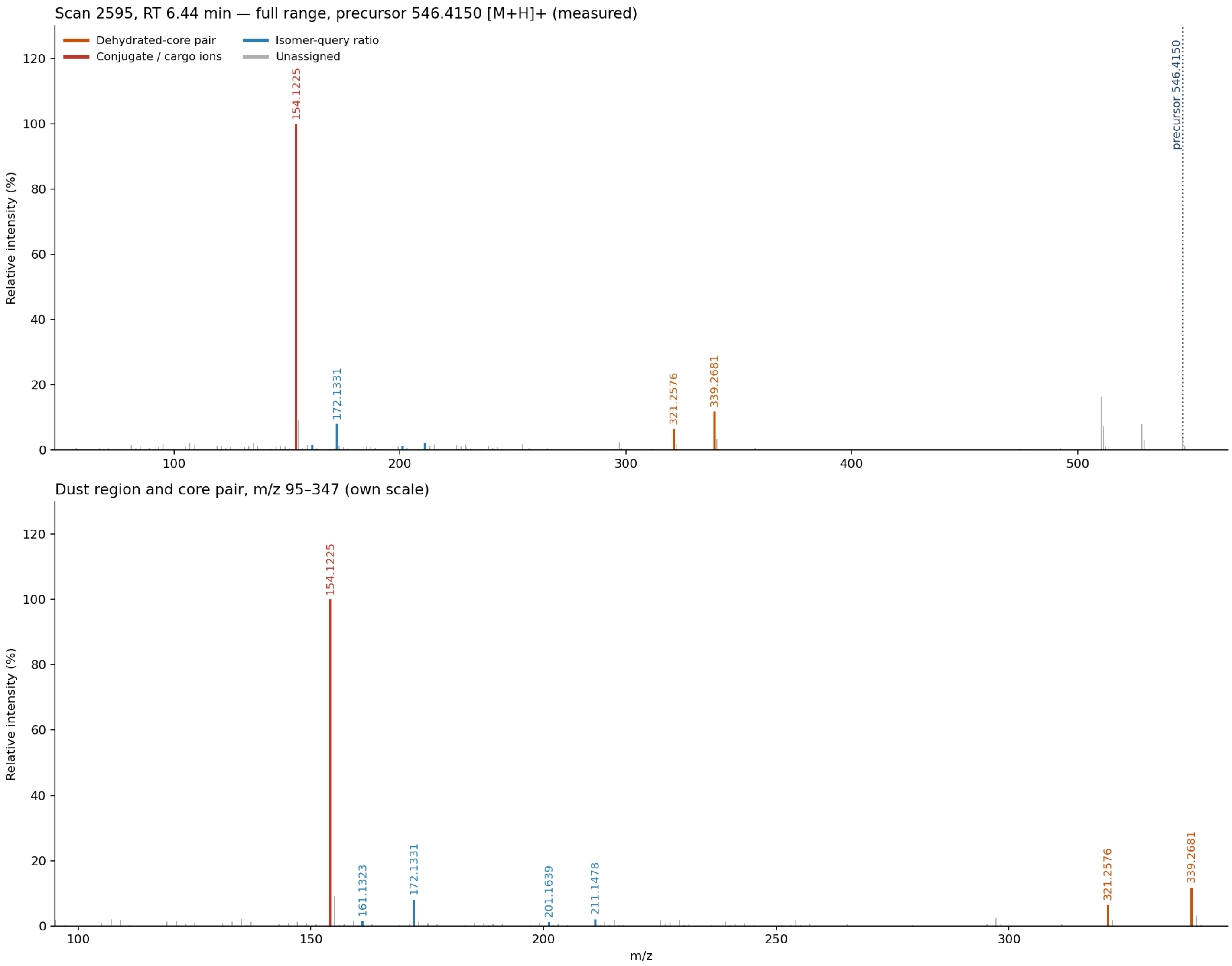


Scan 2595 (RT 6.44 min, precursor 546.4150 [M+H]+). Top: full range. Bottom: m/z 95–347 on its own scale, where the cargo ions and the ring-derived “dust” sit. Colors follow the shared palette; gray stems are unassigned peaks.

#### 3. Proposed Annotation

C24oic(OH)2–C9H17NO2 — C24 dihydroxy bile-acid amidate with an unlisted C9H17NO2 cargo — i.e. **C33H55NO5 C24 dihydroxy bile-acid amidate** formula-level cargo; 3,12α versus 7,12α and full stereochemistry unresolved

| Aspect | Call | Tier |
| --- | --- | --- |
| Bile acid (not bile alcohol or screened contaminant class) | Yes | **Confirmed** |
| Core carbon count | C24 | **Confirmed** |
| Hydroxylation count | 2 (true dihydroxy favoured; no monoketone signature) | **Probable** |

| Aspect | Call | Tier |
|---|---|---|
| Cargo composition | C9H17NO2; exact constitutional identity unresolved — see A1 – A3 | Probable |
| Bond to the bile acid | C-24 carboxamide favoured — see A4 | Probable |
| Hydroxyl-position bin | 3,12α or 7,12α (published intermediate bin; FDR 0.18) | Confirmed |
| Specific isomer and remaining α/β stereochemistry | — | Unresolved |

C24 standard cholanic-acid skeleton (core_structure_reference.md family 2), with two ring hydroxyls and an N-containing cargo amidated at the side-chain carboxyl. One hydroxyl is in the 12α branch; the second is at C3 or C7. The cargo is fixed only to C9H17NO2.

#### 4. Evidence Table

##### Confirmatory (positive) evidence

| Ref | Obs. m/z | %base | Assignment | Calc. | Δppm | Interpretation | Source |
|---|---|---|---|---|---|---|---|
| C1 | 339.2681<br>321.2576 | 11.78<br>6.34 | C24H35O+<br>C24H33+ | 339.2682<br>321.2577 | -0.5<br>-0.2 | Core-class query **C24 - Dihydroxy** passed: dehydrated-core pair present (query targets 339.27, 321.26, ±0.01 Da, ≥1% each — floor adapted from the published 5%); the tree's **Dihydroxy_stage1** query passes on the same two ions — “the more hydroxyls, the smaller the number” | carbon_count_massql_table.md — [TrackSheet], [Mohanty2026-AC]; floor 5 % → 1 % [Adaptation-2026-10-18]; scripts/rules/massql_queries.yaml — [Mohanty2026-AC], [Transcript-UCSD9] |

| Ref | Obs. m/z | %base | Assignment | Calc. | Δppm | Interpretation | Source |
|---|---|---|---|---|---|---|---|
| C2 | 23 ring ions | ≥1 | odd-mass C/H/O | — | max \|9.2\| | Dust region m/z 100–250: 23 ring-derived ions (need 7) → passes: 107.0854 ($C_8H_{11}^+$), 109.1011 ($C_8H_{13}^+$), 119.0853 ($C_9H_{11}^+$), 121.1012 ($C_9H_{13}^+$), 133.1012 ($C_{10}H_{13}^+$), 135.1167 ($C_{10}H_{15}^+$), 137.0960 ($C_9H_{13}O^+$), 145.1010 ($C_{11}H_{13}^+$), 147.1168 ($C_{11}H_{15}^+$), 159.1168 ($C_{12}H_{15}^+$), 161.1323 ($C_{12}H_{17}^+$), 173.1341 ($C_{13}H_{17}^+$), 185.1323 ($C_{14}H_{17}^+$), 187.1480 ($C_{14}H_{19}^+$), 199.1477 ($C_{15}H_{19}^+$), 201.1639 ($C_{15}H_{21}^+$), 211.1478 ($C_{16}H_{19}^+$), 213.1638 ($C_{16}H_{21}^+$), 215.1792 ($C_{16}H_{23}^+$), 225.1634 ($C_{17}H_{21}^+$), 227.1793 ($C_{17}H_{23}^+$), 229.1590 ($C_{16}H_{21}O^+$), 239.1792 ($C_{18}H_{23}^+$). Set aside as N/S-bearing (cargo-like): 154.1225, 172.1331 — “the dust tells you where the things are attached. It doesn't tell you... if it was alpha or beta.” | dust_region_rules.md — [Specialist heuristic — Revision-2026-09-21], [Transcript-UCSDxx] |
| C3 | 546.4150 | MS1 | [M+H]+, z = 1 | — | — | Ion form measured from MS1 scan 2592: isotope spacing 1.0033 Da; [M+Na]+ 568.3968 (expected 568.3970), [M+K]+ 584.3715 (expected 584.3709), [M+H-H2O]+ 528.4000 (expected 528.4044) in the same scan. Confidence: probable. | ms1_context.assess_ion_form (SKILL.md Input handling, step 2) |

| Ref | Obs. m/z | %base | Assignment | Calc. | Δppm | Interpretation | Source |
|---|---|---|---|---|---|---|---|
| C4 | 339.2681<br>321.2576<br>172.1331 | 11.78<br>6.34<br>8.03 | C24H35O+<br>C24H33+<br>C9H18NO2+ | 339.2682<br>321.2577<br>172.1332 | -0.5<br>-0.2<br>-0.8 | Dihydroxy_stage2 query PASSED (Amidated dihydroxy): presence | scripts/rules/massql_queries.yaml — [Mohanty2026-AC] |
| C5 | 161.1323<br>201.1639 | 1.53<br>1.12 | C12H17+<br>C15H21+ | 161.1325<br>201.1638 | -1.0<br>+0.4 | Di-3,6-OH query PASSED (3,6-OH2): obs 0.731× vs required 0.7× ±60% (0.28–1.12) — “keno always has a nice 201” | scripts/rules/massql_queries.yaml — [Mohanty2026-AC], [Notebook p.14], [Transcript-STE0xx] |
| C6 | 211.1478<br>201.1639 | 2.05<br>1.12 | C16H19+<br>C15H21+ | 211.1481<br>201.1638 | -1.6<br>+0.4 | Di-3,12a-OH\|Di-7,12a-OH query PASSED (3,12a-OH2; 7,12a-OH2): obs 0.545× vs required 0.6× ±80% (0.12–1.08) — “a big 201 would mean it's a 3,7 [substitution]... a small 201 is dihydroxy [3,12, i.e. missing the 7-position].” | scripts/rules/massql_queries.yaml — [Mohanty2026-AC], [Specialist heuristic] |
| C7 | 545.40774 (neutral) | — | C33H55NO5 | 545.40802 | -0.5 | Exact-mass fit of the neutral from [M+H]+ (measured by MS1 ion-form assessment). The C24H40O4 dihydroxy-acid core plus C9H17NO2 cargo minus H2O gives C33H55NO5 exactly, and the 172.1331/154.1225 cargo-ion pair independently carries C9H17NO2. The alternative S-bearing precursor formulas lack corresponding fragment support, and the precursor 34S A+2 line is absent. — “close is not close enough... it has to be exact.” | precursor arithmetic (formula_table.py), [Specialist heuristic] |

| Ref | Obs. m/z | %base | Assignment | Calc. | Δppm | Interpretation | Source |
|---|---|---|---|---|---|---|---|
| C8 | 172.1331<br>154.1225 | 8.03<br>100.00 | C9H18NO2+ ([X+H]+)<br>C9H16NO+ ([X+H-H2O]+) | 172.1332<br>154.1226 | -0.8<br>-0.7 | Two independent cargo ions establish neutral X = C9H17NO2: its protonated ion and its water-loss product. This formula is not a named cargo in bile_acid_deltas.tsv; delta lookup finds mass-equivalent peptidomic labels, but those labels are hypotheses rather than identities. | formula_table.py; conjugate_fragments.md §Cargos not in this table; [Bittremieux 2023] hypothesis-grade delta labels |
| C9 | 154.1225 | 100.00 | precursor – neutral C24H40O4 bile acid | 154.1224 | +0.9 | The base peak equals precursor m/z minus the neutral dihydroxy C24 bile-acid mass, satisfying the formal amidate-confirmation arithmetic and favouring an N-linked C-24 carboxamide. The exact constitution of the C9H17NO2 residue is not established. | conjugate_fragments.md §Amino-acid and amine conjugates; [Mohanty2026-AC] |

#### Exclusionary (negative) evidence — what this rules out

| Ref | Not observed / failed | Rules out | Source |
|---|---|---|---|
| X1 | Di-1-ketone failed: no peak within 20 ppm of 161.1320 at >= 95% base peak | The Di-1-ketone bin (1 ketone) — below the node where the tree stopped, so this is the query's own result, not a tree step | scripts/rules/massql_queries.yaml — [Mohanty2026-AC] |
| X2 | Di-3,12a-OH failed: 239.179: obs 1.570× vs required 1× ±30% (0.7–1.3) | The Di-3,12a-OH bin (3,12a-OH2) | scripts/rules/massql_queries.yaml — [Mohanty2026-AC] |
| X3 | Di-7,12a-OH failed: 239.179: obs 1.570× vs required 0.25× ±30% (0.175–0.325) | The Di-7,12a-OH bin (7,12a-OH2) | scripts/rules/massql_queries.yaml — [Mohanty2026-AC] |

| Ref | Not observed / failed | Rules out | Source |
|---|---|---|---|
| X4 | Di-3,7-OH\|Di-3,6-OH\|Di-3,12b-OH\|Di-7,12b-OH failed: 201.163: obs 0.545× vs required 4.2× ±40% (2.52–5.88) | The Di-3,7-OH\|Di-3,6-OH\|Di-3,12b-OH\|Di-7,12b-OH bin (3,7-OH2; 3,6-OH2; 3,12b-OH2; 7,12b-OH2) — below the node where the tree stopped, so this is the query's own result, not a tree step | scripts/rules/massql_queries.yaml — [Mohanty2026-AC] |
| X5 | Di-3,7-OH\|Di-3,12b-OH\|Di-7,12b-OH failed: 201.163: obs 0.731× vs required 2× ±20% (1.6–2.4) | The Di-3,7-OH\|Di-3,12b-OH\|Di-7,12b-OH bin (3,7-OH2; 3,12b-OH2; 7,12b-OH2) — below the node where the tree stopped, so this is the query's own result, not a tree step | scripts/rules/massql_queries.yaml — [Mohanty2026-AC] |
| X6 | Di-3,7-OH failed: 173.132: obs 1.395× vs required 0.5× ±30% (0.35–0.65) | The Di-3,7-OH bin (3,7-OH2) — below the node where the tree stopped, so this is the query's own result, not a tree step | scripts/rules/massql_queries.yaml — [Mohanty2026-AC] |
| X7 | Di-3,12b-OH\|Di-7,12b-OH failed: 173.132: obs 1.395× vs required 1× ±20% (0.8–1.2) | The Di-3,12b-OH\|Di-7,12b-OH bin (3,12b-OH2; 7,12b-OH2) — below the node where the tree stopped, so this is the query's own result, not a tree step | scripts/rules/massql_queries.yaml — [Mohanty2026-AC] |

| Ref | Not observed / failed | Rules out | Source |
|---|---|---|---|
| X8 | Core-class rows absent: C24 - Nonhydroxy (343.3/325.29); C24 - Monohydroxy (341.28/323.27); C24 - Trihydroxy (337.25/319.24); C24 - Tetrahydroxy (335.24/317.23); C24 - Pentahydroxy (333.22/315.21); C23 - Monohydroxy (327.27/309.26/309.26/285.25); C23 - Dihydroxy (325.25/307.24/307.24/283.24); C23 - Trihydroxy (323.24/305.23/305.23/281.23); C23 - Tetrahydroxy (321.22/303.21/303.21/279.22); C23 - Pentahydroxy (319.21/301.19/301.19/277.19); C27 - Monohydroxy (383.33/159.12); C27 - Dihydroxy (381.32/159.12); C27 - Trihydroxy (379.29/159.12); C27 - Tetrahydroxy (377.28/159.12); C27 - Pentahydroxy (375.27/159.12); Aceto dihydroxy (347.294/329.284); C24 alkylamine - Monohydroxy (359.295/341.284/341.284/359.295); C24 alkylamine - Dihydroxy (357.279/339.268/339.268/357.279); C24 alkylamine - Trihydroxy (355.263/337.253/337.253/355.263); C24 alkylamine - Tetrahydroxy (353.248/335.237/335.237/353.248); C24 alkylamine - Pentahydroxy (351.232/333.221/333.221/351.232); C28 alcohol - 1OH (125.13/143.14); C28 alcohol - 2OH (123.12/141.13); C28 alcohol - 3OH (121.1/139.11) | The hydroxylation counts and core classes those rows define | carbon_count_massql_table.md — [TrackSheet], [Mohanty2026-AC] |
| X9 | no 86.0964/104.1070/124.9998/184.0733 (PC / lysoPC head group); no 149.0233/167.0339 (phthalate); no 85.0284 (acylcarnitine); no 44.0262 Da repeat series (PEG) (±0.005 Da, ≥1% base) | Those non-bile-acid contaminant classes | nomenclature_glossary.md |
| X10 | [M+NH4]+ 563.4416 absent from MS1 scan 2592, although [M+Na]+ 568.3968, [M+K]+ 584.3715, [M+H-H2O]+ 528.4000 were found in it (the MS1 could have shown the adduct) | Candidates with a free acidic group (free –COOH, –SO3H …) are disfavoured; blocked candidates (esters, amides, lactones) favoured. Tentative; names no substituent. Caveat: mobile-phase ammonium not stated: assumed present (most acetonitrile-containing buffers carry some) -- say so in the report; if the mobile phase in fact lacks ammonium, this absence means nothing. — “What we don't see is informative as well.” | dust_region_rules.md §5 / §5a — [Specialist heuristic — Revision-2026-10-12] |

#### Cargo evidence — what is attached to the bile acid

Every cargo candidate still in play, grouped by the question it answers. Each row cites the C/X rows above that support or argue against it; candidates already ruled out appear only in the exclusionary table. The Section 3 calls and Section 6 open questions are drawn from this table.

| Ref | Candidate | Evidence for | Evidence against | Status | Tier |
|---|---|---|---|---|---|
| **1. Cargo composition** → C9H17NO2 (probable); identity unresolved | | | | | |
| A1 | **Unlisted C9H17NO2 cargo** C9H17NO2; neutral cargo before condensation | The protonated cargo at 172.1331 and its water-loss ion at 154.1225 both fit within 1 ppm; whole-molecule arithmetic gives C33H55NO5. C7 C8 | — | Supported | Probable |
| **2. Constitutional identity of C9H17NO2** → Blocked/non-acidic constitution favoured; exact identity unresolved | | | | | |
| A2 | **Unsaturated or cyclic amino-acid-like isomer with a free COOH** C9H17NO2; at least one free acidic group in the intact conjugate | The formula and nitrogen parity are compatible, but no isomer-specific fragment was found. C8 | Disfavoured because an informative ammonium-adduct search was negative; the mobile-phase ammonium content was assumed rather than supplied. X10 | Disfavoured | Tentative |
| A3 | **Blocked or non-acidic C9H17NO2 constitutional isomer** C9H17NO2; no free COOH/SO3H/phosphate | Favoured by the informative absence of [M+NH4]+; the 172/154 pair fixes formula but not functional-group layout. C8 X10 | — | Favoured | Tentative |
| **3. Bond between the bile acid and cargo** → C-24 carboxamide favoured (probable) | | | | | |
| A4 | **Amide at the bile-acid side-chain carboxyl** C24H40O4 + C9H17NO2 − H2O | The base peak is the precursor-minus-free-bile-acid amidate fragment, while 172.1331 recovers protonated C9H17NO2. C8 C9 | — | Favoured | Probable |

#### Step-7 validation pass

Every peak was re-checked against an elemental formula (C, H, N ≤3, O, S ≤2; even-electron ion, electron mass applied): 123 of 199 fit within ±10 ppm (largest |error| 9.2 ppm; ions used as evidence above: ≤1.6 ppm). 48 peak(s) are $^{13}C$ isotope peaks of a stronger ion and are not fragments. No plausible formula: 52.3312, 52.4200, 56.7075, 56.7106, 56.8221, 57.5703, 58.7086, 58.7118, 58.8231, 60.7127, 60.8244, 62.3269, 63.6551, 64.1907, 65.7182, 66.8669, 73.2566, 135.1069, 150.3392, 154.1428,

154.1484, 154.1563, 154.1713, 254.2000, 296.2478, 312.2813, 479.3102, 546.4781 — not used as evidence. 2 ion(s) have a second formula within 5 ppm (see formula_table.tsv) and are named by both where cited. Sulfur near-miss check (§2a): not triggered on the precursor or the main S-bearing ions. Shared CHNO offset: none. $^{34}$S A+2 on the precursor: absent — no peak at the 34S position; weighs against S if the A+2 region was resolved.

Odd/even parity check (conjugate_fragments.md): 14 even-mass fragment(s) (154.1225, 172.1331, 208.1314, 236.1640, 254.1748, 348.2539, 374.2681, 474.3734, 492.3834, 510.2366, 510.3938, 528.4042, 546.4145, 548.4181) — each carries an odd number of N in its best formula. No odd-mass ion carries an even number of nitrogen atoms. The nitrogen rule holds for every assigned ion.

#### 5. Why the Automated Tree Stalled at “Di-3,12a-OH|Di-7,12a-OH”

The pre-pass reached **Di-3,12a-OH|Di-7,12a-OH** (not a leaf): Di-3,12a-OH, Di-7,12a-OH did not match. The unmet conditions:

- **Di-3,12a-OH**: MS2MZ=239.179 resolves to **239.1790** — intensity 4.17e+04 outside required 1.859e+04-3.453e+04 (= Y*1 +/- 30%). Target – H2O (221.1684) is also absent.
- **Di-7,12a-OH**: MS2MZ=239.179 resolves to **239.1790** — intensity 4.17e+04 outside required 4648-8633 (= Y*0.25 +/- 30%). Target – H2O (221.1684) is also absent.

The stall is a real ratio-window failure, not a precursor-relative mass shift: both child rules compare the observed 239/243 dust-ion ratio, and m/z 239.1790 is above both allowed windows. The published intermediate 3,12α|7,12α bin remains valid (FDR 0.18), but no single positional isomer is called. HCD 30 is below the approximately 44–45 setting used to characterize the rules; reacquisition near NCE 45 and standards for both child isomers would test whether the unusually high 239 signal is energy-dependent.

#### 6. Unresolved Aspects

1. **Exact cargo identity cannot be determined.** The spectrum fixes C9H17NO2 through two independent ions C8 , but this formula has multiple constitutional isomers. Delta tables attach hypothesis-grade peptidomic labels (including N-desmethyldolaisoleuine and N-methyl homo-vinylogous valine mass equivalents), without diagnostic fragments or a network analog to support either name. An authentic standard, MS3 of m/z 172/154, or NMR would be needed. See A1 – A3 .
2. **C3 versus C7 placement remains unresolved.** The published tree reaches the 3,12α|7,12α intermediate bin (FDR 0.18) but neither terminal child passes: m/z 239.1790 is too intense for both child windows. Reacquisition near NCE 45 and comparison with authentic 3,12α and 7,12α standards would be the cleanest test.
3. **The low-energy dust shape conflicts with a simple named-isomer reading.** At HCD 30, m/z 159.1168 and 161.1323 are nearly equal, a provisional 3α,7α-like fingerprint, whereas the paper-validated 201/211 branch excludes the 3,7 terminal path C5 – C6 , X4 – X7 . The heuristic fingerprint does not override the published tree.
4. **Full alpha/beta stereochemistry cannot be assigned.** Apart from the tree's 12α branch, MS/MS alone does not establish the remaining hydroxyl stereochemistry or C5 ring fusion. NMR or co-injection with stereochemically defined standards is required (stereochemistry_and_caveats.md §1).
5. **Ammonium-adduct ranking depends on an unstated mobile phase.** The LC solvent was not supplied, so ammonium availability was assumed. The absent adduct therefore disfavours, rather than rules out, free-acid cargo isomers X10 . If the mobile phase lacked ammonium, that ranking is uninformative.
6. **No same-run RT reference or network analog is available.** This filename is absent from the internal RT-order reference. Feature 1614 is a singleton in the supplied molecular network, so neither retention-order corroboration nor propagation from a connected library seed is possible.

#### 7. Species / Taxonomic Context

Contextual prior only; no host, tissue, diet, or treatment metadata were inferable from the mzML filename, so none influenced the spectral call.

| Part | Finding | Source |
|---|---|---|
| **Host prior** | None available — no host or sample type recognisable in the file name, and none is recorded in the mzML. No prior was applied. | file name |
| **Expected producer** | Not assessed. The formula-level C9H17NO2 cargo is not tied to a documented host clade by the evidence available here. | source_and_diet_context.md §3 |
| **Plausible route** | Diet and exposure history were not provided; origin remains open (host, microbial, dietary, or xenobiotic). | source_and_diet_context.md §3 |

Follow-ups that would test the route:

- Recover the study metadata for sample P1_G7_Sample_X3148665 (host, tissue/biofluid, diet, treatment and collection conditions).
- Check whether the same m/z/RT/MS2 feature occurs reproducibly in biologically related samples and blanks.
- Acquire MS3 of m/z 172.1331 and 154.1225 and compare with candidate C9H17NO2 standards.

#### 8. Chromatographic Context (Supporting, Not Evidence)

Extracted-ion chromatograms (±5 ppm) over the whole 10.0-min run, for M = 545.40774 ([M+H]+, measured (MS1 ion-form assessment)):

| Trace | Chromatographic peaks (RT min, apex intensity) |
|---|---|
| 546.4150 [M+H]+ (precursor) | 4.99 (9.2e5), 5.33 (9.9e5), 5.99 (1.9e6), 6.13 (6.8e6), 6.45 (2.3e7) ⟵ this scan |
| 563.4416 [M+NH4]+ | 6.13 (5.1e4) |
| 568.3970 [M+Na]+ | 5.97 (2.8e5), 6.14 (2.9e6), 6.45 (1.0e7) ⟵ this scan, 6.66 (2.2e5) |
| 584.3709 [M+K]+ | 6.14 (1.6e5), 6.45 (4.8e5) ⟵ this scan |
| 528.4044 [M+H-H2O]+ | 6.14 (1.2e6), 6.46 (4.5e5) ⟵ this scan, 7.34 (1.2e7) |

Supporting orientation only: retention time alone identifies nothing (stereochemistry_and_caveats.md). 2 other MS2 scan(s) were acquired on this precursor in the run.

#### 9. Diagnostic Decision Tree

Each node's accent color is the color of the corresponding ion family highlighted in the Section 2 spectrum. The numbered chips (C = confirmatory, X = exclusionary, A = cargo evidence) are the Ref column of the Section 4 evidence tables, so every node traces back to its ions, mass errors and sources.

Dehydrated-core pair · Conjugate / cargo ions · Isomer-query ratio · Grouping / precursor arithmetic · Ruled out (no diagnostic ion family)

✓ supports the annotation · ⊘ alternative ruled out (dashed box) · – context only

**MS2 scan 2595**
precursor 546.4150 [M+H]+ · 199 peaks · base peak 154.1225

- **Core identity & conjugate**
  carbon count, hydroxyl count, cargo and linkage
  - ✓ **C24 dihydroxy core pair**
    339.2681 / 321.2576; dust passes with 23 ring ions
    C1 C2
  - ✓ **Neutral formula $C_{33}H_{55}NO_5$**
    -0.5 ppm; core + cargo condensation arithmetic agrees
    C7
  - ✓ **$C_9H_{17}NO_2$ cargo**
    172.1331 and 154.1225; exact identity unresolved
    C8 A1
  - ✓ **C-24 carboxamide**
    formal precursor-minus-free-acid amidate fragment
    C9 A4
- **Hydroxylation position**
  published tree reaches an intermediate ambiguity bin
  - ✓ **3,12α | 7,12α bin**
    201/211 = 0.545; published intermediate-bin FDR 0.18
    C6
  - ✗ **3,12α terminal child**
    239/243 ratio above its allowed window
    X2
  - ✗ **7,12α terminal child**
    239/243 ratio above its allowed window
    X3
- **Alternatives ruled out or disfavoured**
  negative evidence and consistency checks
  - ⊘ **Monoketone bin**
    161.1323 is only 1.53% of base, not >95%
    X1
  - ⊘ **Other C24 OH counts and core classes**
    their diagnostic rows do not pass
    X8
  - ⊘ **PC, PEG, phthalate, acylcarnitine**
    class-specific contaminant ions absent
    X9
  - ✗ **Free-acid cargo isomers**
    disfavoured by absent ammonium adduct; mobile phase assumed
    X10 A2 A3

↓

**C24 dihydroxy bile-acid $C_9H_{17}NO_2$ amidate (Tentative)**

core and formula are strong; exact cargo constitution and 3,12α versus 7,12α identity remain unresolved

#### 10. Proposed Final Structures

One row per structure the reasoning in Sections 3–9 still supports, drawn with RDKit with the ABCD rings in the standard bile-acid orientation. A partial structure draws only what the evidence claims: wavy bonds mark stereocentres that are not resolved, substituents that are counted but not placed are listed under the drawing and their possible rings shaded, and a highlighted unit is weaker than the rest (the Notes say why). The Tier column is copied from the reasoning. Names are standardized with the metabolite-annotation-cleanup rules (PubChem's title when the exact, stereo-defined structure's record lists the proposed name; otherwise the cleaned label); standardization never changes a tier.

| Structure | Name | Tier | Notes |
|---|---|---|---|
| 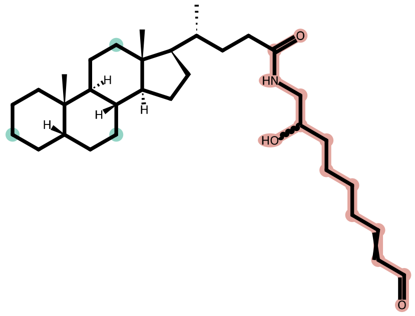<br>+ 1 OH, 1 OH (positions not resolved) | **C24 dihydroxy bile-acid C9H17NO2 amidate (partial structure)**<br>partial structure<br>C33H55NO5 · 545.40802 Da<br>drawn skeleton: C33H55NO3 | **Tentative** | Formula-level cargo and positional ambiguity are displayed, not a named compound assignment.<br>Wavy bonds: stereochemistry not resolved at 1 other<br>1 × OH on C12 (shaded); positions not resolved<br>1 × OH on C3/C7 (shaded); positions not resolved<br>Highlighted unit: C-24 amide linkage favoured<br>Highlighted unit: formula-valid drawing of C9H17NO2 only; internal connectivity is unresolved<br>Not resolved:<br>• whether the second hydroxyl is at C3 or C7<br>• cargo constitutional identity and functional-group layout<br>• hydroxyl stereochemistry outside the tree's 12α branch<br>• C5 ring fusion and other stereocentres not set by MS/MS<br>Assumed, not shown by this spectrum:<br>• standard C24 vertebrate bile-acid backbone stereochemistry<br>• C9H17NO2 cargo shown with a formula-valid placeholder topology solely to render the measured composition<br>Name: partial structure (substituent positions not resolved): no PubChem lookup; cleaned label shown |

#### 11. Network Analogs (graphml; hypotheses about relatedness, not evidence)

Network analogs from network_recovered.graphml: query node 1614 (m/z 546.4149), component of 1 node(s), 0 direct neighbour(s), 0 library-annotated. Shown: every library-annotated node and every direct neighbour.

---

*Generated by the bile-acid-msms-annotation-v5-6 skill (deterministic MassQL pre-pass, pure evaluator) plus literature-grounded reasoning in the referenced skill files. Source: P1_G7_Sample_X3148665.mzML, scan 2595. Run time: 27 min 8 s, start to finish (started 2026-10-27 17:04:00 UTC, report rendered 2026-10-27 17:31:08 UTC).*
