## Supplementary File 4 for "Capturing human expertise for AI-assisted molecular structure annotation"

### Bile-Acid MS/MS Annotation Report

SW034_SOWETO_3_B10.mzML — Scan 4280 | Generated with the "bile-acid-msms-annotation-v57" skill

Run time: 8 min 40 s, start to finish (started 2026-11-04 19:53:31 UTC, report rendered 2026-11-04 20:02:11 UTC)

#### 1. Precursor / Input Summary

| Field | Value | How obtained |
|---|---|---|
| Precursor m/z | 622.33178 | mzML scan header |
| Ion form | [M+H]+ (adopted; probable) — the Step 0 screen's [M+Na]+ choice is rejected | MS1 scan 4273: [M+Na]+ 644.3138 (–0.4 ppm) and [M+K]+ 660.2867 (–1.9 ppm) partners with the same isotope shape as 622.33; the co-eluting 600.3505 that supported [M+Na]+ has a different M+1 ratio (29.9 % vs 45.8 %), so it is another compound. Step 0 had no F in its element set. See C11 |
| Neutral formula (C/H/N/O/S + F, fitted by hand) | C32H48FN3O6S, M = 621.3248 ([M+H]+ calc 622.3321, –0.5 ppm). The script's CHNOS list (under [M+Na]+) is set aside; it had no F | manual formula search with F added (SKILL.md Step 3, drug cargos); fragments C6 – C9 |
| Retention time | 4.756 min (10.0 min total run) | mzML |
| Polarity / collision energy | positive ESI / HCD 41.6667 | mzML; filter: FTMS + p ESI d Full ms2 622.3318@hcd41.67 [65.5288-655.2884] |
| Instrument | Orbitrap Exploris 240 (Xcalibur 4.2-4.2.319.30-SP4/4.2.362.42; pwiz 3.0.24230) | mzML header |
| Isolation window | 622.332 –0.5/+0.5 Da; 1 MS1 ion(s) ≥1% inside it | mzML; MS1 scan 4273 |
| Peaks in MS2 | 66; base peak 130.0411 | mzML |

| Field | Value | How obtained |
|---|---|---|
| Other MS2 on this precursor (whole run) | scan 4289 @ 4.766 min | extract_context.py |
| Engine cross-check | not run (pure evaluator only; no borderline ratio) | --engine pure (default) |

Collision energy 41.6667 is below the NCE ~44–45 at which the dust-region rules were characterised. The dust region was still evaluated; expect fewer and weaker dust ions and some ratio shift. Lower energy alone does not lower a tier (stereochemistry_and_caveats.md §4).

#### 2. Annotated MS/MS Spectrum

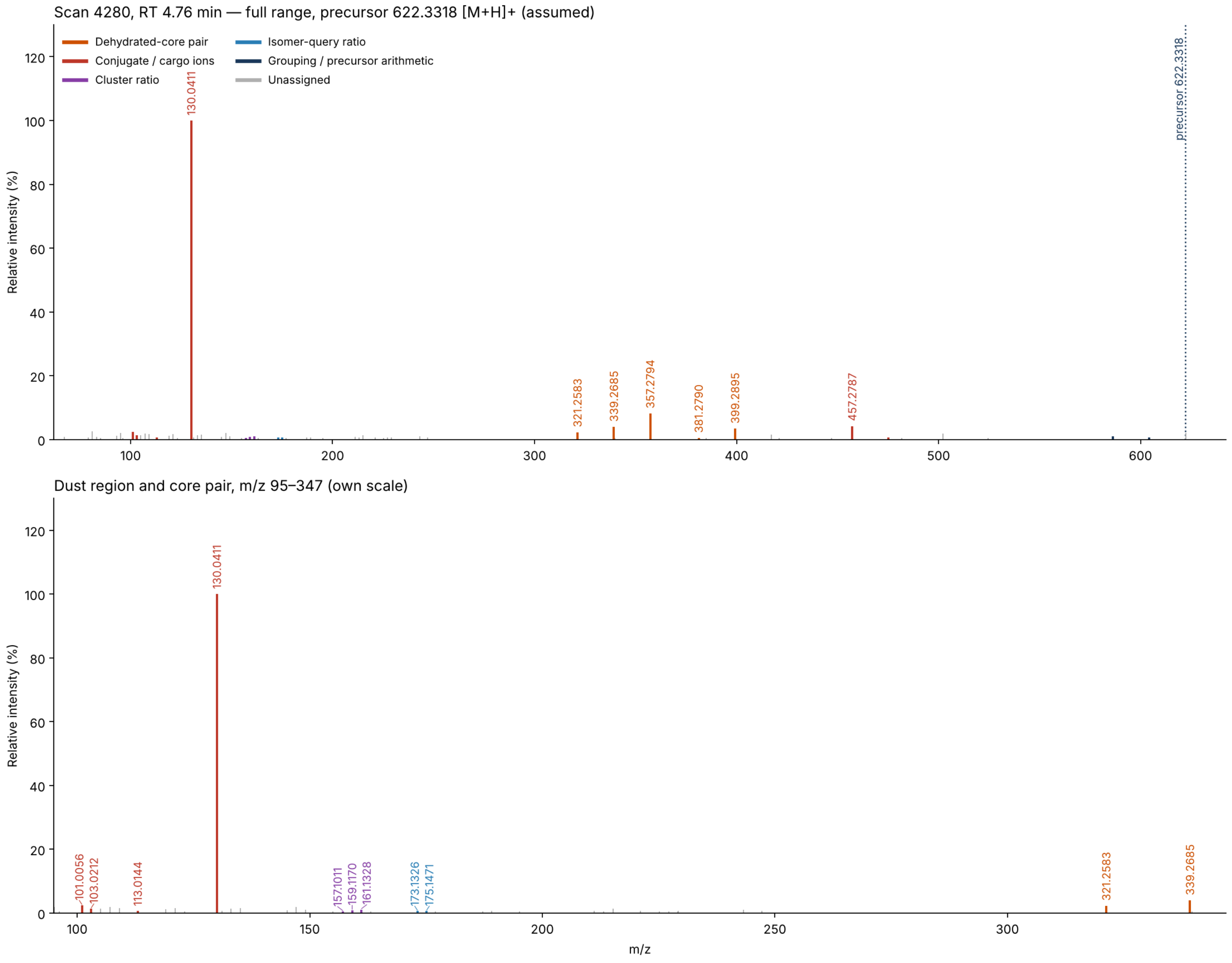


Scan 4280 (RT 4.76 min, precursor 622.3318 [M+H]+). Top: full range. Bottom: m/z 95–347 on its own scale, where the cargo ions and the ring-derived “dust” sit. Colors follow the shared palette; gray stems are unassigned peaks.

#### 3. Proposed Annotation

C24oic(OH)2–O-FTC (C-24 ester), hydroxyl positions unresolved — Dihydroxy C24 bile acid esterified at its side-chain carboxyl to the hydroxymethyl of emtricitabine (FTC) — i.e. **formula consistent with an emtricitabine conjugate of a dihydroxy-cholanoic acid (e.g. deoxycholoyl- or chenodeoxycholoyl-emtricitabine)** — hydroxyl positions, α/β, C5 and confirmation of the drug identity (standard or dosing records) remain open; overall tentative.

| Aspect | Call | Tier |
| --- | --- | --- |
| Bile acid (not bile alcohol, not a contaminant class) | Yes: dust passes, C24 dihydroxy core ions, bile-acid acyl ions still bound to the cargo — see C1 – C3 , C8 – C9 | Probable |
| Ion form | [M+H]+ (Step 0 [M+Na]+ rejected) — C11 , X16 | Probable |
| Core carbon count | C24 | Probable |
| Hydroxylation count | 2 (true dihydroxy; the precursor formula leaves no room for a ketone) | Probable |
| Cargo / conjugate | C8H10FN3O3S condensed (–H2O); formula and fragments consistent with emtricitabine — see A1 – A2 | Tentative |
| Bond to the bile acid | Ester of C-24 carboxyl to the emtricitabine hydroxymethyl — see A3 | Tentative |
| Hydroxyl positions | — | Unresolved |
| α/β stereochemistry, C5 ring fusion | — | Unresolved |

[Structures] skeleton family 2 (standard C24 cholanic core), R = O-CH2-(emtricitabine oxathiolane), two OH on rings A–D at unresolved positions.

#### 4. Evidence Table

##### Confirmatory (positive) evidence

| Ref | Obs. m/z | %base | Assignment | Calc. | Δppm | Interpretation | Source |
|---|---|---|---|---|---|---|---|
| C1 | 339.2685<br>321.2583 | 4.00<br>2.28 | C24H35O+<br>C24H33+ | 339.2682<br>321.2577 | +0.8<br>+1.9 | Core-class query **C24 - Dihydroxy** passed: dehydrated-core pair present (query targets 339.27, 321.26, ±0.01 Da, ≥1% each — floor adapted from the published 5%; *passes only under the adapted floor (relaxed gate): the 339/321 pair alone would cap the core call at tentative; the published-floor pass comes from* C2 ); the tree's **Dihydroxy_stage1** query passes on the same two ions — "the more hydroxyls, the smaller the number". 321.2583 is read as C24H33+ (+1.9 ppm); the script's C19H38SNa+ needs Na, which the precursor does not carry. | carbon_count_massql_table.md — [TrackSheet], [Mohanty2026-AC]; floor 5 % → 1 % [Adaptation-2026-10-18]; scripts/rules/massql_queries.yaml — [Mohanty2026-AC], [Transcript-UCSD9] |
| C2 | 357.2794<br>339.2685<br>339.2685<br>357.2794 | 8.15<br>4.00<br>4.00<br>8.15 | C24H37O2+<br>C24H35O+<br>C24H35O+<br>C24H37O2+ | 357.2788<br>339.2682<br>339.2682<br>357.2788 | +1.6<br>+0.8<br>+0.8<br>+1.6 | Core-class query **C24 alkamine - Dihydroxy** passed: dehydrated-core pair present (query targets 357.279, 339.268, 339.268, 357.279, ±0.01 Da) — "the more hydroxyls, the smaller the number". Here 357.2788 (C24H37O2+, [dihydroxy acid + H – 2H2O]+ side-chain-retained ion) is read as a dihydroxy C24 acyl core, *not* as alkamine evidence: the cargo carries no alkamine (Step 4 caution: free and esterified acids also give 357). | carbon_count_massql_table.md — [TrackSheet], [Mohanty2026-AC], [Transcript-UCSD9] |

| Ref | Obs. m/z | %base | Assignment | Calc. | Δppm | Interpretation | Source |
|---|---|---|---|---|---|---|---|
| C3 | 10 ring ions | ≥1 | odd-mass C/H/O | — | max \|2.0\| | Dust region m/z 100–250: 10 ring-derived ions (need 7) → passes: 105.0698 ($C_8H_9^+$), 107.0856 ($C_8H_{11}^+$), 109.1012 ($C_8H_{13}^+$), 119.0857 ($C_9H_{11}^+$), 121.1011 ($C_9H_{13}^+$), 133.1013 ($C_{10}H_{13}^+$), 135.1168 ($C_{10}H_{15}^+$), 147.1171 ($C_{11}H_{15}^+$), 161.1328 ($C_{12}H_{17}^+$), 215.1791 ($C_{16}H_{23}^+$). Set aside as N/S-bearing (cargo-like): 101.0056, 103.0212, 130.0411 — “the dust tells you where the things are attached. It doesn't tell you... if it was alpha or beta.” | dust_region_rules.md — [Specialist heuristic — Revision-2026-09-21], [Transcript-UCSDxx] |
| C4 | 175.1471<br>173.1326 | 0.59<br>0.57 | $C_{13}H_{19}^+$<br>$C_{13}H_{17}^+$ | 175.1481<br>173.1325 | -6.1<br>+0.4 | Di-3,12b-OH\|Di-7,12b-OH query PASSED (3,12b-OH2; 7,12b-OH2): obs 0.980× vs required 1× ±20% (0.8–1.2) | scripts/rules/massql_queries.yaml — [Mohanty2026-AC] |
| C5 | 622.3318 | 2.06 | $C_{32}H_{49}FN_3O_6S^+$ [M+H]+ | 622.3321 | -0.5 | Exact mass fits a dihydroxy C24 acid ($C_{24}H_{40}O_4$) + emtricitabine ($C_8H_{10}FN_3O_3S$) – $H_2O$. Precursor – dihydroxy C24 acid core leaves 229.0318 Da (pre-pass residual), which is emtricitabine – $H_2O$ (229.0321). The CHNOS candidates the script listed under [M+Na]+ explain neither the F-bearing base peak nor the S-bearing fragments. — “close is not close enough... it has to be exact.” | precursor arithmetic (manual, F added); pre-pass residual_candidates; [Specialist heuristic] |

| Ref | Obs. m/z | %base | Assignment | Calc. | Δppm | Interpretation | Source |
|---|---|---|---|---|---|---|---|
| C6 | 130.0411<br>113.0144 | 100<br>0.59 | C4H5FN3O+<br>C4H2FN2O+ | 130.0411<br>113.0146 | +0.0<br>-1.9 | Base peak = protonated 5-fluorocytosine (the nucleobase of emtricitabine), with its NH3 loss. No C/H/N/O/S formula fits 130.0411 within 5 ppm without Na (formula_table gave C2H9N3SNa+, chemically implausible). Released unmodified, so the base is not acylated. | manual formula fit (F added); SKILL.md Step 3 drug cargos |
| C7 | 101.0056<br>103.0212 | 2.38<br>1.30 | C4H5OS+<br>C4H7OS+ | 101.0056<br>103.0212 | +0.5<br>-0.1 | Oxathiolane (sugar-ring) ions of emtricitabine. Best CHNO fits are 34 ppm off, so sulfur is required. This places the sulfur in the cargo and is a second, independent cargo ion type. | formula_table.py; stereochemistry_and_caveats.md §2a |
| C8 | 475.2874<br>457.2787 | 0.66<br>4.20 | C28H43O4S+<br>C28H41O3S+ | 475.2877<br>457.2771 | -0.6<br>+3.5 | [M+H – 5-fluorocytosine – H2O]+ and a further H2O loss: the bile-acid acyl stays attached to the S-bearing oxathiolane after the base leaves. So the bile acid is bonded to the sugar ring, not to the base, which points to an ester at the hydroxymethyl. Both still carry O on the core, so one ring OH remains free. | manual fragment arithmetic |
| C9 | 399.2895<br>381.2790 | 3.41<br>0.51 | C26H39O3+<br>C26H37O2+ | 399.2894<br>381.2788 | +0.4<br>+0.6 | 475.2874 – C2H4OS (75.998 Da, the S-containing part of the ring): a C24 acyl ion that keeps the ester oxygen and two oxathiolane carbons (plus –H2O for 381). It supports a C-24 ester and is inconsistent with an amide to N4. | manual fragment arithmetic |

| Ref | Obs. m/z | %base | Assignment | Calc. | Δppm | Interpretation | Source |
|---|---|---|---|---|---|---|---|
| C10 | 604.3210<br>586.3145 | 0.61<br>0.93 | C32H47FN3O5S+<br>— | 604.3215<br>586.3110 | -0.8<br>+6.0 | Water-loss ladder from the precursor (ring hydroxyls free). 586.3145 sits 6 ppm from −2H2O and is noted but not used as evidence. This ladder is typical of a protonated, not a sodiated, precursor. | manual fragment arithmetic |
| C11 | 644.3138<br>660.2867 | MS1<br>MS1 | [M+Na]+<br>[M+K]+ | 644.3140<br>660.2879 | -0.4<br>-1.9 | MS1 scan 4273: sodium and potassium partners of M = 621.3248. 644.3138 (1.5e6) has M+1 = 42.3 %, matching the precursor's 45.8 %. The 600.3505 ion that made [M+Na]+ look plausible has M+1 = 29.9 %, so it is a different molecule. Ion form [M+H]+ adopted. | ms1_context (ion_form candidates), manual isotope read |
| C12 | 624.3326 | 9.8 (of A) | A+2 ($^{34}S$ + $^{13}C_2$ unresolved) | ≈624.334 (S1 blend) | — | The A+2 centroid lies between $^{34}S$ (624.3276) and $^{13}C_2$ (624.3385), close to the blend expected for one S (≈624.334) and 5 mDa from the S-free value. The script flagged $^{34}S$ "absent" only because no peak sat exactly at 624.3276. Corroboration only (§2a). A+1 (45.8 %) is above the ~37 % predicted for C32; see Section 6. | stereochemistry_and_caveats.md §2a; formula_table s34_check |
| C13 | 157.1011<br>159.1170<br>161.1328 | 0.43<br>0.83<br>1.04 | C12H13+<br>C12H15+<br>C12H17+ | 157.1012<br>159.1168<br>161.1325 | -0.8<br>+1.1<br>+2.0 | 157/159/161 cluster, 159/161 = 0.80. This lies between the 3α,7α shape (159 ≈ 161) and the 3α,12α shape (161 dominant), at ≤ 1 % of base and at HCD 41.7, so no position call is made (heuristic, §A). | hydroxylation_position_rules.md §A — [Notebook p.15]; fragment_pattern_fingerprints.md |

#### Exclusionary (negative) evidence — what this rules out

| Ref | Not observed / failed | Rules out | Source |
|---|---|---|---|
| X1 | Dihydroxy_stage2 failed: no peak within 0.01 Da of 339.2700 at >= 5% base peak; no peak within 0.01 Da of 321.2600 at >= 5% base peak; no peak within 0.01 Da of 248.0424 at >= 5% base peak | The Dihydroxy_stage2 bin (Amidated dihydroxy) | scripts/rules/massql_queries.yaml — [Mohanty2026-AC] |
| X2 | Di-1-ketone failed: no peak within 20 ppm of 161.1320 at >= 95% base peak | The Di-1-ketone bin (1 ketone) — below the node where the tree stopped, so this is the query's own result, not a tree step | scripts/rules/massql_queries.yaml — [Mohanty2026-AC] |
| X3 | Di-3,12a-OH\|Di-7,12a-OH failed: no peak within 20 ppm of 201.1630 | The Di-3,12a-OH\|Di-7,12a-OH bin (3,12a-OH2; 7,12a-OH2) — below the node where the tree stopped, so this is the query's own result, not a tree step | scripts/rules/massql_queries.yaml — [Mohanty2026-AC] |
| X4 | Di-3,12a-OH failed: no peak within 20 ppm of 243.1740; no peak within 20 ppm of 239.1790 | The Di-3,12a-OH bin (3,12a-OH2) — below the node where the tree stopped, so this is the query's own result, not a tree step | scripts/rules/massql_queries.yaml — [Mohanty2026-AC] |
| X5 | Di-7,12a-OH failed: no peak within 20 ppm of 243.1740; no peak within 20 ppm of 239.1790 | The Di-7,12a-OH bin (7,12a-OH2) — below the node where the tree stopped, so this is the query's own result, not a tree step | scripts/rules/massql_queries.yaml — [Mohanty2026-AC] |
| X6 | Di-3,7-OH\|Di-3,6-OH\|Di-3,12b-OH\|Di-7,12b-OH failed: no peak within 20 ppm of 201.1630 | The Di-3,7-OH\|Di-3,6-OH\|Di-3,12b-OH\|Di-7,12b-OH bin (3,7-OH2; 3,6-OH2; 3,12b-OH2; 7,12b-OH2) — below the node where the tree stopped, so this is the query's own result, not a tree step | scripts/rules/massql_queries.yaml — [Mohanty2026-AC] |
| X7 | Di-3,7-OH\|Di-3,12b-OH\|Di-7,12b-OH failed: no peak within 20 ppm of 201.1630 | The Di-3,7-OH\|Di-3,12b-OH\|Di-7,12b-OH bin (3,7-OH2; 3,12b-OH2; 7,12b-OH2) — below the node where the tree stopped, so this is the query's own result, not a tree step | scripts/rules/massql_queries.yaml — [Mohanty2026-AC] |
| X8 | Di-3,7-OH failed: 173.132: obs 0.980× vs required 0.5× ±30% (0.35–0.65) | The Di-3,7-OH bin (3,7-OH2) — below the node where the tree stopped, so this is the query's own result, not a tree step | scripts/rules/massql_queries.yaml — [Mohanty2026-AC] |

| Ref | Not observed / failed | Rules out | Source |
|---|---|---|---|
| X9 | Di-3,6-OH failed: no peak within 20 ppm of 201.1630 | The Di-3,6-OH bin (3,6-OH2) — below the node where the tree stopped, so this is the query's own result, not a tree step | scripts/rules/massql_queries.yaml — [Mohanty2026-AC] |
| X10 | Core-class rows absent: C24 - Nonhydroxy (343.3/325.29); C24 - Monohydroxy (341.28/323.27); C24 - Trihydroxy (337.25/319.24); C24 - Tetrahydroxy (335.24/317.23); C24 - Pentahydroxy (333.22/315.21); C23 - Monohydroxy (327.27/309.26/309.26/285.25); C23 - Dihydroxy (325.25/307.24/307.24/283.24); C23 - Trihydroxy (323.24/305.23/305.23/281.23); C23 - Tetrahydroxy (321.22/303.21/303.21/279.22); C23 - Pentahydroxy (319.21/301.19/301.19/277.19); C27 - Monohydroxy (383.33/159.12); C27 - Dihydroxy (381.32/159.12); C27 - Trihydroxy (379.29/159.12); C27 - Tetrahydroxy (377.28/159.12); C27 - Pentahydroxy (375.27/159.12); Aceto dihydroxy (347.294/329.284); C24 alkamine - Monohydroxy (359.295/341.284/341.284/359.295); C24 alkamine - Trihydroxy (355.263/337.253/337.253/355.263); C24 alkamine - Tetrahydroxy (353.248/335.237/335.237/353.248); C24 alkamine - Pentahydroxy (351.232/333.221/333.221/351.232); C28 alcohol - 1OH (125.13/143.14); C28 alcohol - 2OH (123.12/141.13); C28 alcohol - 3OH (121.1/139.11) | The hydroxylation counts and core classes those rows define | carbon_count_massql_table.md — [TrackSheet], [Mohanty2026-AC] |
| X11 | no 86.0964/104.1070/124.9998/184.0733 (PC / lysoPC head group); no 149.0233/167.0339 (phthalate); no 85.0284 (acylcarnitine); no 44.0262 Da repeat series (PEG) (±0.005 Da, ≥1% base) | Those non-bile-acid contaminant classes | nomenclature_glossary.md |

| Ref | Not observed / failed | Rules out | Source |
|---|---|---|---|
| X12 | [M+NH4]+ 639.3583 absent from MS1 scan 4273 | Consistent with the ester call: the bile-acid carboxyl is blocked, and emtricitabine has no acid. Candidates with a free acidic group (free –COOH, –SO3H …) are disfavoured; blocked candidates (esters, amides, lactones) favoured. Tentative; names no substituent. Caveat: mobile-phase ammonium not stated: assumed present (most acetonitrile-containing buffers carry some) -- say so in the report; if the mobile phase in fact lacks ammonium, this absence means nothing. — “What we don't see is informative as well.” | dust_region_rules.md §5 / §5a — [Specialist heuristic — Revision-2026-10-12] |
| X13 | No N4-acyl-5-fluorocytosine ion ([BA-acyl–5-FC + H]+, C28H43FN3O4+, 504.3232); the base is released as free 5-FC (130.0411, base peak), and the acyl-retaining ions 475/457/399 contain no N or F | Amide linkage of the bile-acid carboxyl to the cytosine N4 (capecitabine-type N4-acyl) | manual fragment arithmetic |
| X14 | [emtricitabine + H]+ 248.0500 absent from the MS2 (and not seen in MS1 scan 4273 above 2e4) | Nothing on its own. An expected drug ion is missing, which counts mildly against the drug name; free emtricitabine is known to fall apart to m/z 130 at this energy | SKILL.md Step 3 rule (c) |
| X15 | No cytosine ion 112.0505 (C4H6N3O+) and no S-free sugar ions; every ion that keeps the cargo carries F (130/113) or S (101/103/475/457) | Lamivudine (non-fluorinated analogue) and 5′-deoxy-5-fluorocytidine or other S-free 5-fluorocytosine nucleosides (capecitabine metabolites), which do not fit the S-containing precursor formula | manual formula and fragment arithmetic |
| X16 | For [M+Na]+ (M = 599.3426), the "[M+H]+" partner 600.3505 should share the precursor's isotope shape; it does not (M+1 29.9 % vs 45.8 %). No Na-retaining fragments are needed to explain the MS2 | The Step 0 [M+Na]+ reading and its tyrosine/phenylalanine–C25 cargo fits | ms1_context; manual isotope read |
| X17 | The precursor is 18.0106 Da lighter than a non-covalent [bile acid + emtricitabine + H]+ cluster (640.3427), and the S-bearing ions 475/457 still hold the bile-acid core | An ion-source cluster of a free bile acid and free emtricitabine; the bond is covalent (condensation) | SKILL.md Step 3 (cluster vs conjugate) |

#### Cargo evidence — what is attached to the bile acid

Every cargo candidate still in play, grouped by the question it answers. Each row cites the C/X rows above that support or argue against it; candidates already ruled out appear only in the exclusionary table. The Section 3 calls and Section 6 open questions are drawn from this table.

| Ref | Candidate | Evidence for | Evidence against | Status | Tier |
|---|---|---|---|---|---|
| **1. Cargo composition** → C8H10FN3O3S, condensed with loss of H2O (tentative) | | | | | |
| A1 | **Emtricitabine (FTC), 4-amino-5-fluoro-1-[2-(hydroxymethyl)-1,3-oxathiolan-5-yl]pyrimidin-2-one** C8H10FN3O3S; precursor – C24H40O4 + H2O | Exact precursor mass; 5-fluorocytosine base ion and its NH3 loss; S-bearing oxathiolane ions; acyl–oxathiolane ions after base loss; human cohort site where FTC is a common antiretroviral C5 C6 C7 C8 C12 | No [FTC+H]+ 248.0500; no dosing record or standard; A+1 higher than predicted X14 | Favoured | Tentative |
| A2 | **Another C8H10FN3O3S 5-fluorocytosine oxathiolane (stereoisomer or regio-isomer of FTC)** same formula | Same ions expected; MS/MS cannot tell the enantiomers or anomers apart C6 C7 | No known exposure source other than the drug | Possible | Unresolved |
| **2. Bond between the bile acid and the cargo** → ester at C-24 to the oxathiolane hydroxymethyl (tentative) | | | | | |
| A3 | **C-24 carboxyl ester to the FTC CH2OH (5′-O-acyl)** blocked carboxyl, 0 free acidic groups | Base released unmodified; acyl stays on the S-ring after base loss; acyl ion with ester O + 2 C of the ring; no [M+NH4]+ (no free COOH); covalent, not a cluster C6 C8 C9 X12 X17 | No standard; esters can form during sample handling (not tested) | Favoured | Tentative |

#### Step-7 validation pass

Every peak was re-checked against an elemental formula (C, H, N ≤3, O, S ≤2, Na ≤1; even-electron ion, electron mass applied): 62 of 66 fit within ±10 ppm (largest |error| 7.3 ppm; ions used as evidence above: ≤6.1 ppm). No plausible formula: 96.1243, 339.6741, 481.8416, 524.5070 — not used as evidence. 15 ion(s) have a second formula within 5 ppm (see formula_table.tsv) and are named by both where cited. Sulfur near-miss check (§2a): not triggered on the precursor or the main S-bearing ions. Shared CHNO offset: none. $^{34}$S A+2 on the precursor: absent — no peak at the 34S position; weighs against S if the A+2 region was resolved.

Odd/even parity check (conjugate_fragments.md): 6 even-mass fragment(s) (130.0411, 130.0579, 502.2894, 586.3144, 604.3210, 622.3335) — each carries an odd number of N in its best formula. 6 odd-mass ion(s) carry an even number of N (e.g. N2). The nitrogen rule holds for every assigned ion.

#### 5. Why the Automated Tree Stalled at "Dihydroxy_stage1"

The pre-pass reached **Dihydroxy_stage1** (not a leaf): Dihydroxy_stage2 did not match. The unmet conditions:

- **Dihydroxy_stage2**: MS2PROD=339.27 resolves to **339.2700** — no peak within 0.01 Da of 339.2700 at >= 5% base peak. Target – H2O (321.2594) **is present**: 321.2583 at 2.28% of base (C19H38SNa+, -1.1 ppm; or C24H33+, +1.9 ppm) — a dehydrated cargo ion would sit exactly here.
- **Dihydroxy_stage2**: MS2PROD=321.26 resolves to **321.2600** — no peak within 0.01 Da of 321.2600 at >= 5% base peak. Target – H2O (303.2494) is also absent.
- **Dihydroxy_stage2**: MS2PROD=X-374.2894 resolves to **248.0424** — no peak within 0.01 Da of 248.0424 at >= 5% base peak. (X = precursor 622.3318 as acquired, adduct assumed.) Target – H2O (230.0318) is also absent.

The stall is mechanical. Dihydroxy_stage2 is built for amidates: it needs 339/321 at ≥ 5 % and an amidated-cargo ion at X – 374.2894. In this ester, the collision energy went into the 5-fluorocytosine base peak, so the core ions sit at 2–8 %, and the cargo ion the query targets does not exist here. The position call stays unresolved. Hydrolysing the ester and analysing the freed bile acid, or MS3 on 357.28 or 475.29, would let the tree run.

#### 6. Unresolved Aspects

1. **Hydroxyl positions: cannot be determined.** The tree stopped at Dihydroxy_stage1 (Section 5). The 201/211 rule is validated for amidates, not esters (201 absent, X3 ). The 157/159/161 shape falls between the 3α,7α and 3α,12α patterns ( C13 ). DCA, CDCA, UDCA, HDCA and others remain possible. Hydrolysing the ester and running the freed acid against standards would settle it.
2. **α/β at the hydroxyls and C5 (5α/5β): not resolvable from MS/MS alone.** (stereochemistry_and_caveats.md §1). No RT-order corroboration: this FILENAME is not in the internal Orbitrap library (run_not_in_library).
3. **Drug identity is a formula-and-fragment match, not a confirmed identification.** Two independent ion types support it ( C6 , C7 ), but no dosing record and no synthetic standard are available, and [FTC+H]+ is absent ( X14 ). So it stays tentative: "formula consistent with an emtricitabine conjugate".
4. **Biological vs artefactual origin.** Whether the ester forms in vivo (host or microbial acyl transfer) or during extraction or storage was not tested. A blank matrix spiked with FTC and a bile acid would show whether it forms during handling.
5. **Isotope A+1 is higher than predicted.** 45.8 % observed vs ~37 % for C32H49FN3O6S+ (the [M+Na]+ partner gives 42.3 %). Orbitrap isotope ratios at low ion counts can be off this much, but an unrecognised co-isolated ion cannot be ruled out. 502.2894, 417.2374 and 586.3145 are not assigned.
6. **Pre-pass flags, as handled here.** (i) Two core classes matched (C24 dihydroxy and C24 alkamine dihydroxy). The alkamine hit is mechanical: 357/339 also come from esterified acids, and no alkamine is present. (ii) The MS1 ion form was ambiguous. It is settled here as [M+H]+ by isotope shape ( C11 ), at probable, not confirmed. (iii) Collision energy 41.7 is below NCE 44–45: dust ions are weak (all ≤ 2.5 %), and the energy went into the 130 base peak. (iv) Mobile-phase ammonium was not stated and was assumed present ( X12 ).
7. **No network context.** The GNPS2 graphml could not be downloaded in this session (direct access to gnps2.org was blocked). Section 11 is absent and no network neighbours were checked for related FTC conjugates.

#### 7. Species / Taxonomic Context

Prior, not evidence. Nothing here changes a tier.

| Part | Finding | Source |
|---|---|---|
| **Host prior** | Probably human. The file name (SOWETO) and other samples in the GNPS2 job (AGINCOURT, NANORO) are the names of African population-cohort sites. This is inferred from the names only; it is not recorded in the mzML. | file name; GNPS2 feature table (top samples) |
| **Expected producer** | Not documented. Bile acid–drug esters are not an established host or microbial product. Microbial bile-salt hydrolase acyl transfer is known for amidation; esterification to a drug hydroxyl has not been shown. | source_and_diet_context.md §3 (no matching record) |
| **Plausible route** | Drug-derived plausible (diet not provided and not relevant). Emtricitabine is part of first-line HIV treatment and PrEP regimens (with tenofovir). Dosing for SW034 is unknown. | source_and_diet_context.md §3 |

Follow-ups that would test the route:

- Check antiretroviral records for participant SW034 (FTC vs lamivudine-based regimen).
- Look for free emtricitabine (248.0500) and tenofovir in the same run, and for mono-/trihydroxy BA–FTC esters ([M+H]+ 606.337 / 638.332) across the cohort.
- Synthesize deoxycholoyl- and chenodeoxycholoyl-FTC esters as standards for RT and MS2.
- Spike FTC and a bile acid into blank matrix to test for ex vivo ester formation.

#### 8. Chromatographic Context (Supporting, Not Evidence)

Extracted-ion chromatograms (±5 ppm, whole 10.0-min run) were computed by the script under the rejected [M+Na]+ basis (M = 599.34256). Only the precursor trace applies to the adopted [M+H]+ reading. The others are kept, relabelled, for transparency; the MS1 partners of the adopted form are added from scan 4273:

| Trace | Chromatographic peaks (RT min, apex intensity) |
|---|---|
| 622.3318 [M+H]+ (precursor) | 4.54 (6.9e4), 4.75 (2.4e6) ← this scan |
| 644.3138 [M+Na]+ (MS1 scan 4273 only) | 4.75 (1.5e6), same isotope shape as 622.33 |
| 660.2867 [M+K]+ (MS1 scan 4273 only) | 4.75 (6.5e4) |
| 600.3498: co-eluting, different compound (M+1 29.9 %) | 3.15 (4.0e4), 4.76 (1.3e6) |
| 617.3764 / 638.3057 / 582.3393 (only meaningful under [M+Na]+) | no peak co-eluting at 4.76 except 582.34 (9.1e4) |
| GNPS2 feature 36272 (orientation) | 622.3318, RT 4.746, EB adduct call [M+H]1+, top sample = this file (2.4e6). Other 622.33 features: 74918 (622.3314, RT 4.261, BF225_NANORO), MS2 not retrieved; a possible isomer to check |

Supporting orientation only: retention time alone identifies nothing (stereochemistry_and_caveats.md). 1 other MS2 scan(s) were acquired on this precursor in the run.

#### 9. Diagnostic Decision Tree

Each node's accent color is the color of the corresponding ion family highlighted in the Section 2 spectrum. The numbered chips (C = confirmatory, X = exclusionary, A = cargo evidence) are the Ref column of the Section 4 evidence tables, so every node traces back to its ions, mass errors and sources.

Dehydrated-core pair · Conjugate / cargo ions · Cluster ratio · Isomer-query ratio

Grouping / precursor arithmetic · Ruled out (no diagnostic ion family)

✓ supports the annotation · ⊘ alternative ruled out (dashed box) · – context only

**MS2 scan 4280**
precursor 622.3318 [M+H]+ · 66 peaks · base peak 130.0411

**Core identity & conjugate**
carbon count, OH count, cargo, bond

- ✓ **C24 dihydroxy core ions** — 357.2794 / 339.2685 / 321.2583 — C1 C2
- ✓ **Dust region passes** — 10 odd-mass ring ions (≤ 2.5 % each) — C3
- ✓ **Exact mass C32H48FN3O6S** — [M+H]+ −0.5 ppm; Na/K partners in MS1 — C5 C11
- ✓ **5-Fluorocytosine base peak** — 130.0411 (100 %), 113.0144 — C6
- ✓ **Oxathiolane S ions** — 101.0056 / 103.0212; A+2 centroid fits one S — C7 C12
- ✓ **Acyl kept on S-ring** — 475/457/399 → C-24 ester to CH2OH — C8 C9
- – **No [FTC+H]+** — 248.0500 absent — X14

**Hydroxylation position**
automated tree stalled at Dihydroxy_stage1 (Section 5)

- ✗ **Stage 2 not reached** — amidate-specific query; core ions < 5 % — X1
- – **157/159/161 ≈ 0.4/0.8/1.0** — between the 3,7 and 3,12 shapes; no call — C13

**Alternatives ruled out**
what this is not

- ⊘ **[M+Na]+ reading** — isotope shape mismatch with 600.35 — X16
- ⊘ **N4-amide linkage** — base released unmodified — X13
- ⊘ **Lamivudine / S-free fluorocytidines** — F and S both required — X15
- ⊘ **Non-covalent cluster** — precursor = BA + FTC – H2O — X17
- ⊘ **Contaminant classes** — no PC, phthalate, PEG, acylcarnitine ions — X11

↓

**Dihydroxy C24 bile acid C-24 ester of emtricitabine (formula-consistent) (Tentative)**

core probable; drug cargo and ester bond tentative; positions and stereochemistry unresolved

#### 10. Proposed Final Structures

One row per structure the reasoning in Sections 3–9 still supports, drawn with RDKit with the ABCD rings in the standard bile-acid orientation. A partial structure draws only what the evidence claims: wavy bonds mark stereocentres that are not resolved, substituents that are counted but not placed are listed under the drawing and their possible rings shaded, and a highlighted unit is weaker than the rest (the Notes say why). The Tier column is copied from the reasoning. Names are standardized with the metabolite-annotation-cleanup rules (PubChem's title when the exact, stereo-defined structure's record lists the proposed name; otherwise the cleaned label); standardization never changes a tier.

| Structure | Name | Tier | Notes |
|---|---|---|---|
| 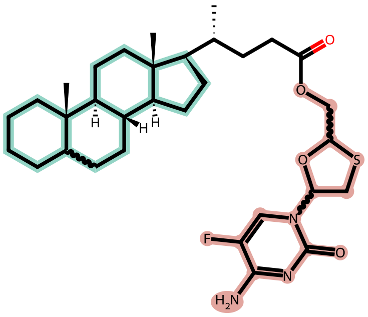<br>+ 2 OH (positions not resolved) | **dihydroxy-5ξ-cholan-24-oyl emtricitabine ester (partial structure)**<br>partial structure<br>C32H48FN3O6S · 621.32479 Da<br>drawn skeleton: C32H48FN3O4S | Tentative | Single partial structure: two OH unplaced on rings A–D.<br>Wavy bonds: stereochemistry not resolved at C5, 2 other<br>2 × OH on rings A–D (shaded); positions not resolved<br>Highlighted unit: emtricitabine unit: formula and fragments only, no standard<br>Not resolved:<br>• positions of the two OH (hydrolysis + standards would settle)<br>• α/β of each OH; C5 (5α/5β)<br>• oxathiolane stereocentres (the drug is (2R,5S); not drawn)<br>• emtricitabine vs another C8H10FN3O3S isomer<br>Assumed, not shown by this spectrum:<br>• backbone stereocentres 8β,9α,10β,13β,14α,17β and 20R of a normal C24 bile acid<br>Name: partial structure (substituent positions not resolved): no PubChem lookup; cleaned label shown |

*Generated by the "bile-acid-msms-annotation-v57" skill (deterministic MassQL pre-pass, pure evaluator) plus literature-grounded reasoning in the referenced skill files. Source: SW034_SOWETO_3_B10.mzML, scan 4280. Run time: 8 min 40 s, start to finish (started 2026-11-04 19:53:31 UTC, report rendered 2026-11-04 20:02:11 UTC).*
