## Supplementary File 5 for "Capturing human expertise for AI-assisted molecular structure annotation"

### Bile-Acid MS/MS Annotation Report

AG078_AGINCOURT_6_B6.mzML — Scan 3956 | Generated with the "bile-acid-msms-annotation-v57" skill

Run time: 6 min 37 s, start to finish (started 2026-11-04 19:39:08 UTC, report rendered 2026-11-04 19:45:45 UTC)

#### 1. Precursor / Input Summary

| Field | Value | How obtained |
|---|---|---|
| Precursor m/z | 604.34142 | mzML scan header |
| Ion form | [M+H]+ (inferred from fragment formulas; MS1 alone does not settle it) | MS1 scan 3954 shows z = 1 (spacing 1.0035) but cannot separate [M+H]+ / [M+K]+ / [M+H-H2O]+. The Step 0 screen (C/H/O + listed cargos only) preferred [M+K]+; only the [M+H]+ neutral C32H49N3O6S contains the N3 and S carried by the fragments C4 – C8 , and the K-form neutrals (one N) cannot ( X5 ). GNPS2 everything_bagel calls feature 15549 [M+H]1+. Basis set with extract_context.py --ion-form (new option). |
| Neutral formula candidates (±5 ppm, C/H/N≤3/O/S≤2) | C32H49N3O6S (-0.11 ppm); also within 5 ppm: C40H45NO4 (-1.19 ppm); C36H49N3OS2 (+4.04 ppm); C28H49N3O11 (-4.26 ppm) | formula_table.py (mass alone does not choose among these) |
| Retention time | 4.427 min (10.0 min total run) | mzML |
| Polarity / collision energy | positive ESI / HCD 41.6667 | mzML; filter: FTMS + p ESI d Full ms2 604.3414@hcd41.67 [63.6938-636.9382] |
| Instrument | Orbitrap Exploris 240 (Xcalibur 4.2-4.2.319.30-SP4/4.2.362.42; pwiz 3.0.24230) | mzML header |

| Field | Value | How obtained |
|---|---|---|
| Isolation window | 604.341 −0.5/+0.5 Da; 1 MS1 ion(s) ≥1% inside it | mzML; MS1 scan 3954 |
| Peaks in MS2 | 100; base peak 112.0505 | mzML |
| Other MS2 on this precursor (whole run) | scan 3967 @ 4.439 min | extract_context.py |
| Engine cross-check | not run (pure evaluator only; no borderline ratio) | --engine pure (default) |

Collision energy 41.6667 is below the NCE ~44–45 at which the dust-region rules were characterised. The dust region was still evaluated; expect fewer and weaker dust ions and some ratio shift. Lower energy alone does not lower a tier (stereochemistry_and_caveats.md §4).

#### 2. Annotated MS/MS Spectrum

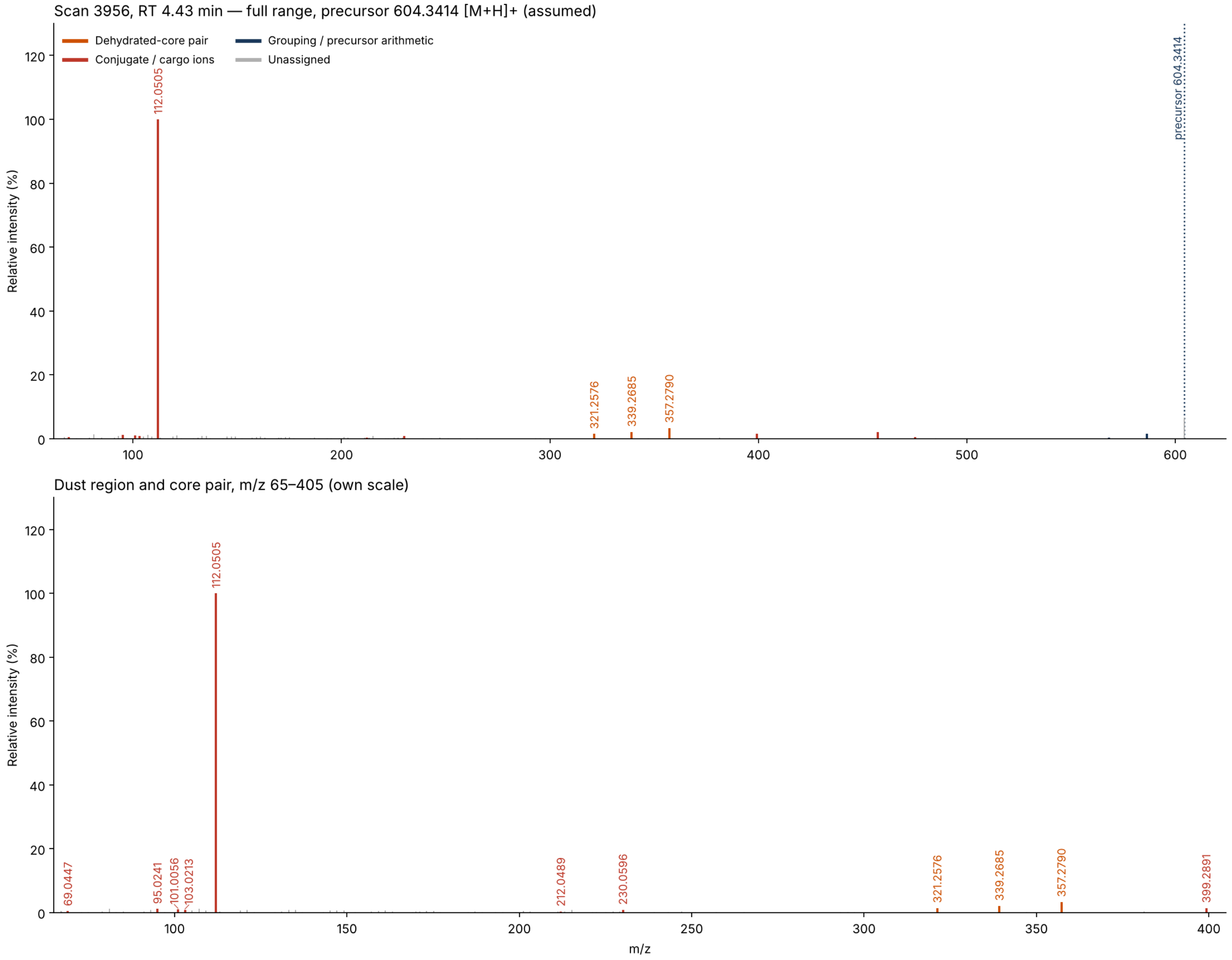


Scan 3956 (RT 4.43 min, precursor 604.3414 [M+H]+). Top: full range. Bottom: m/z 65–405 on its own scale, where the cargo ions and the ring-derived “dust” sit. Colors follow the shared palette; gray stems are unassigned peaks.

#### 3. Proposed Annotation

C24oic(OH)2-O-(C8H10N3O2S), a 24-O-ester — Dihydroxy C24 bile acid esterified at C-24 to a C8H11N3O3S nucleoside analogue (cytosine + oxathiolane) — i.e. **formula consistent with a lamivudine (3TC) 5′-O-(dihydroxycholan-24-oyl) ester** hydroxyl positions, α/β and the 3TC-vs-isomer identity unresolved; tentative pending dosing records and a synthetic standard.

| Aspect | Call | Tier |
|---|---|---|
| Bile acid (not bile alcohol, not a contaminant class) | Yes — C24 dihydroxy pair at 1.4–2.1 % (relaxed gate; published 5 % floor not met) | Tentative |
| Ion form | [M+H]+, from fragment formulas (see C3 , X5 ) | Probable |
| Core carbon count | C24 (357/339/321 ladder, precursor arithmetic) | Probable |
| Hydroxylation count | 2 OH (no ketone: the precursor formula leaves none) | Probable |
| Cargo composition | C8H11N3O3S — see A1 – A2 | Probable |
| Cargo identity | lamivudine-type cytosine oxathiolane nucleoside — see A3 – A5 | Tentative |
| Bond to the bile acid | C-24 ester to the cargo's CH2OH — see A6 – A7 | Tentative |
| Hydroxyl positions | not determined (dust region fails; no tree path) | Unresolved |
| α/β stereochemistry, 5α/5β | not resolvable from MS/MS alone | Unresolved |
| Overall | tentative | Tentative |

[Structures] skeleton family 2 (standard C24 cholanic core) with R = O-ester to the cargo, two OH at unassigned ring positions.

#### 4. Evidence Table

##### Confirmatory (positive) evidence

| Ref | Obs. m/z | %base | Assignment | Calc. | Δppm | Interpretation | Source |
|---|---|---|---|---|---|---|---|
| C1 | 339.2685<br>321.2576 | 2.12<br>1.43 | C24H35O+<br>C24H33+ | 339.2682<br>321.2577 | +0.8<br>-0.1 | Core-class query **C24 - Dihydroxy** passed: dehydrated-core pair present (query targets 339.27, 321.26, ±0.01 Da, ≥1% each — floor adapted from the published 5%; *passes only under the adapted floor (relaxed gate): core call and overall tier ≤ tentative; later steps graded on their own evidence*) — “the more hydroxyls, the smaller the number” | carbon_count_massql_table.md — [TrackSheet], [Mohanty2026-AC]; floor 5 % → 1 % [Adaptation-2026-10-18], [Transcript-UCSD9] |

| Ref | Obs. m/z | %base | Assignment | Calc. | Δppm | Interpretation | Source |
|---|---|---|---|---|---|---|---|
| C2 | 1 ring ions | ≥1 | odd-mass C/H/O | — | max \|0.1\| | Dust region m/z 100–250: 1 ring-derived ions (need 7) → does not pass: 107.0855 (C8H11+). Set aside as N/S-bearing (cargo-like): 112.0505 — “the dust tells you where the things are attached. It doesn't tell you... if it was alpha or beta.” | dust_region_rules.md — [Specialist heuristic — Revision-2026-09-21], [Transcript-UCSDxx] |
| C3 | 603.33414 (neutral) | — | C32H49N3O6S | 603.33421 | -0.1 | Exact-mass fit of the neutral from [M+H]+. Of the four formulas within 5 ppm, only C32H49N3O6S contains every fragment formula: the N3 cytosine ions C4 , the N3S ion 230.0596 C5 and the S-bearing, N-free ions 475/457 C8 . C40H45NO4 (one N, no S) and C36H49N3OS2 (one O, cannot hold C24H37O2+ plus C8H12N3O3S+) and C28H49N3O11 (no S) are not consistent with them. C32H49N3O6S = C24H40O4 (dihydroxy C24 acid) + C8H11N3O3S – H2O. MS1 A+1 = 37.0 % of the monoisotopic peak (C32 predicts ≈ 37 %; a C35–C36 [M+K]+ neutral ≈ 39–41 %). — “close is not close enough... it has to be exact.” | precursor arithmetic (formula_table.py), [Specialist heuristic] |
| C4 | 112.0505<br>95.0241<br>69.0447 | 100<br>1.12<br>0.46 | C4H6N3O+<br>C4H3N2O+<br>C3H5N2+ | 112.0505<br>95.0240<br>69.0447 | -0.4<br>+1.3<br>-0.2 | Base peak = protonated cytosine [B+H]+, with its two standard daughters, –NH3 (95.0241) and –HNCO (69.0447). The nucleobase is released *unmodified*, so the bile acid is not on the base (see X4 ). | formula_table.py; exact-mass formula fit [Specialist heuristic] |

| Ref | Obs. m/z | %base | Assignment | Calc. | Δppm | Interpretation | Source |
|---|---|---|---|---|---|---|---|
| C5 | 230.0596<br>212.0489 | 0.86<br>0.34 | C8H12N3O3S+<br>C8H10N3O2S+ | 230.0594<br>212.0488 | +0.7<br>+0.2 | Intact cargo [X+H]+ C8H12N3O3S+ and its −H2O ion. 230.0596 is also exactly precursor − 374.2821 (the dehydrated C24 dihydroxy core of the Step 0 ladder; calc 230.0593), i.e. the cargo-retaining residual of that rung — below the screen's 2 % floor, so it is reported here, tentative. C8H11N3O3S is the formula of lamivudine (3TC) and its isomers (apricitabine, the 3TC enantiomer). | formula_table.py ; Step 0 B ladder arithmetic [Benchmark201, hypothesis grade] |
| C6 | 101.0056<br>103.0213 | 0.97<br>0.81 | C4H5OS+<br>C4H7OS+ | 101.0056<br>103.0212 | +0.5<br>+0.9 | N-free, S-bearing C4 ions: the oxathiolane (sugar-like) ring of the cargo after loss of the base. Second CHO formulas (C7HO+, C7H3O+) are implausible (RDBE/H count). Supports a C4 O,S-heterocycle carrying the base. | formula_table.py [Specialist heuristic] |
| C7 | 357.2790 | 3.33 | C24H37O2+ | 357.2788 | +0.5 | Side-chain-retained dihydroxy C24 ion (acylium − H2O) — the strongest bile-acid ion; with 339.2685 and 321.2576 C1 it forms the 357/339/321 water ladder of a dihydroxy C24 acyl unit. Not an alkamine marker here: the precursor formula has no room for a C24 alkamine core (Step 4 note). | carbon_count_massql_table.md, conjugate_fragments.md §Free bile acids |
| C8 | 475.2876<br>457.2773 | 0.51<br>2.05 | C28H43O4S+<br>C28H41O3S+ | 475.2877<br>457.2771 | -0.1<br>+0.6 | [M+H − cytosine − H2O]+ and [M+H − cytosine − 2H2O]+ (604.3414 − 111.0433 − n·18.0106). The bile-acid acyl stays bound to the S-bearing oxathiolane after the base leaves ⇒ the acyl is on the sugar ring (its 5′-CH2OH), not on the cytosine NH2. | formula_table.py ; neutral-loss arithmetic [Specialist heuristic] |

| Ref | Obs. m/z | %base | Assignment | Calc. | Δppm | Interpretation | Source |
|---|---|---|---|---|---|---|---|
| C9 | 399.2891 | 1.44 | C26H39O3+ | 399.2894 | -0.8 | Acyl ion + C2H2O (357.2790 + 42.0106): the acyl still carrying an extra oxygen and two carbons, i.e. the ester O–CH2–C fragment of the alcohol. Fits an O-ester; an amide would carry N instead. Assignment tentative (single ion). | formula_table.py ; [NashDunn2024] C2H2O 42.0106 (hypothesis grade) |
| C10 | 586.3322<br>568.3220 | 1.52<br>0.36 | C32H48N3O5S+<br>C32H46N3O4S+ | 586.3309<br>568.3204 | +2.2<br>+2.8 | Two sequential water losses from the precursor: consistent with two hydroxyls on the core (formula_table.py's first choices here are other C/H/N/O formulas; the sub-formulas of the precursor are listed instead, within 3 ppm). | precursor arithmetic |

#### Exclusionary (negative) evidence — what this rules out

| Ref | Not observed / failed | Rules out | Source |
|---|---|---|---|
| X1 | Core-class rows absent: C24 - Nonhydroxy (343.3/325.29); C24 - Monohydroxy (341.28/323.27); C24 - Trihydroxy (337.25/319.24); C24 - Tetrahydroxy (335.24/317.23); C24 - Pentahydroxy (333.22/315.21); C23 - Monohydroxy (327.27/309.26/309.26/285.25); C23 - Dihydroxy (325.25/307.24/307.24/283.24); C23 - Trihydroxy (323.24/305.23/305.23/281.23); C23 - Tetrahydroxy (321.22/303.21/303.21/279.22); C23 - Pentahydroxy (319.21/301.19/301.19/277.19); C27 - Monohydroxy (383.33/159.12); C27 - Dihydroxy (381.32/159.12); C27 - Trihydroxy (379.29/159.12); C27 - Tetrahydroxy (377.28/159.12); C27 - Pentahydroxy (375.27/159.12); Aceto dihydroxy (347.294/329.284); C24 alkamine - Monohydroxy (359.295/341.284/341.284/359.295); C24 alkamine - Dihydroxy (357.279/339.268/339.268/357.279); C24 alkamine - Trihydroxy (355.263/337.253/337.253/355.263); C24 alkamine - Tetrahydroxy (353.248/335.237/335.237/353.248); C24 alkamine - Pentahydroxy (351.232/333.221/333.221/351.232); C28 alcohol - 1OH (125.13/143.14); C28 alcohol - 2OH (123.12/141.13); C28 alcohol - 3OH (121.1/139.11) | The hydroxylation counts and core classes those rows define | carbon_count_massql_table.md — [TrackSheet], [Mohanty2026-AC] |

| Ref | Not observed / failed | Rules out | Source |
|---|---|---|---|
| X2 | no 86.0964/104.1070/124.9998/184.0733 (PC / lysoPC head group); no 149.0233/167.0339 (phthalate); no 85.0284 (acylcarnitine); no 44.0262 Da repeat series (PEG) (±0.005 Da, ≥1% base) | Those non-bile-acid contaminant classes | nomenclature_glossary.md |
| X3 | [M+NH4]+ 621.3680 absent from MS1 scan 3954 | Candidates with a free acidic group (free –COOH, –SO3H …) are disfavoured; blocked candidates (esters, amides, lactones) favoured. Tentative; names no substituent. Caveat: mobile-phase ammonium not stated: assumed present (most acetonitrile-containing buffers carry some) -- say so in the report; if the mobile phase in fact lacks ammonium, this absence means nothing. — "What we don't see is informative as well." | dust_region_rules.md §5 / §5a — [Specialist heuristic — Revision-2026-10-12] |
| X4 | No N4-acyl-cytosine ion [acyl-B+H]+ 486.3327 (C28H44N3O4+), nor its water losses 468.32 / 450.31 (±0.01 Da); no N-bearing ion above 230 except the precursor water losses | Disfavours a C-24 amide on the cytosine 4-NH2 (that bond would put the acyl on the released base); with C8 the cargo is bound through its oxygen. Caveat: the scan stores only its 100 most intense peaks, so an absent weak ion is weaker evidence. | neutral-loss arithmetic [Specialist heuristic] |
| X5 | [M+K]+ and [M+Na]+ neutrals (C36H55NS2, C35H51NO5; Step 0's preferred form) contain one N | Those ion forms: the base peak (C4H6N3O+) and 230.0596 (C8H12N3O3S+) need N3. [M+H]+ is the only form whose neutral contains all fragment formulas. | formula_table.py; SKILL.md Step 0 A |

| Ref | Not observed / failed | Rules out | Source |
|---|---|---|---|
| X6 | Precursor matches no free C22–C29 bile acid and no cargo row of bile_acid_deltas.tsv (class conjugated_unlisted); 622.35 ([BA + C8H11N3O3S + H]+, a non-covalent cluster) is not the precursor | A listed amino-acid / sulfate / glucuronide conjugate, and a non-covalent bile acid–drug cluster: the ion is 18.0106 lighter than the sum (covalent condensation). Free C8H11N3O3S itself elutes at 0.64 min in this run (Section 8), not 4.43 min, so the ion is not an in-source cluster of it. | dust_region.py precursor class; SKILL.md Step 3 (drug cargos) |
| X7 | No F/Cl isotope or mass-defect signature; exact mass fits C/H/N/O/S at −0.2 ppm | Emtricitabine (FTC, C8H10FN3O3S) and other halogenated nucleoside analogues as the cargo. | formula_table.py; SKILL.md Step 3 (drug cargos) |

#### Cargo evidence — what is attached to the bile acid

Every cargo candidate still in play, grouped by the question it answers. Each row cites the C/X rows above that support or argue against it; candidates already ruled out appear only in the exclusionary table. The Section 3 calls and Section 6 open questions are drawn from this table.

| Ref | Candidate | Evidence for | Evidence against | Status | Tier |
|---|---|---|---|---|---|
| **1. Cargo composition** → C8H11N3O3S (probable) | | | | | |
| A1 | **C8H11N3O3S (added as −H2O condensation)** neutral − C24H40O4 + H2O; RDBE 5 | Precursor −0.2 ppm; [X+H]+ 230.0596 (+0.7 ppm) and −H2O 212.0489; base C4H6N3O+ and C4 O,S ions are sub-formulas C3 C5 C4 C6 | 230.0596 is weak (0.86 %); no $^{34}$S peak resolved in MS1 | Supported | Probable |
| A2 | **Same cargo on a monohydroxy-monoketo core (C8H13N3O3S)** 2 H heavier cargo; RDBE 4 | Same precursor formula C3 | 230.0596 is C8H12N3O3S+, not C8H14N3O3S+; ladder 357/339/321 is the dihydroxy one C5 C7 | Disfavoured | Tentative |
| **2. Identity of the cargo** → lamivudine-type nucleoside, isomer unresolved (tentative) | | | | | |

| Ref | Candidate | Evidence for | Evidence against | Status | Tier |
|---|---|---|---|---|---|
| A3 | **Lamivudine (3TC), (2R,5S)-cytosin-1-yl-1,3-oxathiolane-2-methanol** C8H11N3O3S; one primary OH, one aromatic NH2, no acidic group | Cytosine [B+H]+ base peak with –NH3/–HNCO daughters; [X+H]+ 230.06; C4H5OS+/C4H7OS+ oxathiolane ions; a first-line HIV/HBV drug; a free C8H11N3O3S feature (m/z 230.0593) is abundant in this sample at RT 0.64 min C4 C5 C6 | No dosing record; no standard; the 3TC enantiomer and apricitabine give the same ions | Favoured | Tentative |
| A4 | **Apricitabine (oxathiolan-4-yl regioisomer)** C8H11N3O3S | Same formula and expected ions C4 C5 C6 | Investigational only, far less likely in a population sample; not separable here | Possible | Tentative |
| A5 | **Unknown endogenous/microbial C8H11N3O3S cytosine thio-nucleoside** C8H11N3O3S | Formula fits C3 | No such natural product known to the references | Unresolved | Tentative |
| **3. Bond to the bile acid** → C-24 O-ester (tentative) | | | | | |
| A6 | **C-24 ester to the cargo's 5′-CH2OH** no free acidic group in the molecule | Acyl stays on the oxathiolane when cytosine leaves (475/457); acyl + C2H2O ion 399.29; free cytosine is the base peak; no [M+NH4]+ (no free –COOH) C8 C9 C4 X3 X4 | 399.29 is one ion; ester vs amide not confirmed by a standard | Favoured | Tentative |
| A7 | **C-24 amide to the cytosine 4-NH2** no free acidic group | Same formula; no [M+NH4]+ X3 | Would release acylated cytosine (486.33, not seen) rather than free 112.05, and would not leave the acyl on the S-bearing ring X4 C8 | Disfavoured | Tentative |

#### Step-7 validation pass

Every peak was re-checked against an elemental formula (C, H, N ≤3, O, S ≤2; even-electron ion, electron mass applied): 87 of 100 fit within ±10 ppm (largest |error| 6.2 ppm; ions used as evidence above: ≤0.8 ppm). 1 peak(s) are $^{13}C$ isotope peaks of a stronger ion and are not fragments. No plausible formula: 67.1496, 112.0329, 112.0372, 112.0614, 112.0643, 112.0720, 112.0802, 113.4295, 152.4166, 222.3189, 295.4660, 604.4244 — not used as evidence. 13 ion(s) have a second formula within 5 ppm (see formula_table.tsv) and are named by both where cited. Sulfur near-miss check (§2a): not triggered on the precursor or the main S-bearing ions. Shared CHNO offset: none. $^{34}S$ A+2 on the precursor: absent — no peak at the 34S position; weighs against S if the A+2 region was resolved.

Odd/even parity check (conjugate_fragments.md): 12 even-mass fragment(s) (94.0401, 112.0175, 112.0397, 112.0505, 170.0387, 212.0489, 230.0596, 500.4030, 568.3220, 586.3322, 604.2760, 604.3414) — each carries an odd number of N in its best formula. 3 odd-mass ion(s) carry an even number of N (e.g. N2). The nitrogen rule holds for every assigned ion.

#### 5. Automated Tree Outcome

The automated MassQL tree ran to a terminal bin without stalling; the bin reached and its published FDR (where applicable) are reported in Section 3.

#### 6. Unresolved Aspects

1. **Hydroxyl positions — not determined.** No classification-tree path was satisfied: the core pair (weakest ion 1.4 % of base) sits below the tree's floors (stage1 ≥ 2 %, Dihydroxy_stage2 ≥ 5 %). The dust region has 1 ring ion (7 required), so the fingerprint/dust-ratio fallback has nothing to read; the Di-3,6 and Di-3,12β query ions (0.4–0.6 %) are not used. Candidates include DCA, CDCA, UDCA, HDCA and other 2-OH isomers. Energy went to the cargo (base peak 112). HCD 41.7 is below NCE 44–45; lower energy alone does not lower a tier (stereochemistry_and_caveats.md §4).
2. **α/β and 5α/5β — not resolvable from MS/MS alone.** No ratio rule fires; the run is not in the RT-order reference (run_not_in_library), so no RT-order corroboration.
3. **Core call rests on a relaxed-gate pair.** 339.2685 (2.1 %) and 321.2576 (1.4 %) pass the adapted 1 % floor but not the published 5 % ([Adaptation-2026-10-18]); the published FDR/FNR do not apply, and the core call and overall tier are capped at tentative. Other claims are graded on their own evidence.
4. **Ion form chosen from fragments, not MS1.** The MS1 could not separate [M+H]+, [M+K]+, [M+H-H2O]+; the Step 0 screen (C/H/O + listed cargos) preferred [M+K]+. [M+H]+ was set because only its neutral contains the N3/S fragments (X5) and GNPS2 calls the feature [M+H]1+. The $^{34}S$ check is inconclusive: the A+2 peak (606.3459, 8.4 %) sits between the $^{34}S$ (606.3372) and $^{13}C_2$ (606.3481) positions, as expected if the two are not resolved.
5. **Cargo identity: lamivudine vs its isomers.** See A3 – A5 . 3TC, its enantiomer and apricitabine give the same ions here. Settled by dosing records for the participant and co-injection with a synthetic 3TC 5′-O-bile-acyl ester (which would also fix the OH positions).
6. **Ester vs amide — tentative.** See A6 – A7 : the fragments favour the 5′-O-ester. A synthetic standard, or MS3 of 457.28, would settle it. The scan keeps only its 100 most intense peaks, so absent ions (X4) are weaker evidence.
7. **Ammonium result assumes ammonium in the mobile phase.** No [M+NH4]+ at 621.3680 in MS1 scan 3954 (X3): consistent with a blocked carboxylate (ester or amide); not stated, so assumed present. If the mobile phase lacked ammonium, this absence means nothing.
8. **Origin: drug metabolite, microbial product or artefact?** Bile acid–drug esters of this kind are not described in the reference files. Whether it forms in the host, in the gut microbiota or ex vivo is open (Section 7).
9. **GNPS2 network and library context not used.** The feature (15549: m/z 604.3415, RT 4.4255 min, [M+H]1+, max area 6.5e6 in this file) was read from the everything_bagel task, but the network graphml and its library hits could not be downloaded here, so no network analog table (Section 11) or library reconciliation was done.

#### 7. Species / Taxonomic Context

Prior, not evidence. The file name (AGINCOURT; other samples in the job are NAIROBI and DIMAMO) suggests a human multi-site African cohort, which is a setting where lamivudine is a standard part of first-line HIV treatment; this changes no tier.

| Part | Finding | Source |
|---|---|---|
| **Host prior** | Probably human (inferred from the file name only: AGINCOURT, with NAIROBI and DIMAMO samples in the same GNPS2 job — sites of a multi-centre African population study). Dihydroxy C24 bile acids (DCA, CDCA, UDCA) are the expected human pool. Not recorded in the mzML. | file names; GNPS2 feature table |
| **Expected producer** | No documented producer: a bile acid 24-O-ester of a nucleoside drug is not in the reference files. The bile-acid part is host/microbe-typical; the cargo is a xenobiotic. Bacterial acyl-transfer by bile salt hydrolases forms amides with amines; ester formation with a drug alcohol is undocumented. | source_and_diet_context.md §3; conjugate_fragments.md |
| **Plausible route** | Drug-derived, not diet: diet not provided and not relevant to this cargo; the route would be lamivudine dosing (oral, HIV/HBV therapy). A free C8H11N3O3S [M+H]+ feature (230.0593, RT 0.64 min, apex 3.4e7) is present in this sample, consistent with exposure. | source_and_diet_context.md §3 |

Follow-ups that would test the route:

- Check the participant's antiretroviral records (3TC-containing regimen?).
- Look for the same 604.34 feature only in samples that also contain the free 230.059 feature, across the GNPS2 job.
- Synthesize or buy 3TC and esterify with DCA/CDCA/UDCA to compare RT and MS2.
- Search the job for other bile-acid acyl esters of 3TC (mono-, tri-hydroxy cores: 588.35, 620.34) and of other ART drugs.

#### 8. Chromatographic Context (Supporting, Not Evidence)

Extracted-ion chromatograms (±5 ppm) over the whole 10.0-min run, for M = 603.33414 ([M+H]+, set by the annotator (--ion-form): MS1 does not separate [M+H]+/[M+K]+; only [M+H]+ gives a neutral (C32H49N3O6S, -0.2 ppm) whose N3/S-bearing fragments (230.0596 C8H12N3O3S+, 212.0489, 112.0505, 475.2876/457.2773 C28H4xO S+) all fit as sub-formulas; GNPS2 everything_bagel also calls feature 15549 [M+H]1+):

| Trace | Chromatographic peaks (RT min, apex intensity) |
|---|---|
| 604.3414 [M+H]+ (precursor) | 4.26 (3.0e5), 4.42 (6.5e6) ← this scan |
| 621.3680 [M+NH4]+ | 3.37 (2.5e4), 3.79 (3.2e4), 4.54 (2.6e4), 4.73 (4.1e4), 5.09 (2.1e4), 5.94 (1.8e5), 7.47 (2.7e4), 8.23 (3.9e4) |
| 626.3234 [M+Na]+ | 3.11 (4.7e4), 3.33 (4.6e4), 3.47 (6.1e4), 3.75 (5.1e4), 3.88 (7.8e4), 4.03 (4.5e4), 4.26 (1.2e5), 4.44 (6.9e4) ← this scan |
| 642.2973 [M+K]+ | 2.50 (3.6e4) |
| 586.3309 [M+H-H2O]+ | 2.89 (7.0e4), 3.01 (2.8e4), 4.48 (1.9e4) ← this scan |
| 230.0594 C8H12N3O3S+ (free cargo, GNPS2 XIC) | 0.64 (3.4e7) — not co-eluting with the precursor (4.43 min) |

Supporting orientation only: retention time alone identifies nothing (stereochemistry_and_caveats.md). 1 other MS2 scan(s) were acquired on this precursor in the run. GNPS2 everything_bagel feature 15549 (m/z 604.3415, RT 4.4255 min, [M+H]1+, max area

6.5e6, top sample this file) matches this scan; the free-cargo trace comes from the same job's XIC service.

#### 9. Diagnostic Decision Tree

Each node's accent color is the color of the corresponding ion family highlighted in the Section 2 spectrum. The numbered chips (C = confirmatory, X = exclusionary, A = cargo evidence) are the Ref column of the Section 4 evidence tables, so every node traces back to its ions, mass errors and sources.

Dehydrated-core pair · Conjugate / cargo ions · Grouping / precursor arithmetic · Ruled out (no diagnostic ion family)

✓ supports the annotation · ⊘ alternative ruled out (dashed box) · – context only

**MS2 scan 3956**
precursor 604.3414 $[M+H]^+$ · HCD 41.7 · base peak 112.0505

**Core identity & conjugate**
carbon count, OH count, cargo, bond

- ✓ **C24 dihydroxy pair (relaxed gate)**
  357/339/321 at 1.4–3.3 %
  C1 C7
- ✓ **Exact mass $C_{32}H_{49}N_3O_6S$**
  –0.2 ppm; two water losses
  C3 C10
- ✓ **Cytosine base peak**
  112.0505 → 95.0241, 69.0447
  C4
- ✓ **Cargo $[X+H]^+$ $C_8H_{12}N_3O_3S^+$**
  230.0596, 212.0489; oxathiolane 101/103
  C5 C6 A1 A3
- ✓ **Acyl stays on S-ring**
  475/457 after cytosine loss ⇒ 5′-O-ester
  C8 C9 A6
- ✓ **No $[M+NH_4]^+$**
  no free –COOH (mobile phase assumed to carry $NH_4^+$)
  X3

**Hydroxylation position**
tree and dust

- ✗ **Dust region**
  1 ring ion of 7 needed → positions unresolved
  C2

**Alternatives ruled out**

- ⊘ **K/Na ion forms**
  one N; fragments need N3
  X5
- ⊘ **N4-amide on cytosine**
  no 486.33 acyl-cytosine (disfavoured)
  X4 A7
- ⊘ **Listed conjugates / cluster**
  no table cargo fits; $-H_2O$ ⇒ covalent
  X6
- ⊘ **Emtricitabine / halogenated**
  no F in formula
  X7
- ⊘ **Contaminant classes**
  no PC/phthalate/PEG/carnitine markers
  X2 X1

↓

**Dihydroxy C24 bile acid 24-O-ester of a lamivudine-type nucleoside (Tentative)**

formula consistent with a 3TC 5′-O-bile-acyl ester; OH positions, stereo and isomer unresolved (Section 6)

#### 10. Proposed Final Structures

One row per structure the reasoning in Sections 3–9 still supports, drawn with RDKit with the ABCD rings in the standard bile-acid orientation. A partial structure draws only what the evidence claims: wavy bonds mark stereocentres that are not resolved, substituents that are counted but not placed are listed under the drawing and their possible rings shaded, and a highlighted unit is weaker than the rest (the Notes say why). The Tier column is copied from the reasoning. Names are standardized with the metabolite-annotation-cleanup rules (PubChem's title when the exact, stereo-defined structure's record lists the proposed name; otherwise the cleaned label); standardization never changes a tier.

| Structure | Name | Tier | Notes |
|---|---|---|---|
| 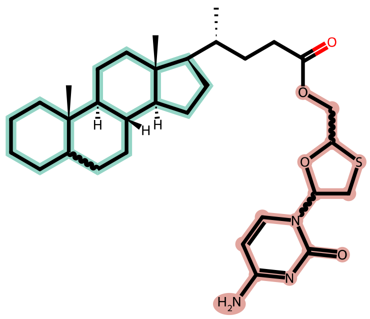<br>+ 2 OH (positions not resolved) | **lamivudine 5′-O-(dihydroxy-cholan-24-oyl) ester (partial)**<br>partial structure<br>$C_{32}H_{49}N_3O_6S$ · 603.33421 Da<br>drawn skeleton: $C_{32}H_{49}N_3O_4S$ | Tentative | Partial structure: the two OH are counted, not placed.<br>Wavy bonds: stereochemistry not resolved at C5, 2 other<br>2 × OH on rings A–D (shaded); positions not resolved<br>Highlighted unit: cargo fixed to $C_8H_{11}N_3O_3S$ and a cytosine-oxathiolane by fragments; identity as 3TC not confirmed<br>Not resolved:<br>• Positions of the two OH (dust region fails; no tree path)<br>• α/β of each OH and 5α/5β<br>• Oxathiolane stereocentres: 3TC (2R,5S) vs its enantiomer; apricitabine (4-yl regioisomer) also fits<br>• Ester to the 5′-CH2OH vs amide to the cytosine NH2 (ester favoured) — a synthetic standard would settle it<br>Assumed, not shown by this spectrum:<br>• Backbone 8β,9α,10β,13β,14α,17β and 20R of a normal C24 bile acid<br>Name: partial structure (substituent positions not resolved): no PubChem lookup; cleaned label shown |

*Generated by the "bile-acid-msms-annotation-v57" skill (deterministic MassQL pre-pass, pure evaluator) plus literature-grounded reasoning in the referenced skill files. Source: AG078_AGINCOURT_6_B6.mzML, scan 3956. Run time: 6 min 37 s, start to finish (started 2026-11-04 19:39:08 UTC, report rendered 2026-11-04 19:45:45 UTC).*
